# A Reptilian Endogenous Foamy Virus Sheds Light on the Early Evolution of Retroviruses

**DOI:** 10.1101/426585

**Authors:** Xiaoman Wei, Yicong Chen, Guangqian Duan, Edward C. Holmes, Jie Cui

## Abstract

Endogenous retroviruses (ERVs) can be thought of as host genomic fossils of ancient viruses. Foamy viruses, including those that form endogenous copies, provide strong evidence for virus-host co-divergence across the vertebrate phylogeny. Endogenous foamy viruses (EFV) have previously been discovered in mammals, amphibians and fish. Here we report a novel endogenous foamy virus, named SpuEFV, in genome of the tuatara (*Sphenodon punctatus*), an endangered reptile species endemic to New Zealand. Phylogenetic analyses revealed that SpuEFV has likely co-diverged with its host over a period of many millions of years. The discovery of SpuEFV fills a major gap in the fossil record of foamy viruses and provides important insights into the early evolution of retroviruses.

## Introduction

Retroviruses (family *Retroviridae*) are viruses of major medical significance as some are associated with severe infectious disease or are oncogenic (Hayward, et al. 2015; Aiewsakun and Katzourakis 2017; Xu, et al. 2018). Retroviruses are also of note because of their ability to integrate into the host germ-line, generating endogenous retroviruses (ERVs) that then exhibit Mendelian inheritance (Stoye 2012; Johnson 2015). ERVs are widely distributed in vertebrates (Hayward, et al. 2013; Cui, et al. 2014; Hayward, et al. 2015; Xu, et al. 2018) and constitute important molecular “fossils” for the study of retrovirus evolution. ERVs related to all seven major retroviral genera have been described, although some of the more complex retroviruses, such as lenti-, delta- and foamy viruses, rarely appear as endogenous copies.

As well as being agents of disease, foamy viruses are of importance because of their long-term virus-host co-divergence (Switzer, et al. 2005). Endogenous foamy viruses (EFVs), first discovered in sloths (class Mammalia) (Katzourakis, et al. 2009) also exhibit co-divergence pattern with their hosts; and they have also been reported in primates and the Cape golden mole (Han and Worobey 2012b, 2014). The discovery of a EFV in the coelacanth genome indicated that foamy viruses could have an ancient evolutionary history (Han and Worobey 2012a), likely co-diverging with their vertebrate hosts over time-scales of hundreds of million years (Aiewsakun and Katzourakis 2017). Although EFVs or foamy-like elements have been reported in fish, amphibians and mammals, they have currently not been reported in genomes of two other major classes of vertebrates - reptiles and birds (Tristem, et al. 1995; Hayward, et al. 2015; Xu, et al. 2018).

## Materials and Methods

### Genomic mining and consensus genome construction

To identify foamy viruses in reptiles, the TBLASTN program (Altschul, et al. 1990) was used to screen relevant taxa from 28 reptile genomes (Supplementary Table S1) and 130 bird genomes (Supplementary Table S2) (as of October 2018) downloaded from GenBank (www.ncbi.nlm.nih.gov/genbank). In each case, the amino acid sequences of the Pol genes of representative EFVs (endogenous foamy viruses), endogenous foamy-like viruses, and exogenous foamy viruses were chosen as queries. As filters to identify significant and meaningful hits, we chose sequences with more than 30% amino acid identity over a 30% region, with an e-value set to 0.00001. Genomes that contained only single hits for EFVs were excluded as likely false-positives. We extended viral flanking sequences of the hits to identify the 5’- and 3’-LTRs using LTR finder (Xu and Wang 2007) and LTR harvest (Ellinghaus, et al. 2008). Sequences highly similar to foamy virus proteins found in tuatara were aligned to generate a SpuEFV consensus genome (Supplementary Table S5). Conserved domains were identified using CD-Search service in NCBI (Marchler-Bauer and Bryant 2004).

### Phylogenetic analysis

To determine the evolutionary relationship of EFVs and retroviruses, the Pol and Env protein sequences were aligned in MAFFT 7.222 (Katoh and Standley 2013) and confirmed manually in MEGA7 (Kumar, et al. 2016). The phylogenetic relationships among these sequences were then determined using the maximum-likelihood (ML) method in PhyML 3.1 (Guindon, et al. 2010), incorporating 100 bootstrap replicates to determine node robustness. The best-fit models of amino acid substitution were determined by ProtTest 3.4.2 (Abascal, et al. 2005): RtREV+Γ+I for Pol, LG+Γ+I+F for concatenated gag, pol and env. All alignments used in the phylogenetic analyses can be found in Data set S1-S2.

## Results and Discussion

### Discovery of foamy viral elements in reptile genomes

To search for potential foamy (-like) viral elements in reptiles and birds, we collated 28 reptilian genomes (Supplementary Table S1) and 130 bird genomes (Supplementary Table S2) and performed *in silico* TBLASTN with full-length Pol protein sequences of various foamy viruses, including EFVs, as screening probes (Supplementary Table S3). We only considered viral hits within long genomic scaffold (>20 kilobases in length) to be *bona fide* ERVs. This genomic mining identified 117 ERV hits in tuatara (*Sphenodon punctatus*) and none in bird genomes. Hence, a total of 117 ERV hits in the tuatara genome were extracted and subjected to evolutionary analysis (Supplementary Table S4). We named this new ERV as SpuEFV (***S**phenodon **pu**nctatus* **e**ndogenous **f**oamy **v**irus).

### Genomic organization

We extracted all significant SpuEFV viral elements and constructed a consensus genomic sequence of SpuEFV (Supplementary Fig. S1, Table S5). The consensus genome harbored a pairwise long terminal repeats (LTRs) and exhibits a typical spuma virus structure, encoding three mainly open reading frames (ORF) – *gag*, *pol* and *env* – and one putative additional accessory genes, ORF1 (Fig. 1). Interestingly, this accessory ORF 1 exhibit no sequence similarity to known foamy accessory genes. Notably, by searching the Conserved Domains Database (www.ncbi.nlm.nih.gov/Structure/cdd), we identified three typical foamy conserved domain for both consensus and one full-length original SpuEFV (Accession no. QEPC01003194.1): (i) Spuma virus Gag domain (pfam03276) (Winkler, et al. 1997), (ii) Spuma aspartic protease (A9) domain (pfam03539) which exists in all mammalian foamy virus pol protein (Aiewsakun and Katzourakis 2017), and (iii) foamy virus envelope protein domain (pfam03408) (Han and Worobey 2012a) (Supplementary Fig. S2, Fig. S3), confirming that SpuEFV is indeed of foamy virus origin.

**Figure 1.**
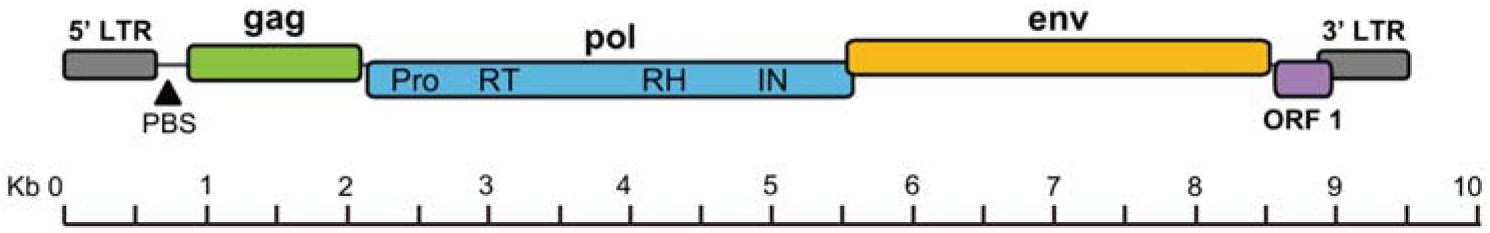
Genomic organizations of SpuEFV. LTR, long-terminal repeat; PBS, primer-binding site; Pro, aspartic protease; RT, reverse transcriptase; RH, ribonuclease H; IN, integrase.

### Phylogenetic analysis

The Pol (490 amino acids) of SpuEFVs were used for phylogenetic analysis. Our maximum likelihood (ML) phylogenetic trees revealed that the EFVs present in that tuatara genome formed a close monophyletic group within the foamy clade, indicative of a single origin, and with high bootstrap support (Fig. 2). The divergent phylogenetic position of SpuEFV is compatible with virus-host co-divergence for the entire history of the vertebrates. However, it is possible that this pattern will change with a larger sampling of taxa such that the EFV phylogeny expands. Failure to detect any SpuEFV related elements in the remaining reptilian genome screening suggests that the virus was not vertically transmitted among reptiles, although this will clearly need to be reassessed with a larger sample size.

Previous studies provided strong evidence for the co-divergence of foamy viruses and their vertebrate hosts over extended time-periods (Katzourakis, et al. 2009). That the reptilian SpuEFV newly described here seemingly follows the same pattern (Fig. 3) thereby implies that it could diverge from the other mammalian foamy viruses with its tuatara host more than 320 million years ago (http://www.timetree.org/). As such, the discovery of SpuEFV fills a major gap in our knowledge of the evolutionary history of the foamy viruses and provides important insights into the early evolution of retroviruses.

**Figure 2.**
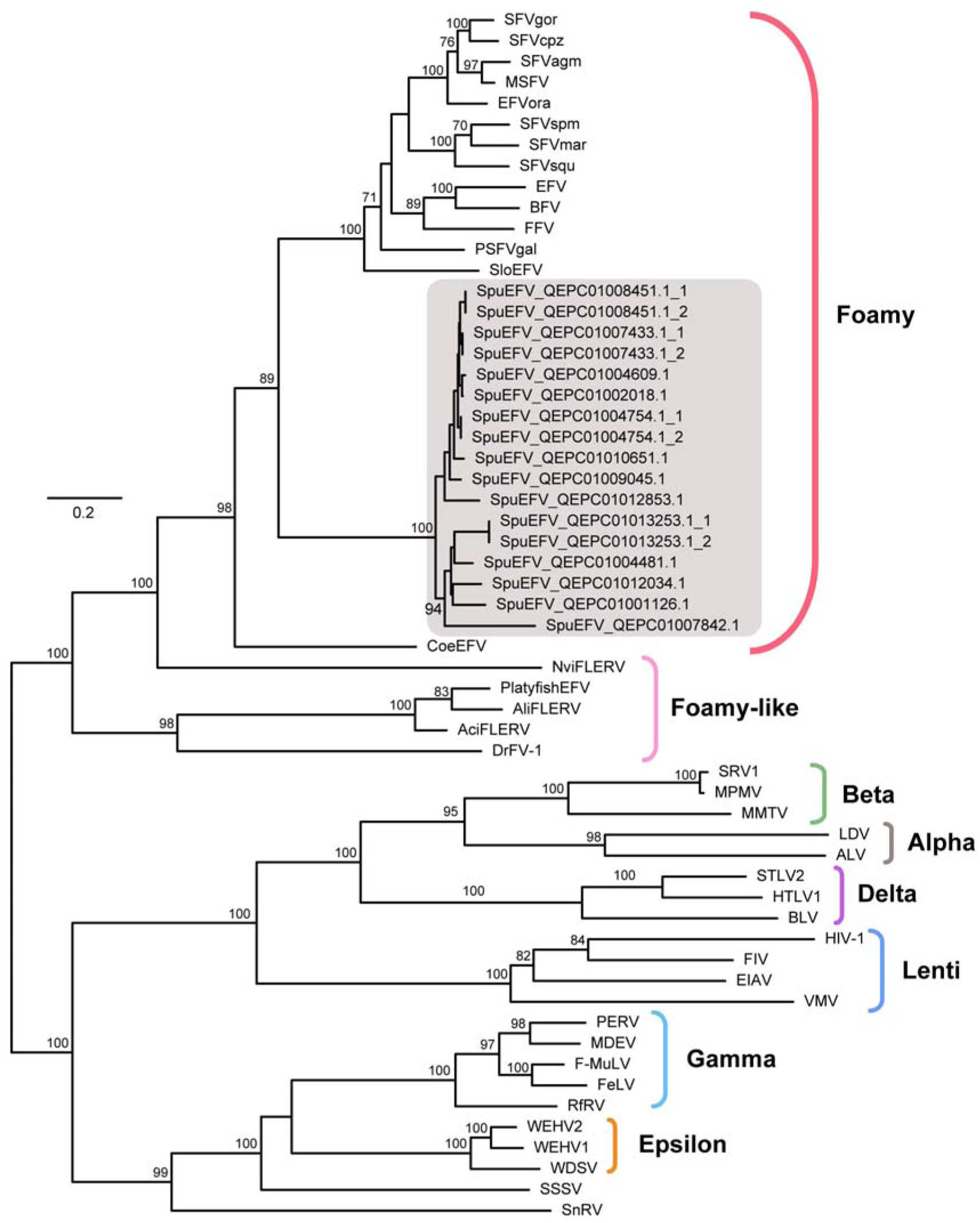
Phylogenetic tree of retroviruses, including SpuEFVs, inferred using amino acid sequences of the Pol gene (490aa). The tree is midpoint rooted for clarity only. The newly identified SpuEFVs are labelled using a grey-shaded box with their accession numbers (different pol sequences in same contig are numbered in the suffix). The scale bar indicates the number of amino acid changes per site. Bootstrap values <70% are not shown. The alignment of pol amino acid sequences is provided in Data set S1.

**Figure 3.**
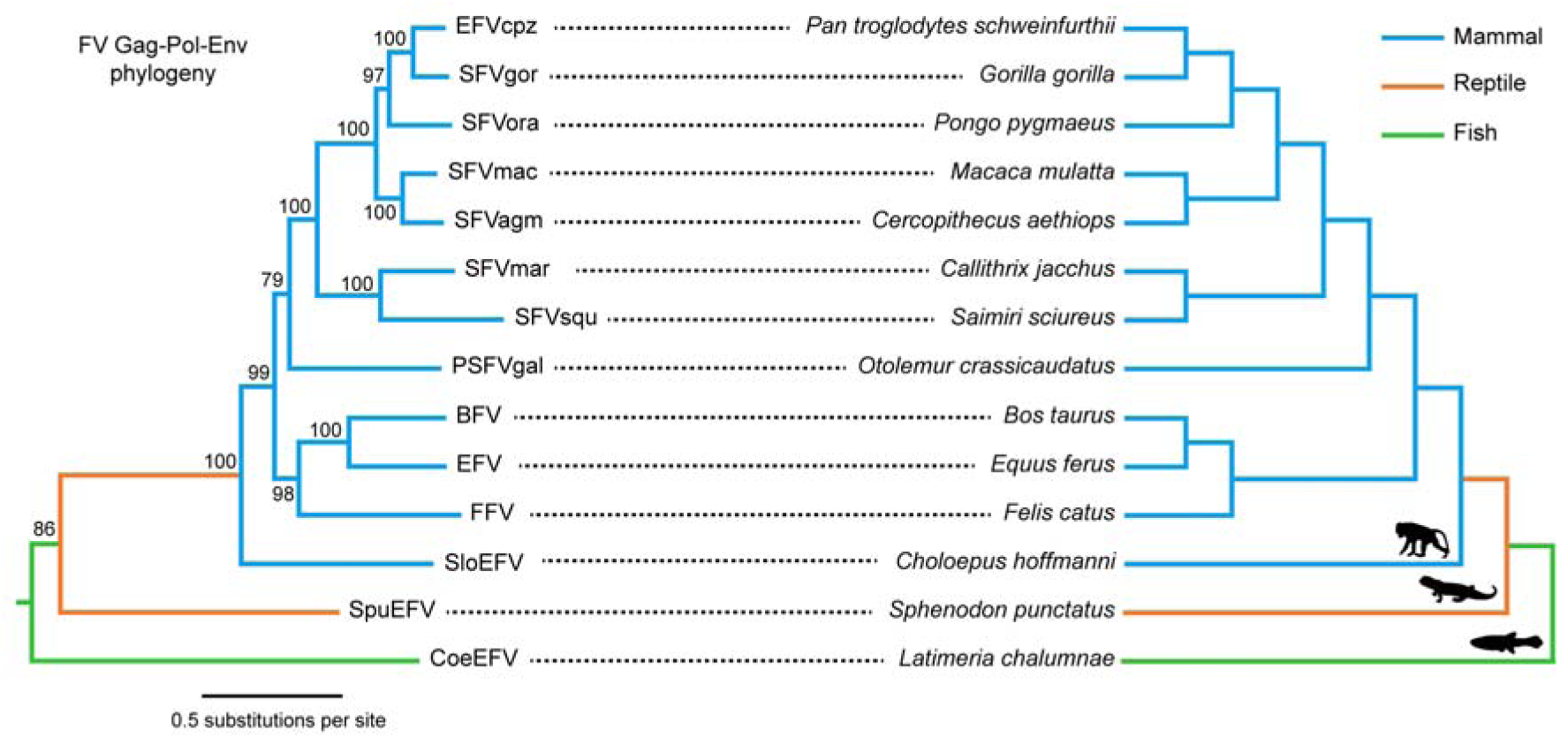
A simplified evolutionary relationship between foamy viruses (left) and their vertebrate hosts (right). The scale bar on the virus phylogeny indicates number of amino acid changes per site with bootstrap support values provided at each node. The alignment of FV gag-pol-env amino acid sequences is provided in Data set S2.

## Supporting information

## Acknowledgments

J.C. is supported by National Natural Science Foundation of China (31671324) and CAS Pioneer Hundred Talents Program. ECH is supported by an ARC Australian Laureate Fellowship (FL170100022).

## Data availability

All data needed to evaluate the conclusions in the paper are present in the paper and/or the Supplementary Materials. Additional data related to this paper may be requested from the authors.

## Conflict of interest

None declare

