## Supplementary material for "A Reptilian Endogenous Foamy Virus Sheds Light on the Early Evolution of Retroviruses"

- **Fig. S1.** Detailed descriptions of consensus SpuEFV genome.
- **Fig. S2.** Conserved domain alignment of the consensus SpuEFV proteins and foamy virus proteins in CDD (Conserved Domain Database).
- **Fig. S3.** Conserved domain alignment of the original SpuEFV proteins and foamy virus proteins in CDD (Conserved Domain Database).
- **Table S1.** The information of 28 reptile genomes used for data mining.
- **Table S2.** The information of 130 bird genomes used for data mining.
- **Table S3.** The information of representative retroviruses used for phylogenetic analysis.
- **Table S4.** The matching contigs identified in *Sphenodon punctatus* genome.
- **Table S5.** The important SpuEFV sequences used for constructing consensus genome.
- **Data set S1.** The alignments used to build the phylogenetic trees of Pol represented in Fig. 2.
- **Data set S2.** The alignments used to build the phylogenetic trees of endogenous/exogenous foamy virus in Fig. 3.

### SpuEFV consensus genome

| 5′LTR start

1 ACTCAGGCAGGGCAGGAAAAGGCCTTGTTTGTTTGGAGTTTACTCATTATTAATCCTATCTTATACGATAGGAACCTGCTTAGTATGTAT

91 CAACAGACGTCTTACTATGTATTTTGAGAAAAAACCCTCCCTGTCAGCAGTCGTGACCTGACGTCATTTTTAGCTAACCTCCAAGCATCT

181 TTCATTCTATAAGAAGAGAGAGGAAAGATGGTTTTTTCAGAAAGCTTCCTTTCCTGAAGCTTTTCTCATCTTCCATCTGGCAGAATCGGG

271 TAAAGGACTGGGTAATATGTTATATCCAATATGATGCACTGAGTCAACCATAGCAATAATACTTAGAGCATATGTTATATCTTATAGATG

361 TTAAGATATTTGAATTTGTAACAGAGCTGTTCTTTGAACAAGACATGTCTTTCTAAGTCATGTACTTTTAACTGGCACTGTTTCTTTTGT

451 ACTATGTCTTTATTTGAACTTAACTATTGCTTCATATTTTTATAATATTCTTAAAGTCTCATAGCCTCATTTATATAAATGAATTGGTCT

541 TTTCTATAATTTGCTGAGCTGGCTGGCTGATTAGCAAAGTGTAAACTCGAATTCCACATATTGGAAAAGAGCGCCCTATATCCAGACCAG

5′LTR end | PBS

631 ATCATGTCAGGCCTGGACCACTAAGCGCAAATTCCTTTTGGAGAAACTGAGGCACAACGTGACAAT**TGGCACCCATTCGTGGGGC**TCGAG

721 ACATAAGAATATATAATTAAGTACACTGGGGTGTTCACCAGACCGTTAACTACTTTTGTCCTCTTGATATATTCAAGTTTGTTAAGCTTA

| Gag start

M A I Q L N Q V P Y A L W V A N L Q N

811 ACTAAAAGAAAAGGATTTTGTGCATTTTATATCATGGCTATTCAATTAAACCAAGTGCCTTATGCACTATGGGTTGCTAATCTGCAAAAT

V T V R D G D D Y C L E I R N G E W G I G H R F L I V S F E

901 GTAACAGTCAGAGATGGAGATGATTATTGTTTAGAAATAAGAAATGGAGAATGGGGAATTGGACACAGATTTCTAATTGTTTCTTTTGAA

A N D A G V I V S M R L R D V V F N P L T M P V L P I N R Q

991 GCAAATGATGCAGGTGTAATAGTTTCTATGAGACTAAGAGATGTAGTTTTTAATCCATTGACAATGCCTGTGTTACCTATCAATAGGCAA

D L N L M A G I V V E I P K N I R R H G P I S I S N A D Y I

1081 GATTTGAACCTAATGGCAGGGATAGTAGTTGAAATTCCCAAAAATATAAGGAGGCATGGGCCTATTTCTATAAGTAATGCTGATTATATT

S G R Y S S Y H R G I A W L Q C S P V G T G V H S I K K R I

1171 TCTGGAAGATACTCCAGCTACCACCGAGGAATTGCGTGGCTTCAATGCAGCCCAGTGGGAACAGGAGTACACTCTATTAAGAAGAGAATA

F K S E E F A K S N V G R Q P I V P G V H P A E I S V L N M

1261 TTTAAATCTGAGGAATTTGCAAAAAGTAATGTTGGAAGACAACCTATTGTGCCAGGAGTACACCCAGCTGAAATTTCTGTCTTAAATATG

A N I R A V T G P T P K D F K E I P M W F E S H L S A L E A

1351 GCAAATATAAGGGCTGTAACTGGGCCCACTCCAAAGGATTTTAAAGAAATTCCTATGTGGTTTGAATCCCATTTATCTGCTTTAGAAGCT

V T S T A S P L Q K M R L C N S L V P A S A S L I E Q E C N

1441 GTCACTTCCACAGCATCACCATTGCAGAAGATGAGGTTATGCAATTCTCTAGTCCCAGCATCTGCTTCTTTGATAGAGCAAGAATGTAAT

N W E S V L A N L Y V K T H G Q V G I A D L N E I L R K I T

1531 AATTGGGAAAGTGTATTGGCTAATTTATATGTAAAAACTCATGGCCAAGTGGGCATAGCTGACTTAAATGAAATTCTAAGAAAGATAACT

Q E Q G I V R A Y G V G M K F L S N H D L I W G I L K A L C

1621 CAAGAGCAAGGCATTGTAAGGGCCTATGGGGTTGGAATGAAATTTCTCTCTAATCATGATTTGATATGGGGAATCCTTAAAGCCCTCTGT

K G D V L K A A I Q S K L D L L I T E Q E K I R S F P K I V

1711 AAAGGTGATGTATTGAAAGCTGCCATTCAATCTAAATTAGATTTGCTGATTACTGAGCAGGAAAAAATAAGGTCCTTCCCAAAAATTGTC

Q D I Y K T L G R D Y L G N N P N R K N L E Q E G S K S K K

1801 CAGGATATTTATAAGACTCTGGGGAGAGATTATTTGGGAAATAATCCAAACAGAAAGAATCTTGAACAGGAAGGTTCTAAATCTAAGAAA

S P S N I N S R K K I L P Q N S Q P N Y R G K G G K I Q G S

1891 TCTCCCTCAAATATTAATTCTAGAAAAAAAATTCTTCCCCAGAATTCTCAGCCAAATTATAGAGGCAAAGGAGGGAAGATCCAAGGCTCA

N R K Q F Q A Q K D Q D S E N P E G A A T Y D L R K G S Q F

1981 AATAGAAAGCAGTTCCAAGCTCAGAAAGACCAGGATTCTGAAAATCCAGAAGGTGCAGCTACATATGACTTAAGAAAAGGATCACAGTTT

**Gag end |**

P H R D F D K S K Q D F N S R G G F R G N K *

2071 CCCCATAGAGACTTTGATAAATCAAAGCAAGATTTTAATTCCAGAGGTGGATTCAGAGGAAATAAATGAGGTTTTAATAAAAGAGGAAAT

**| Pol start**

M A E N R Q

2161 TATCAGACCTCTAAAGACTATGATCAACTGACATCCTCACAAAAAGCTTCAGAAACTCCCTCTGTGACACAGGATGGCAGAGAACAGACA

A Q V S L E F P M I S V E I A D K A F T A D R H R G K S I L

2251 AGCTCAGGTCAGCCTTGAATTTCCAATGATCTCTGTAGAAATAGCAGATAAGGCTTTCACAGCTGATAGACACAGGGGCAAATCAATCTT

Y S I K M F A T C I Y N S K V N V Y V E S Y F L E E K V V P

2341 GTATTCAATCAAAATGTTTGCCACATGTATATACAACTCAAAGGTCAATGTATACGTTGAGTCTTATTTTTTAGAAGAGAAAGTAGTACC

**Aspartic protease**

L V K L P V Q I A G Q C L E I E F A I C D F T K H D V V I A

2431 TCTTGTAAAATTACCTGTTCAGATAGCAGGACAATGTTTGGAAATTGAATTTGCAATATGTGATTTTACAAAACATGATGTTGTCATAGC

H E R V K D L F P I G S I N I I G T Q D N R G E Q I K E Q I

2521 TCATGAAAGGGTTAAGGATTTGTTTCCAATAGGTTCAATAAACATAATTGGAACTCAAGATAATAGGGGTGAGCAAATCAAAGAGCAAAT

A S A D C A K N E K T K L R D I L Y S L K P Y F Q Q F D N Q

2611 TGCATCAGCTGACTGTGCAAAAAATGAAAAAACAAAATTGAGAGATATTTTATACTCTCTGAAACCATATTTTCAGCAATTTGATAATCA

Q K Q Y P I N K A A I

I G H R K I K P H D L S V K T Q P K P Q K Q Y P I N K A A I

2701 AATTGGCCACAGGAAAATAAAGCCTCATGATTTATCTGTCAAAACACAGCCCAAACCGCAAAAGCAATATCCTATTAACAAAGCTGCAAT

N D I Q K V I N D L I A Q G A L I R Q Y S S M N T P V Y P V

2791 TAATGATATTCAAAAAGTAATTAATGATTTAATTGCACAAGGAGCATTGATTAGACAATATAGCTCCATGAATACTCCAGTATATCCAGT

P K P N G K W H M V L D Y R A L N R V S P S F N V Q N L H V

2881 GCCTAAACCTAATGGGAAATGGCATATGGTATTAGACTACCGCGCCCTAAACAGGGTTTCACCCTCTTTTAATGTGCAAAACCTACACGT

W H A R K F G K A Q I Q N Y T G F I Q W I L G S S Y R E Q D

2971 CTGGCATGCTAGGAAATTTGGAAAGGCACAAATACAAAACTACACTGGATTTATCCAATGGATTTTGGGCTCATCCTATAGAGAACAAGA

**Reverse transcriptase domain**

R Y L T A F S W Q G T Q Y C W T R L P Q G Y L N S P A L F S

3061 CAGATATTTGACTGCGTTTAGTTGGCAGGGAACACAGTACTGTTGGACACGCCTTCCACAAGGCTATTTAAATAGTCCAGCGTTGTTTTC

A D V I Q L L K N I P G V H S Y M D D I Y F T N E D L D Q H

3151 AGCTGACGTAATTCAGCTTTTGAAAAATATTCCAGGTGTACATTCATATATGGATGACATCTACTTTACAAATGAAGATTTAGATCAACA

L A T L K Q I V T V L G E A G Y I I N L K K S Q I C R S K V

3241 TTTAGCTACATTAAAGCAAATTGTCACTGTTTTGGGAGAAGCAGGATATATTATCAATCTGAAAAAATCACAGATTTGTAGAAGCAAAGT

K F L G F L L T D S G R G L T Q E F K E K L L T L Q P P K T

3331 AAAGTTCCTTGGTTTCTTATTAACAGACTCTGGCAGAGGATTAACTCAGGAATTCAAAGAAAAACTGTTGACATTGCAACCTCCTAAAAC

L K E L Q S V L G F L N V A R I L V P D Y A Q R T K P L Y N

3421 TTTAAAAGAATTGCAGTCAGTTTTAGGTTTTTTAAATGTTGCAAGAATATTGGTTCCTGACTATGCTCAGAGAACCAAACCCCTTTATAA

L I P L A S K G N F W T L E A Q Q T L D N L I T L I N Q A V

3511 TCTAATTCCATTAGCAAGCAAAGGGAATTTTTGGACCTTGGAAGCTCAGCAAACATTGGATAATTTGATAACCTTAATTAACCAAGCTGT

E L N T R N S T V S L E I L V G A T Q K G G F A S Y F N Y G

3601 TGAATTAAACACAAGAAACAGCACAGTTTCCCTAGAAATTCTAGTTGGAGCCACACAAAAAGGAGGATTTGCTTCATATTTCAATTATGG

E S K P L Q Y I S Y V F S N A E Q K F L P I E R I L C M C N

3691 AGAATCTAAACCTCTGCAGTACATATCATATGTCTTCTCAAATGCAGAACAAAAGTTTTTGCCTATTGAAAGAATCTTATGTATGTGTAA

L A I L K G K D L A Q G Q K M I V K T P I A S L R Q V K K G

3781 TCTTGCAATTTTAAAAGGTAAAGATCTTGCACAGGGCCAAAAGATGATAGTAAAAACTCCTATTGCCTCCTTAAGGCAGGTTAAAAAAGG

S I P N A K A L H S R W V Q W M S H F E N P Q I E F E Y V E

3871 ATCAATACCCAATGCAAAAGCTCTTCATAGCAGATGGGTACAATGGATGTCACATTTTGAAAATCCACAAATAGAATTTGAATATGTGGA

P S N D L E N L P A F T I E P V S S K N H T P D K P L H E Y

3961 ACCAAGCAATGATTTAGAAAATTTGCCAGCTTTTACTATTGAACCTGTATCTTCTAAAAATCATACCCCAGATAAACCTCTCCATGAATA

Q K V I Y T D G S A M S C K Q G K H M W K A G C A V V I G T

4051 TCAAAAAGTTATCTATACTGATGGTTCAGCCATGTCATGTAAGCAAGGCAAACACATGTGGAAAGCAGGTTGTGCTGTTGTAATTGGGAC

F N D K G E Y H M S D S I Q M P L G N N T A Q Y A E L M A V
4141 ATTTAATGATAAAGGGGAATATCATATGTCTGACAGTATTCAAATGCCTTTAGGAAACAATACTGCGCAATACGCTGAGTTGATGGCAGT

**RNaseH domain**

H K A I E I S P P D E T V L I C T D S F Y I A R G I N E E L
4231 ACATAAAGCAATAGAGATCTCTCCTCCTGATGAAACTGTTCTCATTTGTACTGATTCCTTCTATATAGCTAGAGGAATAAATGAAGAATT

P I W R S N G F L D N K R K P L K H A H R W Q K L A T L L D
4321 GCCAATCTGGCGGTCTAATGGTTTTCTAGATAATAAAAGAAAACCTTTGAAACATGCCCATAGATGGCAAAAATTAGCAACACTTTTGGA

D K P L I T V M H V P G H S K Y G S H V N G N T L V D L L A
4411 TGACAAGCCATTGATTACTGTAATGCACGTGCCAGGCCATTCCAAATATGGATCTCATGTTAACGGGAACACCCTTGTAGACTTACTGGC

K E A M K A S S V C V L T R S Q V K K C L D G E L T Q C I S
4501 AAAGGAAGCAATGAAAGCTTCCTCAGTCTGTGTACTTACCCGGTCACAAGTTAAAAAATGCTTGGATGGGGAATTGACTCAATGCATCAG

P D S I N P K G Y P S A Y D Y A L K D G K C V V T F T N G E
4591 CCCAGATTCCATTAACCCTAAGGGTTACCCTTCTGCCTATGATTATGCCCTAAAAGATGGGAAGTGTGTTGTAACATTCACTAATGGGGA

K R V I P P V D T R P N L I Q E A H N S L G H V H Q G V N A
4681 GAAACGTGTAATACCCCCTGTTGACACAAGACCTAACTTAATACAAGAGGCTCACAATAGTCTTGGGCATGTTCACCAAGGTGTTAATGC

T V E S L Q R S Y W W P G L R K Q V Q R H L A Q C E P C L R
4771 CACAGTTGAATCTTTACAGCGTTCTTACTGGTGGCCTGGACTGCGCAAACAAGTCCAACGGCATTTGGCCCAGTGTGAACCTTGTCTAAG

T N P G P V T R P P Y L K N P K P L S P F D K V Y M D Y I G
4861 AACAAACCCTGGTCCAGTTACCAGACCACCTTACTTAAAAAATCCTAAGCCTCTGTCTCCTTTTGATAAGGTGTATATGGATTATATTGG

P L P P S H G H N H C L V I V D A C T G F V W I Y P T R D Q
4951 TCCTTTGCCACCTTCCCATGGGCATAATCACTGTTTGGTTATTGTTGATGCTTGTACTGGTTTTGTATGGATTTACCCCACACGAGATCA

**Integrase core domain**

S A S T T V R T L T S F I S L G L P C I L H S D K G G A F T
5041 ATCTGCCTCTACCACTGTTAGAACTCTCACCTCCTTCATTTCGCTTGGGCTTCCATGTATCCTACACTCTGACAAGGGAGGTGCCTTCAC

S H Q M Q S F A K S F G I V L E Y S T P Y H P Q S A G V V E
5131 CTCTCACCAAATGCAAAGTTTTGCAAAGAGTTTTGGGATAGTGTTGGAATATAGCACACCTTATCACCCCCAAAGTGCTGGGGTTGTGGA

R K N G E I K R A L M K L L V G R S R Q W Y S L L P L V Q L
5221 AAGGAAAAATGGAGAGATAAAACGAGCTTTAATGAAGCTATTGGTGGGGAGGTCCCGGCAGTGGTATTCTTTGCTTCCTCTGGTACAGCT

G L N N L P R S D C H L T P Y K L L F A K D M T T P L E Q L
5311 TGGACTTAATAACCTTCCTAGGAGTGACTGTCACTTAACTCCTTATAAATTGCTATTTGCAAAGGACATGACCACTCCTTTAGAACAACT

A L S S P I S Q Q E Q L A L I D E L R E E K N P V P I L K V
5401 GGCTTTATCTTCTCCTATCTCTCAACAGGAGCAGTTGGCACTGATAGATGAATTGAGAGAAGAAAAAAACCCTGTACCTATTCTCAAAGT

```
      L  S  P  R  A  V  E  I  Q  T  S  P  G  N  S  K  I  V  S  I  D  N  L  K  R  T  P  I  H  Y
5491 TTTGAGTCCCCGAGCAGTAGAAATTCAGACATCACCAGGTAATTCGAAAATTGTGTCTATTGACAACTTAAAGCGCACTCCCATCCACTA
                                                                          Pol end |
      G  Q  G  N  D  G  L  L  S  N  S  T  S  T  P  T  S  P  T  N  S  T  *
     | Env start
     M  V  R  V  M  M  D  Y  C  Q  I  P  L  Q  P  Q  P  L  P  P  T  A  P  D  Y  Y  N  F  Y  Y
5581 TGGTCAGGGTAATGATGGATTATTGTCAAATTCCACTTCAACCCCAACCTCTCCCACCAACAGCACCTGACTATTATAACTTTTACTATT

     Y  F  F  S  P  S  F  N  M  G  S  Y  C  F  I  T  R  Y  S  N  S  L  E  P  L  R  G  T  Q  W
5671 ATTTTTCAGTCCTTCGTTTAACATGGGCTCATACTGTTTCATTACCCGCTACTCCAATTCATTGGAACCTCTCCGAGGCACACAATGGT

     Y  V  V  Y  R  R  M  H  N  A  R  P  K  R  A  L  H  I  E  V  V  P  V  L  V  E  T  A  G  I
5761 ATGTTGTGTATCGAAGGATGCATAATGCCAGACCTAAACGTGCTCTTCATATAGAGGTTGTTCCAGTACTCGTGGAGACTGCAGGTATTC

     P  F  G  I  I  H  N  P  F  P  K  P  I  V  S  Q  R  S  E  L  L  V  P  F  T  L  N  I  D  T
5851 CATTTGGGATAATTCATAATCCCTTTCCTAAGCCTATTGTTTCTCAACGAAGTGAACTGTTGGTTCCTTTTACTTTAAACATAGACACAA

     R  A  L  A  Y  C  S  G  L  F  S  K  D  A  N  T  H  L  A  K  T  I  E  E  D  L  Q  D  L  D
5941 GAGCACTTGCTTATTGTTCTGGCTTGTTTTCAAAAGATGCTAACACACATCTTGCAAAGACTATAGAAGAAGACTTGCAAGATTTGGACA

     S  R  N  A  H  F  L  V  P  G  T  D  P  W  H  Q  T  S  Y  A  D  K  M  C  F  A  S  Y  G  H
6031 GTAGAAATGCACATTTTCTTGTTCCAGGTACAGATCCTTGGCATCAAACAAGCTATGCTGATAAAATGTGCTTTGCTTCCTATGGACATT

     C  Y  F  V  S  Y  G  K  P  R  K  W  P  R  P  H  V  Y  A  D  H  C  D  R  P  Q  F  W  T  D
6121 GTTATTTTGTTTCCTATGGGAAACCGAGGAAATGGCCAAGACCTCATGTGTATGCAGATCACTGTGACAGACCTCAATTTTGGACTGACA

     I  K  T  A  T  Q  G  L  P  Q  W  Y  L  A  I  D  D  F  S  D  H  L  R  Y  A  K  Q  Q  R  S
6211 TTAAAACTGCAACACAAGGCCTGCCTCAGTGGTACCTAGCTATTGATGATTTTTCTGATCACTTGCGCTATGCTAAGCAGCAACGTTCTG

     G  G  E  D  R  E  Y  R  V  P  G  G  Q  L  P  Y  T  G  A  I  F  C  T  S  F  L  Y  N  T  S
6301 GAGGTGAAGACCGTGAATACAGAGTTCCAGGGGGCCAATTGCCCTATACTGGAGCAATTTTTTGCACATCTTTTTTATATAATACCTCTT

     W  W  D  E  S  N  L  S  V  D  G  S  L  E  L  K  S  I  L  T  S  C  L  A  N  S  T  T  G  K
6391 GGTGGGATGAATCAAATTTATCTGTGGATGGCTCTTTAGAATTAAAGTCCATTTTGACTTCTTGTCTTGCAAATTCTACTACTGGGAAAC

     L  K  P  K  C  L  A  S  Q  W  H  D  N  G  A  N  E  M  F  I  G  V  T  G  T  S  F  C  D  I
6481 TCAAACCTAAGTGTTTAGCCTCTCAATGGCATGATAATGGTGCTAATGAGATGTTTATTGGTGTTACTGGGACTTCATTTTGTGATATCC

     P  R  Y  P  I  F  L  N  R  S  E  S  I  V  S  C  K  S  T  F  V  N  P  R  Q  Q  P  L  E  C
6571 CTAGGTATCCTATTTTTCTCAATAGATCTGAGAGTATTGTATCTTGTAAAAGTACTTTTGTTAATCCTCGACAACAGCCTCTTGAATGTG

     G  N  N  K  T  L  A  A  K  G  L  H  S  W  N  C  G  P  C  S  V  N  I  T  A  N  L  M  G  N
6661 GCAATAATAAAACACTTGCTGCTAAGGGTTTACATTCATGGAATTGTGGGCCATGTAGTGTTAATATTACTGCAAATCTCATGGGTAATT
```

Y T A K E R A S L G N K R W F N L I Q G P L F V N A T P F F

6751 ATACAGCCAAAGAGAGGGCGAGTTTGGGAAATAAGCGTTGGTTTAACTTGATACAAGGTCCATTATTTGTCAACGCTACTCCATTTTTTG

A D N Y A I Y S L Y Q K C K T L S E K Y S L F S V L Q A L E

6841 CTGATAATTATGCCATTTATTCGCTGTATCAGAAATGTAAAACCTTATCTGAAAAATATTCTTTATTTTCAGTGCTTCAAGCATTAGAAG

E F I M V P Q E N E D Y P C T H S C I N A S L L Q M N P K R

6931 AATTTATCATGGTACCTCAGGAAAATGAAGATTACCCTTGTACACATTCTTGTATTAATGCTTCTTTATTACAGATGAACCCCAAAAGAG

A I W G T N K T L N D I H I L A T P D A T S E S F P S S T Y

7021 CAATATGGGGCACAAATAAAACTCTTAATGATATTCATATTTTAGCAACACCTGATGCTACTTCAGAGAGTTTCCCTTCTTCCACTTATA

K S R K I L Q V E S L N N A Q I F R K T N F L L A K S M E K

7111 AATCTAGGAAGATTTTACAAGTAGAATCTTTGAATAATGCACAAATATTTAGAAAAACTAATTTCTTGCTCGCAAAATCTATGGAAAAGA

I S R L Q D A N N I N L R N G V Y L V K D A L T Q V A L I V

7201 TTTCTAGATTGCAGGATGCCAATAATATAAATTTAAGAAATGGAGTTTATTTGGTCAAGGATGCTTTAACTCAAGTGGCCTTGATAGTCA

K H D L A V L S D E L I M E I I V T Q L Q K I I F S L S N G

7291 AACATGATTTGGCAGTTTTATCTGATGAATTGATTATGGAAATTATTGTCACTCAGTTGCAAAAAATTATTTTTTCTTTAAGCAATGGGC

H V P W T I G Y F Q I L R D H S D L H G E Y P P V L T V T Q

7381 ACGTTCCCTGGACTATTGGTTATTTTCAGATATTACGAGATCACAGTGACCTTCACGGTGAATACCCACCTGTGCTCACTGTGACACAGC

H V L R M K L L P D S V L T N P L N M M V Y S Y Y L M E Y S

7471 ACGTGCTTCGAATGAAACTATTACCAGATTCAGTTCTTACTAACCCTCTAAACATGATGGTATATTCCTACTACTTGATGGAATATTCAT

F F Y S F E N F Y Y G L T N W N I L N M G F L V A T G T N I

7561 TTTTTATTCATTTGAAAATTTCTACTATGGTTTAACTAATTGGAATATTTTGAATATGGGTTTTTTAGTAGCTACTGGAACAAATATTG

A H A H M K L H M N M S P Q M F L E T Q H T Y I L T N V K I

7651 CTCATGCTCATATGAAATTACATATGAATATGTCTCCACAGATGTTTTTGGAAACACAACATACTTACATACTGACCAATGTCAAGATTT

L D I C F A N K L H I S P C G L V P P S T T S C P I R I Q S

7741 TGGATATTTGTTTTGCAAACAAATTACACATTTCCCCTTGTGGACTGGTACCACCTAGTACCACTTCTTGTCCTATTAGGATTCAATCAA

S N I S F V F I D S L T N G S Y L I L A G K S E C N I P A L

7831 GTAATATTTCATTTGTTTTTATTGATTCCCTTACAAATGGATCATATTTAATCCTTGCTGGAAAATCTGAATGCAATATTCCAGCTTTGC

Q P S I V T V N T T I T C Y G R Q L F P P P N L G S M H S Q

7921 AACCATCTATAGTAACAGTGAATACCACAATCACTTGTTATGGCAGACAACTTTTCCCACCTCCTAATTTGGGATCAATGCATTCTCAGG

V T F F V P H F S L Q F P L L T G I I A K L Q R S E I T L F

8011 TTACATTTTTTGTACCTCACTTTTCTTTGCAATTTCCACTATTAACTGGCATCATAGCTAAATTGCAAAGGTCAGAAATTACTCTTTTTA

N T H D A T E D I L Q E V K Q L L Q R I D I H E G D F P L W

8101 ATACACATGATGCAACTGAAGATATCTTGCAGGAAGTAAAGCAACTACTTCAAAGGATTGATATACATGAAGGAGATTTTCCTCTGTGGC

L N R L A S A V S A A W P S L A N M A N S I A H A D T S I G

8191 TCAATAGATTGGCTTCAGCTGTTTCTGCAGCCTGGCCTTCTTTAGCTAACATGGCTAATTCCATTGCTCACGCTGATACTTCTATAGGTA

T S I L G T G L Q I L T Y L K P I F I A I V L I V L L I I V

8281 CTTCAATTTTAGGAACGGGTTTACAAATACTTACCTACCTCAAACCAATCTTTATTGCTATAGTGTTGATTGTACTATTAATCATCGTTA

I K I F R F F S G L H L K K I S A P N E D W M S L P S P L V

8371 TTAAGATTTTTAGGTTCTTCTCTGGACTGCATCTGAAAAAAATATCAGCCCCCAACGAAGATTGGATGTCTCTTCCCTCACCACTTGTGA

**Env end |**

K K V P V L C T F L M S S *

8461 AGAAGGTACCAGTGCTTTGTACCTTCCTGATGTCTTCCTGAAGACATGACTGATTGGACATTTGTTGAATTTTCTTTCTTGTATTGTATA

**| ORF1 start**

M Y P V Y R H I C L T I C A S I Y

8551 CTACATTCTTGAAACACATTCTCGCCAAAGAGAGGGTGTCATGTATCCTGTATATAGGCATATATGTTTGACCATCTGTGCTTCAATATA

W K S A S L Y D G F L K E L S Y S D L S T E L L T R Y W K I

8641 TTGGAAATCAGCCTCTCTGTATGATGGCTTCTTAAAAGAGCTGTCCTACAGTGACCTTTCTACTGAACTCTTGACCAGATACTGGAAAAT

R L Y T G I D N V I K S V S Y E I N S F C R L V S R Y M T L

8731 CAGACTGTATACCGGAATAGACAATGTAATCAAGTCTGTCTCTTACGAAATTAACAGTTTCTGTCGACTAGTCTCAAGATACATGACTCT

R Q G R K R P C L F G V Y S L L I L S Y T I G T C L V C I N

8821 CAGGCAGGGCAGGAAAAGGCCTTGTTTATTTGGAGTTTACTCATTGTTAATCCTATCTTATACGATAGGAACCTGTTTAGTATGTATCAA

**3′LTR start** **|ORF1 end |**

R L L T M Y F E K S P S L S A V V T *

8911 CAGGCTTCTTACTATGTATTTTGAGAAAAGCCCCTCCCTATCAGCAGTTGTGACCTGACGTCATTTTTAGCTGACCTCCAAGCATCTTTC

9001 ATTCTATAAGAAGAGAGAGGAAAGATGGATTTTTCAGAAAGCTTCCTTTCCTGAAGCTTTTCTCATCTTCTATCTGGCAGAATCGGGTAA

9091 AGGACTGGGTAATATGTTATATCCAATATGATGCACTGAGTCAACCATAGCAATAATACTTAGAGCATATGTTATATCTTATAGATGTTA

9181 AGATATTTGAATTTGTAACAGCTGTTCTTTGAACAAGACATGTCTTTCTAAGTCATGTACTTTTAACTGGCACTGTTTCTTTTGTACTAT

9271 GCCTTTATTTGAACTTAACTATTGCTTCATATTTTTATAATATTCTTAAAGTCTCATAGCCTCATTTATTTAAATGAATTGGTCTTTTCT

9361 ATATTTTGCTGAGCTGGCTGGCTGATTAGCAAAGTGTAAACTCGAATTCCACATATTGGAAAAGAGCGCCCTATATCCAGACCAGATCAT

**3′LTR end |**

9451 GTCAGGCCTGGACCACTAAACGCAAATTCCTTTTGGGGAAACTGAGGCACAACGCGACAGGCCC

**Fig. S1**. **Detailed descriptions of consensus SpuEFV genome**. Protein Open reading frame (ORF) of gag, pol, env and ORF1 locations were determined by sequence similarity to representative foamy viruses using BLASTP and based on the distribution of stop and start codons, determined by ORFfinder (https://www.ncbi.nlm.nih.gov/orffinder/). The conserved domains were identified by searching the conserved domain database (www.ncbi.nlm.nih.gov/Structure/cdd) and highlighted in darker colors. The SpuEFV PBS is nearly identical to the PBS of human foamy virus (HFV) (SpuEFV is 5'-TGGCACCCATTCGTGGGG-3' and HFV is 5'-TGGCGCCCAACGTGGGG-3'). The inferred pre-substitution nucleotides are indicated in red.

**(a)** **Spuma virus Gag domain (pfam03276)**

**E-value:**9.16e-24

```
SpuEFV_consensus  435 PIVPGvhPAEISVLNMANIRAVTGPTPKDFKEIPMWFESHLSALEAVTSTASPLQKMRLCNSLVPASA--SLIEQECNNW 512
Cdd:pfam03276     291 PVPPP--PPVGAVIPIQHIRSVTGEPPRNPREIPIWLGRNAPAIDGVFPTTTPDLRCRIINALLGGNLglSLTPGDCITW 368

                  513 ESVLANLYVKTHGQVGIADLNEILRKITQEQGIVRAYGVGMKFL-SNHDLIWGILKALCKGDVLKAAIQSKLDLLITEQE 591
                  369 DSAVATLFIRTYGQYPLHQLGNVLKGIADQEGVATAYTLGMMLSgQNYQLVSGIIRGYLPGQAVVTAMQQRLDQEIDDQT 448

                  592 KIRSFPKIVQDIYKTLGRDYLGNNPNRKNLEQEGSKSKKSPSNINSRKKILPQNSQpnyRGKGGKIQGSNRKQFQAQKDQ 671
                  449 RAETFIQHLNAVYEILGLNARGQSIRASVTPQPRPSRGRGRGQSAPEPSQGPVNSG---RGRQCPAPGQNDRGSNIQNQG 525

                  672 DSENPEGAatYDLRKGSQFPHR 693
                  526 QENSSQGG--YNLRSRTYQPQR 545
```

**(b)** **Spuma aspartic protease (A9) domain (pfam03539)**

**E-value:**1.02e-08

```
SpuEFV_consensus  864 EQIKEQIASADCAKNEKTKLRDILYSLKPYFQQFDNQIGHRKIKPHDLSVKT-QPKPQKQYPINKAA 929
Cdd:pfam03539      95 EQQETLLQQSALSKEGKELLKKLFLKYDALWQHWENQVGHRRIKPHKIATGTlKPRPQKQYHINPKA 161
```

**(c)** **foamy virus envelope protein domain (pfam03408)**

**E-value:**1.96e-77

```
SpuEFV_consensus 1922 VYRRMHNARPKRALHIE-VVPVLVET--AGIPFGIIHNPFPKPIVSQRSELLVPFTLNIDTRALAYCSGLfSKDANTHLA 1998
Cdd:pfam03408     111 VYQPLQTRRIARSLRMQhPVPKYIEVnmTSIPQGVYYEPHPEPIVVTERVLGLSQVLMINSENIANNANL-TQEVKKLLA 189

                 1999 KTIEEDLQDLDSRNAHFLVPGTDPWHQTSYADKMCFASYGHCYFVSYGKPRKWPRPHVYADHCDRPQFWTDIKTATQGLP 2078
                  190 EVVNEEMQSLSDVMIDFEIPLGDPRDQEQYIHRKCYQEFAHCYLVKYKTPKSWPTEGLIADQCPLPGYHAGLSYKPQSIW 269

                 2079 QWYLAID-----DFSDHLRYAkQQRSGgedrEYRVPGG--QLPYTGAIFCTSFLYnTSWWDesnlsVDGSLEL-KSILTS 2150
                  270 DYYIKVEitrpaNWSSQAVYG-QARLG----SFYVPKGirQNNYSHVLFCSDQLY-SKWYN-----IENSIEQnEKFLLN 338

                 2151 CLANSTTGK--LKPKCLASQWHDNGANEMFIGVTGTSFCDIPRYPIFLNRSESIVS-----CKSTFVNPRQQPLEC---- 2219
                  339 KLDNLTTGSslLKKRALPKEWSSQGKNALFKEINVLDVCSKPELVILLNTSYYSFSlwegdCNFTKNMISQLVPECegfy 418
```

```
2220 GNNKTLAakgLHSWNCGPC-SVNITANLMGNYTAKERA-------SLGNKRWFNLIQ------GPLFVNATPFFADNYAI 2285
 419 NNSKWMH---MHPYACRFWrSKNEKEETKCRPGEKEKClyypyqdSLESTYDFGFLAyqknfpAPICIEQQEIRDKDYEV 495

2286 YSLYQKCKTLSEKYSLFSVLQALEEFImvpqenedypcTHSciNASLLQMNPKRAIWGTnktLNDIHILATPDATSESFP 2365
 496 YSLYQECKLASKVHGIDTVLFSLKNFL-----------NHT--GRPVNEMPNARAFVGL---VDPKFPPSYPNVTREHYT 559

2366 SSTYKSRKilqvESLNNAQIFRKTNFLLAKSMEKISRLQDANNINLRNGVYLVKDALTQVALIVKHDLAVLSDELIMEII 2445
 560 SCNNRKRR----STDNNYAKLKSMGYALTGAVQTLSQISDINDENLQQGIYLLRDHVITLMEATLHDISVMEGMFAVQHL 635

2446 VTQLQKIIFSLSNGHVPWTI---GYFQILRDHSDlhGEYPPVLTVTQH-VLRMKLLPDSVLTNPLNMMVYsYYLMEYSFF 2521
 636 HTHLNHLKTMLLERRIDWTYmssAWLQQQLQKSD--DEMKVIKRIAKSlVYYVKQTYNSPTATAWEIGLY-YELTIPKHV 712

2522 YsfenfyygLTNWNILNMGFLVATGTNIAHAHMKLHMNMSPQMFLETQHTYiLTNVKILDICFANKLHI-SPCGlVPPST 2600
 713 Y--------LNNWNVVNIGHLVQSAGQLTHVTIAHPYEIINKECTETKYLH-LKDCRRQDYVICDVVEIvQPCG-NSTDT 782

2601 TSCPIRIQSSNISFVFIDSLTNGSYLILAGKSECNIPALQPSIVTVNTTITCYGRQlFPPPNLGSMHSQVTFFVPHFSLQ 2680
 783 SDCPVWAEAVKEPFVQVNPLKNGSYLVLASSTDCQIPPYVPSIVTVNETTSCYGLN-FKKPLVAEERLGFEPRLPNLQLR 861

2681 FPLLTGIIAKLQRSEITLFNTHDATEDILQEVKQLLQRIDIHEGDFPLWLNRLASAVSAAWPSLANMANSIAHADTSIGT 2760
 862 LPHLVGIIAKIKGLKIEVTSSGESIKDQIERAKAELLRLDIHEGDTPAWIQQLAAATKDVWPAAASALQGIGNFLSGAAH 941

2761 SILGTGLQILTYLKP 2775
 942 GIFGTAFSLLGYLKP 956
```

**Figure S2. Conserved domain alignment of the consensus SpuEFV proteins and foamy virus proteins in CDD (Conserved Domain Database).** (a) The alignment of consensus SpuEFV Gag proteins and Spuma virus Gag domain (pfam03276); (b) The alignment of consensus SpuEFV pro proteins and Spuma aspartic protease (A9) domain (pfam03539); (c) The alignment of consensus SpuEFV envelope proteins and foamy virus envelope protein domain (pfam03408). Numbers refer to the position in the consensus SpuEFV each protein or conserved domain. Identical amino acid residues are highlighted in red, and black and blue indicate gaps or different amino acid residues, respectively. The E-value was generated by Conserved Domain search.

**(a)** **Spuma virus Gag domain (pfam03276)**

**E-value:**2.16e-12

```
SpuEFV_QEPC01003194.1  488  RLLNSLVPASA--SLMELECNNLENVLANLYVKTHGXVGIADLNXILRKITQEQGIVRAYGVGMKFL-SNHDLIWGILKT  564
Cdd:pfam03276          346  RIINALLGGNLglSLTPGDCITWDSAVATLFIRTYGQYPLHQLGNVLKGIADQEGVATAYTLGMMLSgQNYQLVSGIIRG  425

                       565  LCKGDALKAAIQSKLDLLITEQEQIRSFPKIVQDIYKILGRDYLGDNPNRKNLEQEgskskn SPSNTNFRKKILPQNSQ-  643
                       426  YLPGQAVVTAMQQRLDQEIDDQTRAETFIQHLNAVYEILGLNARGQSIRASVTPQP------RPSRGRGRGQSAPEPSQg  499

                       644  PNHRGKGGKI----QDSNRKQFQAQKDKDSenpeSAATYDLRN*SQFPHR  689
                       500  PVNSGRGRQCpapgQNDRGSNIQNQGQENS----SQGGYNLRSRTYQPQR  545
```

**(b)** **Spuma aspartic protease (A9) domain (pfam03539)**

**E-value:**8.72e-08

```
SpuEFV_QEPC01003194.1  859  EQIKEQIASADCAKNEKIKLRDILYTLKPYFQQFDNQIGHRKIKPHDLSVKT-QPKPQKQYPINKAA  924
Cdd:pfam03539           95  EQQETLLQQSALSKEGKELLKKLFLKYDALWQHWENQVGHRRIKPHKIATGTlKPRPQKQYHINPKA  161
```

**(c)** **Foamy virus envelope protein domain (pfam03408)**

**E-value:**7.69e-19

```
SpuEFV_QEPC01003194.1 1926  SVLRLTWAHTVSLPATPIHWNLSEahngTVVYRRMHNAKPKRALHIE-VIPVLVET--AGMPFGIIHNPFPKPILnqvnc  2002
Cdd:pfam03408           85  TISRIQWNRDIQVLGPVIDWNVTQ----RAVYQPLQTRRIARSLRMQhPVPKYIEVnmTSIPQGVYYEPHPEPIV-----  155

                      2003  wclll*t*TSER*piVLACSQKKL---------TNILQRL*KMTCEIWTVEMH-----IFLFQVQI---LGIKQAMLIKC  2065
                       156  --------VTER---VLGLSQVLMinseniannANLTQEVKKLLAEVVNEEMQslsdvMIDFEIPLgdpRDQEQYIHRKC  224

                      2066  aLLLIGHCYFVSDGKPRKWPRPHVYADHCDRPQFWTDIKTATQGLPQWYLAID-----DFSDHLRYAkQQRSGgedrEYR  2140
                       225  -YQEFAHCYLVKYKTPKSWPTEGLIADQCPLPGYHAGLSYKPQSIWDYYIKVEitrpaNWSSQAVYG-QARLG----SFY  298

                      2141  VPGG--QLPYTGAIFCTSFLYnTSWWDesnlsVDGSLEL-KSVLTSCLENSTTGK--LKPKCLSSQWHDNGANEMFVGVT  2215
                       299  VPKGirQNNYSHVLFCSDQLY-SKWYN-----IENSIEQnEKFLLNKLDNLTTGSslLKKRALPKEWSSQGKNALFKEIN  372

                      2216  GTSFCDIPRYPIFLNR------------------SESIVSCKSTFVNHQ---QQPLECGTNKTLTAKglHSWNCGPCSV  2273
                       373  VLDVCSKPELVILLNTsyysfslwegdcnftknmiSQLVPECEGFYNNSKwmhMHPYACRFWRSKNEK--EETKCRPGEK  450
```

```
                           2274 NITTNLMGNYTAKERVSLGNKRWFNLIQSPLFVNATPFFADNYAIYSLYQKCKTLSEKHSLFSVLQTLE*FI 2345
                            451 EKCLYYPYQDSLESTYDFGFLAYQKNFPAPICIEQQEIRDKDYEVYSLYQECKLASKVHGIDTVLFSLKNFL 522
```

**E-value:**3.41e-19

```
SpuEFV_QEPC01003194.1 2683 EHAILQHTARALIFNMKLLPDSvltnplnmwyiPT-TXWNIHFFIHLKISNMVYLTNWNILNMGFLVATGTNIAHAHMKL 2761
Cdd:pfam03408          671 EMKVIKRIAKSLVYYVKQTYNS-----------PTaTAWEIGLYYELTIPKHVYLNNWNVVNIGHLVQSAGQLTHVTIAH 739

                      2762 PYEWVSTDVfGNTTYLHTDECQDFGYLFCKQI------------THFPLWVgtt*yyflsy*DSIK*YFICFY*fPYKwi 2829
                       740 PYEIINKEC-TETKYLHLKDCRRQDYVICDVVeivqpcgnstdtSDCPVWA-----------EAVKEPFVQVN--PLK-- 803

                      2830 ifNSCWTI*m*ysgfanIYHNSDYHSHLLWQTAFSTS*fEINAFSGYIFRNSLF--------------------LTGIIA 2889
                       804 --NGSYLV---------LASSTDCQIPPYVPSIVTVN--ETTSCYGLNFKKPLVaeerlgfeprlpnlqlrlphLVGIIA 870

                      2890 KLQRSEITLFNIHDATEDTLQEVKQLLQRIDIHEGDFPLWFNRLATAISAAWPSLASMAHSIAHAATSLGNSILGTGLQI 2969
                       871 KIKGLKIEVTSSGESIKDQIERAKAELLRLDIHEGDTPAWIQQLAAATKDVWPAAASALQGIGNFLSGAAHGIFGTAFSL 950

                      2970 ITYLKP 2975
                       951 LGYLKP 956
```

**Figure S3. Conserved domain alignment of the original SpuEFV proteins and foamy virus proteins in CDD (Conserved Domain Database)**. (a) The alignment of original SpuEFV Gag proteins and Spuma virus Gag domain (pfam03276); (b) The alignment of original SpuEFV pro proteins and Spuma aspartic protease (A9) domain (pfam03539); (c) The two parts of alignment of original SpuEFV envelope proteins and foamy virus envelope protein domain (pfam03408). Numbers refer to the position in the original SpuEFV each protein or conserved domain. Identical amino acid residues are highlighted in red, and black and blue indicate gaps or different amino acid residues, respectively. The E-value was generated by Conserved Domain search.

**Table S1. The information of 28 reptiles used for data mining**

| No. | Species name | Common name | Accession no. |
|---|---|---|---|
| 1 | *Alligator mississippiensis* | American alligator | GCA_000281125.4 |
| 2 | *Alligator sinensis* | Chinese alligator | GCA_000455745.1 |
| 3 | *Gavialis gangeticus* | Gharial | GCA_001723915.1 |
| 4 | *Gekko japonicus* | Schlegel's Japanese gecko | GCA_001447785.1 |
| 5 | *Lacerta bilineata* | Western green lizard | GCA_900245895.1 |
| 6 | *Lacerta viridis* | European green lizard | GCA_900245905.1 |
| 7 | *Paroedura picta* | Ocelot gecko | GCA_003118565.1 |
| 8 | *Sphenodon punctatus* | Tuatara | GCA_003113815.1 |
| 9 | *Anolis carolinensis* | Green anole | GCA_000090745.2 |
| 10 | *Apalone spinifera* | Spiny softshell turtle | GCA_000385615.1 |
| 11 | *Chelonia mydas* | Green sea turtle | GCA_000344595.1 |
| 12 | *Chrysemys picta* | Painted turtle | GCA_000241765.2 |
| 13 | *Crocodylus porosus* | Saltwater crocodile | GCA_001723895.1 |
| 14 | *Crotalus horridus* | Timber rattlesnake | GCA_001625485.1 |
| 15 | *Crotalus pyrrhus* | Southwestern speckled rattlesnake | GCA_000737285.1 |
| 16 | *Crotalus viridis* | Prairie rattlesnake | GCA_003400415.1 |
| 17 | *Gopherus agassizii* | Desert tortoise | GCA_002896415.1 |
| 18 | *Malaclemys terrapin* | Diamondback terrapin | GCA_001728815.2 |
| 19 | *Ophiophagus hannah* | King cobra | GCA_000516915.1 |
| 20 | *Pantherophis guttatus* | Corn snake | GCA_001185365.1 |
| 21 | *Pelodiscus sinensis* | Chinese softshell turtle | GCA_000230535.1 |
| 22 | *Pogona vitticeps* | Central bearded dragon | GCA_900067755.1 |
| 23 | *Protobothrops flavoviridis* | Okinawa habu | GCA_003402635.1 |
| 24 | *Protobothrops mucrosquamatus* | Brown spotted pit viper | GCA_001527695.3 |
| 25 | *Python bivittatus* | Burmese python | GCA_000186305.2 |
| 26 | *Terrapene mexicana* | Mexican box turtle | GCA_002925995.2 |
| 27 | *Thamnophis sirtalis* | Common garter snake | GCA_001077635.2 |
| 28 | *Vipera berus* | Common European adder | GCA_000800605.1 |

**Table S2. The information of 130 bird genomes used for data mining**

| No. | Species name | Common name | Accession no. |
|---|---|---|---|
| 1 | Acanthisitta chloris | Rifleman | GCA_000695815.1 |
| 2 | Acridotheres javanicus | Javan myna | GCA_002849675.1 |
| 3 | Agapornis roseicollis | Peach-faced lovebird | GCA_002631895.1 |
| 4 | Amazona aestiva | Blue-fronted amazon | GCA_001420675.1 |
| 5 | Amazona vittata | Puerto Rican parrot | GCA_000332375.1 |
| 6 | Anas platyrhynchos | Mallard | GCA_000355885.1 |
| 7 | Anas zonorhyncha | Eastern spot-billed duck | GCA_002224875.1 |
| 8 | Anser brachyrhynchus | Pink-footed goose | GCA_002592135.1 |
| 9 | Anser cygnoides domesticus | Domestic goose | GCA_000971095.1 |
| 10 | Antrostomus carolinensis | Chuck-will's-widow | GCA_000700745.1 |
| 11 | Apaloderma vittatum | Bar-tailed trogon | GCA_000703405.1 |
| 12 | Aptenodytes forsteri | Emperor penguin | GCA_000699145.1 |
| 13 | Apteryx australis mantelli | Southern brown kiwi | GCA_001039765.1 |
| 14 | Apteryx haastii | Great spotted kiwi | GCA_003342985.1 |
| 15 | Apteryx owenii | Little spotted kiwi | GCA_003342965.1 |
| 16 | Apteryx rowi | Okarito brown kiwi | GCA_003343035.1 |
| 17 | Aquila chrysaetos canadensis | Golden eagle | GCA_000766835.1 |
| 18 | Ara macao | Scarlet macaw | GCA_000400695.1 |
| 19 | Athene cunicularia | Burrowing owl | GCA_003259725.1 |
| 20 | Balearica regulorum gibbericeps | East African grey crowned-crane | GCA_000709895.1 |
| 21 | Bambusicola thoracicus | Chinese bamboo-partridge | GCA_002909625.1 |
| 22 | Buceros rhinoceros silvestris | Rhinoceros hornbill | GCA_000710305.1 |
| 23 | Calidris pugnax | Ruff | GCA_001431845.1 |
| 24 | Calidris pygmaea | Spoon-billed sandpiper | GCA_003697955.1 |
| 25 | Callipepla squamata | Scaled quail | GCA_002218305.1 |
| 26 | Calypte anna | Anna's hummingbird | GCA_000699085.1 |
| 27 | Cariama cristata | Red-legged seriema | GCA_000690535.1 |
| 28 | Casuarius casuarius | Southern cassowary | GCA_003342895.1 |
| 29 | Cathartes aura | Turkey vulture | GCA_000699945.1 |
| 30 | Chaetura pelagica | Chimney swift | GCA_000747805.1 |
| 31 | Charadrius vociferus | Killdeer | GCA_000708025.2 |
| 32 | Chlamydotis macqueenii | Macqueen's bustard | GCA_000695195.1 |
| 33 | Chlamydotis undulata undulata | Houbara bustard | GCA_003400225.1 |
| 34 | Chrysolophus pictus | Golden pheasant | GCA_003413605.1 |
| 35 | Cicinnurus regius | King bird of paradise | GCA_003713305.1 |
| 36 | Ciconia boyciana | Oriental stork | GCA_002002965.1 |
| 37 | Colinus virginianus | Northern bobwhite | GCA_000599465.2 |
| 38 | Colius striatus | Speckled mousebird | GCA_000690715.1 |
| 39 | Columba livia | Rock pigeon | GCA_000337935.1 |
| 40 | Corvus brachyrhynchos | American crow | GCA_000691975.1 |
| 41 | Corvus cornix cornix | Hooded crow | GCA_000738735.2 |
| 42 | Corvus hawaiiensis | Hawaiian crow | GCA_003402825.1 |

| | | | |
|---|---|---|---|
| 43 | Coturnix japonica | Japanese quail | GCA_001577835.1 |
| 44 | Crypturellus cinnamomeus | Thicket tinamou | GCA_003342915.1 |
| 45 | Cuculus canorus | Common cuckoo | GCA_000709325.1 |
| 46 | Cyanistes caeruleus | Blue tit | GCA_002901205.1 |
| 47 | Diphyllodes magnificus | Magnificent bird-of-paradise | GCA_003713285.1 |
| 48 | Dromaius novaehollandiae | Emu | GCA_003342905.1 |
| 49 | Egretta garzetta | Little egret | GCA_000687185.1 |
| 50 | Empidonax traillii | Willow flycatcher | GCA_003031625.1 |
| 51 | Eopsaltria australis | Eastern yellow robin | GCA_003426825.1 |
| 52 | Erythrura gouldiae | Gouldian finch | GCA_003676055.1 |
| 53 | Eudromia elegans | Elegant crested-tinamou | GCA_003342815.1 |
| 54 | Eurypyga helias | Sunbittern | GCA_000690775.1 |
| 55 | Falco cherrug | Saker falcon | GCA_000337975.1 |
| 56 | Falco peregrinus | Peregrine falcon | GCA_000337955.1 |
| 57 | Ficedula albicollis | Collared flycatcher | GCA_000247815.2 |
| 58 | Fulmarus glacialis | Northern fulmar | GCA_000690835.1 |
| 59 | Gallirallus okinawae | Okinawa rail | GCA_002003005.1 |
| 60 | Gallus gallus | Chicken | GCA_000002315.5 |
| 61 | Gavia stellata | Red-throated loon | GCA_000690875.1 |
| 62 | Geospiza fortis | Medium ground-finch | GCA_000277835.1 |
| 63 | Grus japonensis | Red-crowned crane | GCA_002002985.1 |
| 64 | Haliaeetus albicilla | White-tailed eagle | GCA_000691405.1 |
| 65 | Haliaeetus leucocephalus | Bald eagle | GCA_000737465.1 |
| 66 | Hemignathus virens | Hawaii amakihi | GCA_003286495.1 |
| 67 | Hirundo rustica rustica | Barn swallow | GCA_003692655.1 |
| 68 | Junco hyemalis | Dark-eyed junco | GCA_003829775.1 |
| 69 | Lepidothrix coronata | Blue-crowned manakin | GCA_001604755.1 |
| 70 | Leptosomus discolor | Cuckoo roller | GCA_000691785.1 |
| 71 | Limosa lapponica baueri | Bar-tailed godwit | GCA_002844005.1 |
| 72 | Lonchura striata domestica | Bengalese finch | GCA_002197715.1 |
| 73 | Lyrurus tetrix tetrix | Black grouse | GCA_000586395.1 |
| 74 | Manacus vitellinus | Golden-collared manakin | GCA_001715985.2 |
| 75 | Meleagris gallopavo | Turkey | GCA_000146605.3 |
| 76 | Melopsittacus undulatus | Budgerigar | GCA_000238935.1 |
| 77 | Merops nubicus | Carmine bee-eater | GCA_000691845.1 |
| 78 | Mesitornis unicolor | Brown roatelo | GCA_000695765.1 |
| 79 | Mixornis gularis | Striped tit-babbler | GCA_003546035.1 |
| 80 | Nannopterum auritus | Double-crested cormorant | GCA_002173455.1 |
| 81 | Nannopterum brasilianus | Neotropic cormorant | GCA_002174335.1 |
| 82 | Nannopterum harrisi | Galapagos flightless cormorant | GCA_002173475.1 |
| 83 | Nestor notabilis | Kea | GCA_000696875.1 |
| 84 | Nipponia nippon | Crested ibis | GCA_000708225.1 |
| 85 | Nothoprocta perdicaria | Chilean tinamou | GCA_003342845.1 |
| 86 | Numida meleagris | Helmeted guineafowl | GCA_002078875.2 |

| | | | |
|---|---|---|---|
| 87 | Opisthocomus hoazin | Hoatzin | GCA_000692075.1 |
| 88 | Paradisaea raggiana | Raggiana bird of paradise | GCA_003713265.1 |
| 89 | Paradisaea rubra | Red bird of paradise | GCA_003713215.1 |
| 90 | Parotia lawesii | Lawes's parotia | GCA_003713295.1 |
| 91 | Parus major | Great tit | GCA_001522545.2 |
| 92 | Passer domesticus | House sparrow | GCA_001700915.1 |
| 93 | Patagioenas fasciata monilis | Band-tailed pigeon | GCA_002029285.1 |
| 94 | Pelecanus crispus | Dalmatian pelican | GCA_000687375.1 |
| 95 | Phaethon lepturus | White-tailed tropicbird | GCA_000687285.1 |
| 96 | Phalacrocorax carbo | Great cormorant | GCA_000708925.1 |
| 97 | Phoenicopterus ruber ruber | American flamingo | GCA_000687265.1 |
| 98 | Phylloscopus plumbeitarsus | Two-barred warbler | GCA_001655115.1 |
| 99 | Phylloscopus trochiloides viridanus | Greenish warbler | GCA_001655095.1 |
| 100 | Phylloscopus trochilus acredula | Phylloscopus trochilus acredula | GCA_002305835.1 |
| 101 | Picoides pubescens | Downy woodpecker | GCA_000699005.1 |
| 102 | Podiceps cristatus | Great crested grebe | GCA_000699545.1 |
| 103 | Pseudopodoces humilis | Tibetan ground-tit | GCA_000331425.1 |
| 104 | Psittacula krameri | Rose-ringed parakeet | GCA_002870145.1 |
| 105 | Pterocles gutturalis | Yellow-throated sandgrouse | GCA_000699245.1 |
| 106 | Pterocnemia pennata | Darwin's rhea | GCA_003342835.1 |
| 107 | Pygoscelis adeliae | Adelie penguin | GCA_000699105.1 |
| 108 | Pygoscelis antarcticus | Chinstrap penguin | GCA_003264595.1 |
| 109 | Pygoscelis papua | Gentoo penguin | GCA_003264615.1 |
| 110 | Rhea americana | Greater rhea | GCA_003343005.1 |
| 111 | Saxicola maurus maurus | Siberian stonechat | GCA_900205225.1 |
| 112 | Serinus canaria | Common canary | GCA_000534875.1 |
| 113 | Setophaga coronata coronata | Yellow-rumped warbler | GCA_001746935.1 |
| 114 | Spheniscus humboldti | Humboldt's penguin | GCA_003264545.1 |
| 115 | Spheniscus magellanicus | Magellanic penguin | GCA_003264715.1 |
| 116 | Spheniscus mendiculus | Galapagos penguin | GCA_003264655.1 |
| 117 | Sporophila hypoxantha | Tawny-bellied seedeater | GCA_002167245.1 |
| 118 | Strix occidentalis caurina | Spotted owl | GCA_002372975.1 |
| 119 | Struthio camelus australis | African ostrich | GCA_000698965.1 |
| 120 | Sturnus vulgaris | Common starling | GCA_001447265.1 |
| 121 | Syrmaticus mikado | Mikado pheasant | GCA_003435085.1 |
| 122 | Taeniopygia guttata | Zebra finch | GCA_000151805.2 |
| 123 | Tauraco erythrolophus | Red-crested turaco | GCA_000709365.1 |
| 124 | Tinamus guttatus | White-throated tinamou | GCA_000705375.2 |
| 125 | Tympanuchus cupido pinnatus | Greater prairie chicken | GCA_001870855.1 |
| 126 | Tyto alba | Barn owl | GCA_000687205.1 |
| 127 | Uria lomvia | Thick-billed guillemot | GCA_002289315.1 |
| 128 | Urile pelagicus | Pelagic cormorant | GCA_002173435.1 |
| 129 | Zonotrichia albicollis | White-throated sparrow | GCA_000385455.1 |
| 130 | Zosterops lateralis melanops | Silver-eye | GCA_001281735.1 |

**Table S3. The information of representative retroviruses used for phylogenetic analysis.**

| No. | Virus name | Genus | Abbreviation | Natural host | Accession no. |
|---|---|---|---|---|---|
| 1 | Avian leukemia virus | Alpharetrovirus | ALV | Chicken | NC_015116 |
| 2 | Lymphoproliferative disease virus | Alpharetrovirus | LDV | Turkey | U09568 |
| 3 | Mouse mammary tumor virus | Betaretrovirus | MMTV | Mouse | NC_001503 |
| 4 | Mason-Pfizer monkey virus | Betaretrovirus | MPMV | Primate | NC_001550 |
| 5 | Simian retrovirus 1 | Betaretrovirus | SRV1 | Primate | M11841 |
| 6 | Bovine leukemia virus | Deltaretrovirus | BLV | Cattle | NC_001414 |
| 7 | Human T-lymphotropic virus 1 | Deltaretrovirus | HTLV1 | Human | NC_001436 |
| 8 | Simian T-lymphotropic virus 2 | Deltaretrovirus | STLV2 | Non-human primate | NC_001815 |
| 9 | Walleye dermal sarcoma virus | Epsilonretrovirus | WDSV | Fish | NC_001867 |
| 10 | Walleye epidermal hyperplasia virus type 1 | Epsilonretrovirus | WEHV1 | Fish | AF133051 |
| 11 | Walleye epidermal hyperplasia virus type 2 | Epsilonretrovirus | WEHV2 | Fish | AF133052 |
| 12 | Atlantic salmon swim bladder sarcoma virus | Gamma-epsilon | SSSV | Atlantic salmon | NC_007654 |
| 13 | Feline leukemia virus | Gammaretrovirus | FeLV | Cat | NC_001940 |
| 14 | Friend murine leukemia virus | Gammaretrovirus | F-MuLV | Mouse | NC_001362 |
| 15 | Mus dunni endogenous retrovirus | Gammaretrovirus | MDEV | Mouse | AF053745 |
| 16 | Porcine endogenous retrovirus B | Gammaretrovirus | PERV-B | Pig | AY099324 |
| 17 | Rhinolophus ferrumequinum retrovirus | Gammaretrovirus | RfRV | Greater horseshoe bat | JQ303225 |
| 18 | Equine infectious anemia virus | Lentivirus | EIAV | Horse | NC_001450 |
| 19 | Feline immunodeficiency virus | Lentivirus | FIV | Cat | NC_001482 |
| 20 | Human immunodeficiency virus 1 | Lentivirus | HIV1 | Human | NC_001802 |
| 21 | Visna/Maedi virus | Lentivirus | VMV | Sheep | NC_001452 |
| 22 | Amphilophus citrinellus fomy-like virus | Spuma-like | AciFLERV | Fish (midas cichlid) | Reference.1 |
| 23 | Austrofundulus limnaeus fomy-like virus | Spuma-like | AliFLERV | Fish (annual killifish ) | Reference.1 |

| | | | | | |
|---|---|---|---|---|---|
| 24 | Notophthalmus viridescens | Spuma-like | NviFLERV | Amphibian(eastern newt) | Reference.1 |
| 25 | latyfish endogenous retrovirus | Spuma-like | PlatyfishEFV | Fish (platyfish) | Reference.2 |
| 26 | Danio rerio foamy virus | Spuma-like | DrFV-1 | Zebrafish | CABZ01054182 |
| 27 | Bovine foamy virus | Spumavirus | BFV | Cattle | NC_001831 |
| 28 | Equine foamy virus | Spumavirus | EFV | Horse | NC_002201 |
| 29 | Feline foamy virus | Spumavirus | FFV | Cat | NC_001871 |
| 30 | Coelacanth endogenous foamy-like virus | Spumavirus | CoeEFV | Fish (coelacanth ) | Reference.3 |
| 31 | Brown greater galago prosimian foamy virus | Spumavirus | PSFVgal | Greater galago | KM233624 |
| 32 | White-tufted-ear marmoset simian foamy virus | Spumavirus | SFVmar | Common marmoset | GU356395 |
| 33 | Squirrel monkey simian foamy virus | Spumavirus | SFVsqu | Squirrel monkey | GU356394 |
| 34 | Orangutan Simian foamy virus | Spumavirus | SFVora | Pongo pygmaeus pygmaeus | AJ544579 |
| 35 | Macaque simian foamy virus | Spumavirus | SFVmac | Macaque | NC_010819 |
| 36 | African green monkey simian foamy virus | Spumavirus | SFVagm | African green monkey | NC_010820 |
| 37 | Western chimpanzee simian foamy virus | Spumavirus | SFVcpz | Western chimpanzee | NC_001364 |
| 38 | Western lowland gorilla simian foamy virus | Spumavirus | SFVgor | Western lowland gorilla | NC_039029 |
| 39 | Spider monkey simian foamy virus | Spumavirus | SFVspm | Spider monkey | NC_039027 |
| 40 | Sloth endogenous virus | Spumavirus | SloEFV | Sloth | Reference.4 |
| 41 | Snakehead retrovirus | Unclassified | SnRV | Fish (snakehead fish) | NC_001724 |

Reference.1 Aiewsakun P, Katzourakis A. 2017. Marine origin of retroviruses in the early Palaeozoic Era. Nat Commun 8:13954.

Reference.2 Katzourakis A, Gifford RJ, Tristem M, Gilbert MT, Pybus OG. 2009. Macroevolution of complex retroviruses. Science 325:1512.

Reference.3 Han GZ, Worobey M. 2012. An endogenous foamy-like viral element in the coelacanth genome. PLoS Pathog 8:e1002790.

Reference.4 Schartl M, Walter RB, Shen Y, Garcia T, Catchen J, Amores A, Braasch I, Chalopin D, Volff JN, Lesch KP, et al. 2013. The genome of the platyfish, Xiphophorus maculatus, provides insights into evolutionary adaptation and several complex traits. Nat Genet 45:567-572.

**Table S4. The matching contigs identified in S*phenodon punctatus g*enome.**

| Accession number | The length of contigs | The number of hits* |
|---|---:|:---:|
| QEPC01000046.1 | 2,261,982 | 1 |
| QEPC01000755.1 | 2,726,839 | 1 |
| QEPC01000841.1 | 6,575,273 | 2 |
| QEPC01000971.1 | 267,574 | 1 |
| QEPC01000978.1 | 4,485,343 | 1 |
| QEPC01001126.1 | 3,925,424 | 1 |
| QEPC01001380.1 | 1,282,949 | 1 |
| QEPC01001501.1 | 2,970,112 | 1 |
| QEPC01001533.1 | 17,747,792 | 2 |
| QEPC01001717.1 | 8,015,171 | 4 |
| QEPC01002018.1 | 3,274,960 | 1 |
| QEPC01002047.1 | 3,931,782 | 1 |
| QEPC01002186.1 | 2,508,909 | 1 |
| QEPC01002218.1 | 1,036,309 | 1 |
| QEPC01002219.1 | 948,854 | 1 |
| QEPC01002862.1 | 2,112,741 | 2 |
| QEPC01003194.1 | 8,830,396 | 1 |
| QEPC01003216.1 | 2,422,517 | 1 |
| QEPC01003596.1 | 1,075,095 | 1 |
| QEPC01003632.1 | 3,332,090 | 1 |
| QEPC01003681.1 | 2,869,604 | 1 |
| QEPC01003743.1 | 7,791,302 | 2 |
| QEPC01004016.1 | 1,026,366 | 1 |
| QEPC01004481.1 | 1,713,101 | 1 |
| QEPC01004587.1 | 2,591,693 | 2 |
| QEPC01004588.1 | 2,510,880 | 1 |
| QEPC01004609.1 | 304,660 | 1 |
| QEPC01004754.1 | 998,333 | 2 |
| QEPC01004817.1 | 5,636,056 | 2 |
| QEPC01004976.1 | 1,923,601 | 1 |
| QEPC01005318.1 | 1,812,687 | 1 |
| QEPC01005364.1 | 927,510 | 1 |
| QEPC01005564.1 | 1,094,855 | 1 |
| QEPC01005592.1 | 430,143 | 1 |
| QEPC01006022.1 | 1,470,196 | 1 |
| QEPC01006337.1 | 853,782 | 1 |
| QEPC01006337.1 | 853,782 | 1 |
| QEPC01006515.1 | 867,373 | 1 |
| QEPC01006862.1 | 2,177,237 | 2 |
| QEPC01007003.1 | 1,535,892 | 1 |

| | | |
|---|---|---|
| QEPC01007237.1 | 6,544,613 | 2 |
| QEPC01007433.1 | 3,616,258 | 2 |
| QEPC01007453.1 | 1,560,635 | 1 |
| QEPC01007467.1 | 4,402,635 | 2 |
| QEPC01007499.1 | 9,879,472 | 1 |
| QEPC01007621.1 | 1,817,577 | 1 |
| QEPC01007748.1 | 3,321,910 | 1 |
| QEPC01007812.1 | 4,155,962 | 1 |
| QEPC01007842.1 | 1,455,502 | 1 |
| QEPC01008061.1 | 4,176,174 | 1 |
| QEPC01008071.1 | 3,181,871 | 1 |
| QEPC01008173.1 | 12,535,514 | 1 |
| QEPC01008309.1 | 13,133,864 | 1 |
| QEPC01008451.1 | 3,157,202 | 2 |
| QEPC01008452.1 | 4,018,016 | 1 |
| QEPC01008871.1 | 1,592,225 | 1 |
| QEPC01009045.1 | 1,866,647 | 1 |
| QEPC01009046.1 | 1,866,647 | 1 |
| QEPC01009539.1 | 190,299 | 1 |
| QEPC01009586.1 | 505,356 | 1 |
| QEPC01009783.1 | 1,171,668 | 2 |
| QEPC01010218.1 | 1,935,863 | 1 |
| QEPC01010231.1 | 2,180,379 | 2 |
| QEPC01010332.1 | 3,049,825 | 1 |
| QEPC01010651.1 | 2,503,641 | 1 |
| QEPC01010723.1 | 1,079,134 | 1 |
| QEPC01011263.1 | 1,245,951 | 1 |
| QEPC01011712.1 | 4,733,220 | 1 |
| QEPC01011882.1 | 3,029,500 | 2 |
| QEPC01011901.1 | 29,987,930 | 1 |
| QEPC01012034.1 | 449,693 | 2 |
| QEPC01012093.1 | 582,806 | 2 |
| QEPC01012163.1 | 1,035,380 | 1 |
| QEPC01012515.1 | 1,668,146 | 1 |
| QEPC01012627.1 | 76,116 | 1 |
| QEPC01012853.1 | 722,533 | 1 |
| QEPC01012962.1 | 1,874,977 | 2 |
| QEPC01013045.1 | 5,599,944 | 1 |
| QEPC01013253.1 | 1,321,719 | 2 |
| QEPC01013568.1 | 4,769,775 | 1 |
| QEPC01013677.1 | 2,261,225 | 1 |
| QEPC01013962.1 | 978,021 | 1 |
| QEPC01013985.1 | 1,532,209 | 1 |

| | | |
|---|---|---|
| QEPC01014049.1 | 917,795 | 1 |
| QEPC01014179.1 | 180,213 | 1 |
| QEPC01014350.1 | 2,578,219 | 1 |
| QEPC01014606.1 | 3,568,174 | 2 |
| QEPC01014636.1 | 2,612,062 | 1 |
| QEPC01014787.1 | 1,793,453 | 1 |
| QEPC01015038.1 | 2,158,932 | 1 |
| QEPC01015112.1 | 1,559,665 | 1 |
| QEPC01015139.1 | 3,320,937 | 1 |
| QEPC01015182.1 | 2,270,417 | 1 |
| QEPC01015561.1 | 3,676,817 | 1 |
| **Total** | | 117 |

* The hits within 30kb are regarded as one hit.

**Table S5. The important SpuEFV sequences used for constructing consensus genome**

| Contig number | Contig size (bp) | Location (start-end) | Genomic region present |
|---|---|---|---|
| QEPC01000046.1 | 2,261,982 | 2082874-2085549 | pol |
| QEPC01000841.1 | 6,575,273 | 6473051-6474912 | env |
| QEPC01001126.1 | 3,925,424 | 1140946-1146409 | 5'LTR-gag-pol |
| QEPC01001126.1 | 3,925,424 | 1150105-1150972 | env |
| QEPC01001501.1 | 2,970,112 | 587272-591846 | pol-env |
| QEPC01001501.1 | 2,970,112 | 990495-991671 | 5'LTR-gag |
| QEPC01001533.1 | 17,747,792 | 7844865-7846050 | env |
| QEPC01001717.1 | 8,015,171 | 3767132-3769195 | env |
| QEPC01002186.1 | 2,508,909 | 2432408-2433392 | env |
| QEPC01002219.1 | 948,854 | 864881-870322 | 5'LTR-gag-pol |
| QEPC01003194.1 | 8,830,396 | 6038514-6048628 | 5'LTR-gag-pol-env-3'LTR |
| QEPC01003216.1 | 2,422,517 | 542585-543835 | env |
| QEPC01003596.1 | 1,075,095 | 148116-150161 | env |
| QEPC01003632.1 | 3,332,090 | 1919183-1920342 | 5'LTR-gag |
| QEPC01003743.1 | 7,791,302 | 2125690-2126673 | env |
| QEPC01003743.1 | 7,791,302 | 4870735-4875331 | 5'LTR-gag-pol |
| QEPC01004481.1 | 1,713,101 | 1529186-1534239 | pol-env |
| QEPC01004609.1 | 304,660 | 79998-82854 | gag-pol |
| QEPC01004754.1 | 998,333 | 809887-813532 | 5'LTR-gag-pol |
| QEPC01004976.1 | 1,923,601 | 702186-707162 | 5'LTR-gag-pol |
| QEPC01006337.1 | 853,782 | 263997-266031 | 5'LTR-gag |
| QEPC01006776.1 | 1,820,561 | 995473-996768 | env |
| QEPC01006862.1 | 2,177,237 | 1552398-1555851 | gag-pol |
| QEPC01007467.1 | 4,402,635 | 2526645-2528709 | env |
| QEPC01007467.1 | 4,402,635 | 2526836-2532419 | pol-env |
| QEPC01007467.1 | 4,402,635 | 2528830-2534236 | 5'LTR-gag-pol |
| QEPC01007499.1 | 9,879,472 | 4175270-4179532 | pol-env |
| QEPC01007812.1 | 4,155,962 | 3366177-3370732 | pol-env |
| QEPC01007842.1 | 1,455,502 | 628428-630375 | 5'LTR-gag |
| QEPC01007989.1 | 1,160,342 | 1016139-1017692 | gag-pol |
| QEPC01008041.1 | 6,575,273 | 6471648-6481808 | 5'LTR-gag-pol-env-3'LTR |
| QEPC01008061.1 | 4,176,174 | 879218-881207 | 5'LTR-gag |
| QEPC01009046.1 | 1,866,647 | 2304064-2305587 | pol |
| QEPC01010218.1 | 1,935,863 | 720863-722618 | env |
| QEPC01012034.1 | 449,693 | 259221-263659 | gag-pol |
| QEPC01012163.1 | 1,035,380 | 925517-929611 | 5'LTR-gag-pol |
| QEPC01012627.1 | 76,116 | 40790-45921 | 5'LTR-gag-pol |
| QEPC01012897.1 | 3,063,928 | 2695003-2698029 | env |
| QEPC01013045.1 | 5,599,944 | 3726992-3732001 | 5'LTR-gag-pol |
| QEPC01014179.1 | 180,213 | 153493-156634 | env |

**Data set S1. The alignments used to build the phylogenetic trees of Pol represented in Fig. 2.**

Pol
58 490
AciFLERV YPLNPGAVKE MDLIVRELLT LGVIRQENPI TNSPIQAVKK PEGWRPVINF
AliFLERV YPLHPEAVAE MDKIVKELHA LGIIREENPL TNSPIQAVKK PEGWRPVINF
PlatyfishEFV YPLNPGAVEE MDKIVKELGA LEIIREENPI TNSPIQAVKK PEGWRPVINF
DrFV-1 YPINRNALPE IKITIEELKR KGIIKEVNAP TNSPIQAVAK PDGWRLVTNY
BFV YHINPRAKAD IQIVIDDLLR QGVLRQQNSE MNTPVYPVPK ADGWRMVLDY
EFV YRINPKAKAD IQIVIDDLLK QGVLKQQTSP MNTPVYPVPK PDGWRMVLDY
EFVora YPINPKAKES IQIVINDLLK QGVLIQQNSI MNTPVYPVPK PDGWRMVLDY
SFVcpz YPINPKAKPS IQIVIDDLLK QGVLTPQNST MNTPVYPVPK PDGWRMVLDY
SFVgor YPINPKARSS IQVVIDDLLK QGVLVQQNST MNTPVYPIPK PDGWGMVLDY
MSFV YPINPKAKPS IQIVIDDLLK QGVLIQQNST MNTPVYPVPK PDGWRMVLDY
SFVagm YPINPKAKAS IQTVINDLLK QGVLIQQNSI MNTPVYPVPK PDGWRMVLDY
SFVmar YHINTKAKPS IQQVIDDLLK QGVLIKQTSV MNTPIYPVPK PDGWRMVLDY
SFVspm YHINPKAKPS IQIVINDLLK QGVLRQSTSP MNTPVYPVPK PDGWRMVLDY
SFVsqu YHINPKAKPS IQIVINDLLK QGVLKQQNSI MNTPIYPVPK TEGWRMVLDY
PSFVgal YHVNPAARPD IQIVIDDLLK QGVLKKQNSP MNTPVYPVPK PDGWRMVLDY
FFV YHINPKAKPD IQIVINDLLK QGVLIQKEST MNTPVYPVPK PNGWRMVLDY
SloEFV YHINYKAKLA IQTVINDLIK QGVLLHQNSS MNTPIYPVPK TNGWRMVLNF
QEPC01001126.1 YPFNKTAIND IQKVIDDLIA QGALIRQYSS MNTPVYPLPK PNGWCMVLDY
QEPC01004481.1 YPINKAAIND IQKVIDYLIA QGALIKQYSS MNTPVYPVPK PNGWCMVLDY
QEPC01002018.1 YPINKAAIND IQKVINDLIA QGALIKQYST MNTPVYPVPK PNGWRMVLDY
QEPC01007433.1_1 YPINKAAIND IQKVINDLIA QGALIKQYST MNTPVYPVPK PNGWRMVLDY
QEPC01007433.1_2 YPINKAAIND IQKVINDLIA QGALIKQYST MNTPVYPVPK PNGWRMVLDY
QEPC01004609.1 YPINKAAIND IQKVINDLIA QGALIKQYST MNTPVYPVPK PNGWRMVLDY
QEPC01008451.1_1 YPINKAAIND IQKVINDLIA QGALIKQYST MNTPVYPVPK PNGWRMVLDY
QEPC01008451.1_2 YPINKAAIND IQKVINDLIA QGALIKQYST MNTPVYPVPK PNGWRMVLDY
QEPC01004754.1_1 YPINKAAIND IQKVINDLIA QGALIKQYST MNTPVYPVPK PNGWRMVLDY
QEPC01004754.1_2 YPINKAAIND IQKVINDLIA QGALIKQYST MNTPVYPVPK PNGWRMVLDY
QEPC01010651.1 YPINKAAIND IQKVINDLIA QGALIKQYST MNTPVYPVPK PNGWRMVLDY
QEPC01009045.1 YPINKAAIND IQNVINDLIA QGALIKQYST MNTPVYPVPK PNGWRMVLDY
QEPC01012034.1 YPINKAAIND IQKVINDLIA EGALIRQYSS MNTPVYPVPK PNGWRMVLDY
QEPC01012853.1 YPINKAAIDD IQKAINDLIA QGALIKQDST MNTPVYPIPK PNGWRMVLDY
QEPC01013253.1_1 HPINKTAIND IQKVIDDLIA QGALIRQYSS MNTSVYPVPK PNGWHMVIDY
QEPC01013253.1_2 HPINKTAIND IQKVIDDLIA QGALIRQYSS MNTSVYPVPK PNGWHMVIDY
QEPC01007842.1 YSINKAAIND IQKLINYLIA QGVLIRQYSS MNTPVYPVPK PNGWCMVLDY
CoeEFV YPINTKAIPS IQVVINELLE QGVLVKQTSP TNMAVLPVPK PDGWRLVLDY
NviFLERV YPIKREAKAS VKEILTHLEN QDVIESCTSD MNNPLFPVAK PDHYHIVLDY
FeLV YPMPHEAYQG IKPHIRRMLD QGILKPCQSP WNTPLLPVKK PGEYRPVQDL
F-MuLV YPMSQEARLG IKPHIQRLLD QGILVPCQSP WNTPLLPVKK PGNYRPVQDL

| | | | | | |
|---|---|---|---|---|---|
| MDEV | YPMSKEAKEG | IRPHIRRLLD | QGILVACQSP | WNTPLLPVRK | PGNYRPVQDL |
| PERV | YPLSREAREG | IWPHVQRLIQ | QGILVPVQSP | WNTPLLPVRK | PGNYRPVQDL |
| RfRV | YPMSAEARKG | IAPHINRLLE | AGILKPCHSA | WNTPLLPVKK | PGKYRPVQDL |
| WDSV | YPLPKDKTEG | LRPLISSLEN | QGILIKCHSP | CNTPIFPIKK | AGDYRMIHDL |
| WEHV1 | YPLPKEKVEG | LRPMIHSLLA | QGVLTECHSS | CNTPIFPIKK | PGEYRMIHDL |
| WEHV2 | YPLPKEKVNG | LRPMIESLLA | QGVLAECHSS | CNTPIFPIQK | PGQYRMIHDL |
| SSSV | YRLKYEQIEG | IRPTVEGLLG | ADCIYATSSP | WNTPILPVPK | ADKYRMVQDF |
| SnRV | YPVPDASHAS | IKETVEALLE | QGVLRKCNST | VNSAIWPVGK | PDGWRLTIDY |
| ALV | WPLPEGKLVA | LTQLVEKELQ | LGHIEPSLSC | WNTPVFVIRK | ASGYRLLHDL |
| LDV | WPLTAQKLDA | VQNIIQDLLK | DGRIIPSRSQ | WNSPIFVIQK | KDSFRMLHDL |
| MPMV | WPLTNDKLAA | AQQLVQEQLE | AGHITESSSP | WNTPIFVIKK | KSGWRLLQDL |
| SRV1 | WPLTSEKLAA | AQQLVQEQLE | AGHITESNSP | WNTPIFVIKK | KSGWRLLQDL |
| MMTV | WPLKQEKLQA | LQQLVTEQLQ | LGHLEESNSP | WNTPVFVIKK | KSGWRLLQDL |
| BLV | FPLNLERLQA | LQDLVHRSLE | AGYISPWDGP | GNNPVFPVRK | PNGWRFVHDL |
| HTLV1 | FPLNPERLQA | LQHLVRKALE | AGHIEPYTGP | GNNPVFPVKK | ANGWRFIHDL |
| STLV2 | FPFKPERLQA | LTDLVSKALE | ASYIEPYSGP | GNNPVFPVKK | PNGWRFIHDL |
| VMV | WPLTQEKLEG | LKEIVDRLEK | EGKVGRAHWT | CNTPIFCIKK | KSGWRMLIDF |
| EIAV | WPLTKEKLEG | AKEIVQRLLS | EGKISEANNP | YNSPIFVIKK | RSGWRLLQDL |
| FIV | WPLTNEKIEA | LTEIVERLER | EGKVKRANNP | WNTPVFAIKK | KSGWRMLIDF |
| HIV-1 | WPLTEEKIRA | LTDICTEMEK | EGKISKIENP | YNTPVFAIKK | KDTWRKLVDF |

| | | | | |
|---|---|---|---|---|
| KALNRRTVAN | RASLISCIDL | ANGFFSLRLT | RGSQGKTAFT | HKGRSYVWER |
| KALNRRTVAN | RASLISCIDL | ANGFFSLRLA | RESQGKTAFT | HKGKSYVWQR |
| KALNRRTIAN | RASLISCIDL | ANGFFSLRLA | RQSQGKTAFT | HKGKSYVWQR |
| KALNKQTVPD | TRYLISKIDL | ANGFWSVPLK | PESRARTAFT | FENKQYVYNV |
| REVNKVTPLV | ATQNCSTLDL | ANGFWAHPIK | PEDYWITAFT | WGGKTYCWTV |
| RAVNKVTPAI | ATQNCTTLDL | ANGFWAHPIQ | ESDQWITSFT | WNGKSYVWTT |
| REVNKTIPLI | AAQNQTTLDL | ANGFWAHPIT | PNSYWLTAFT | WQGKQHCWTR |
| REVNKTIPLT | AAQNQTTLDL | ANGFWAHPIT | PDSYWLTAFT | WQGKQYCWTR |
| REVNKTIPLI | AAQNQTTLVL | ANGFWAHPIT | PESYWLTAFI | WQGKQYCWTR |
| REVNKTIPLI | AAQNQTTLDL | TNGFWAHPIT | PESYWLTAFT | WQGKQYCWTR |
| REVNKTIPLI | AAQNQTTLDL | SNGFWAHSIT | PESYWLTAFT | WLGQQYCWTR |
| RAVNKTVPLI | GAQNQSTIDL | SNGFWAHPIT | KDSQWITAFT | WEGKQHVWTR |
| RAVNKTIPLI | AAQNQSTIDL | SNGFWAHPIT | EDSQWITAFT | WEGKQHVWTR |
| RAVNKTIPLI | AAQNQSTIDL | SNGFWAHPID | QDSQWITAFT | WEGKQYVWTR |
| RAVNKTVPAI | GAQNCTTLDL | SNGFWSHPIT | PESYWLTAFT | WQGSQYVWTR |
| RAVNKVTPLI | AVQNQTTIDL | SNGFWAHPIV | PEDYWITAFT | WQGKQYCWTV |
| RAVNKVIPLI | AVQNQTTLDL | SNGFWAHPIR | KESYWLMAFT | WEGKQLVWTR |
| HTLNRVSPSF | NVQNLTTLDL | SNGFWTHPIR | EQDRYLTAFS | WLGTQYWWTR |
| HALNRVSTSC | NVQNLTSLHL | SNGFWAHPIR | EQDRYLTAFS | WQGTQYSWTH |
| RALNKVSPSF | TVQNLTTLDL | SNGFWAHPIR | EEDRYLTGFT | WQGIQYCWTR |
| RALNKVSPSF | TVQNLTTLDL | SNGFWAHPIR | EEDRYLTGFT | WQGIQYCWTR |
| RALNKVSPSF | TVQNLTTLDL | SNGFWAHPIR | EEDRYLTGFT | WQGIQYCWTR |
| RALNKVSPSF | TVQNLTTLDL | SNGFWAHPIR | EEDRYLTGFT | WQGIQYCWTR |

RALNKVSPSF TVQNLTTLDL SNGFWAHPIR EEDRYLTGFT WQGIQYCWTR
RALNKVSPSF TVQNLTTLDL SNGFWAHPIR EEDRYLTGFT WQGIQYCWTR
RALNKVSPSF TVQNLTTLDL SNGFWAHPIR EEDRYLTGFT WQGNQYCWTR
RALNKVSPSF TVQNLTTLDL SNGFWAHPIR EEDRYLTGFT WQGNQYCWTR
RALNRVSPSF TVQNLTTLDL SNGFWAHPIR EEDRYLTGFT WQGNQYCWTR
RAVNRVSASF TVPNLTTLDL SNGFWAHPIR EEDRYLTSFT WQGNQYCWTR
RALNRVSPSF NVQNLTTLDL SNGFGAHPSR EQDRYLTAFS GQGTQYCWTT
HSLNRVSPSF TVQNLTTIDL SNGFWAHPIR KEERYLTNFT WQGGQYCWTH
RALNRVSASF NVQKLTTLDL YNGFWAYPIR EKDRYLTAFS WQGTQ-CWIR
RALNRVSASF NVQKLTTLDL YNGFWAYPIR EKDRYLTAFS WQGTQ-CWIR
CALNRVSPSF SV-NLTILDL SNGFWAHPFR EQDMYLTAFT WQGTQYCWIR
RALNKHSEPV RAQNQSSVDL ANGFWSHPIR EEDWPKMAFT WCGFQYLWTR
RHLNSHTRSF AIHNTTTLGI SN-FFTQNLA PESQDWSTFT CLGIQYKFWR
REVNKRVEDI HPTVPTVLDL KDAFFCLRLH SESQLLFAFE WRSGQLTWTR
REVNKRVEDI HPTVPTVLDL KDAFFCLRLH PTSQSLFAFE WRSGQLTWTR
REVNKRVLDI HPTVPTVLDL KDAFFCLRLH PKSQLLFAFE WRTGQLTWTR
REVNKRVQDI HPTVPTVLDL KDAFFCLRLH PTSQPLFAFE WRTGQLTWTR
REVNKRVEDI HPTVPTTLDL KDAFFSIALA PSSQHIFAFE WNPGQLTWTR
RAINNIVAPL TAVVATVIDL SNAFFSVPIH KDSQYLFAFT FEGHQYTWTV
RAINEIVAPL TAVVATVIDL SNAFFSVPIH PDSQYLFAFT FEGRQYTWTV
RAINEIVAPL TAVVATVIDL SNAFFSVPVH PDSQYLFAFT FEGRQYTWTV
RAVNDVTTGV DLPVPTVVDL ANAFFSIPLH VDSQPLFAFT YDNQQYTYSV
RPLNSAVSCP YPTVASSLDI SNGFWSIRLE EECQYLFAFT FDTQQYTWTR
RAVNAKLVPF GAVQQMVLDL KDCFFSIPLA EQDREAFAFT LPARRFQWKV
RAVNALIKDW GALQPIAMDI SDCFFSIPLA ERDSERFAFT IPAKRYQWTV
RAVNATMVLM GALQPIIIDL KDCFFSIPLH PSDQKRFAFS LPMQRFQWKV
RAVNATMVLM GALQPIIIDL KDCFFSIPLH PSDQKRFAFS LPMQRFQWKV
RAVNATMHDM GALQPIIIDL QDCFFNIKLH PEDCKRFAFS VPYQRFQWKV
RATNALTKPI PALSPICLDL KDAFFQIPVE DRFRSYFAFT LPHRRFAWRV
RATNSLTIDL SSSSPQTIDL KDAFFQIPLP KQFQPYFAFT VPGTRYAWRV
RATNAITTTL ASPSPQTIDL TDAFFQIPLP KQFQPYFAFT IPGARYAWTV
RELNKQTEDL AEAQLTILDI GDAYFTIPLY EPYRQYTCFT MLCVRYYWKV
RELNKTVQVG TEISRTVLDI GDAYFTIPLD PEFRPYTAFT IPDKRYVWKC
RELNKLTEKG AEVQLTVLDI GDAYFTIPLD PDYAPYTAFT LPGRRFVWCS
RELNKRTQDF WEVQLTVLGV GDAYFSVPLD PGFRKYTAFT IPGIRYQYNV

LPQGYKNSPN VFQAAVMDVL DGL--I--YI DDVFIADDTE EQHLDRLQKI
LPQGYRNSPN VFQSAVLEIL EDV--V--YI DDVFIADDTE EEHLKRLEEV
LPQGYKNSPN VFQSAVMEVL GDV--V--YI DDVFIADDTE EEHLERLRKV
LPQGFKNSPN AFQAIMMEIL KGL---PVYI DDILIVTNDE EEHLKILDET
LPQGFLNSPA LFTADVVDIL KDIPNVQVYV DDVYVSSATE QEHLDILETI
LPQGFLNSPA LFTADVVDLL KDIPNVEVYV DDVYFSNDTE EEHLKTMDLL
LPQGFLNSPA LFTADVVDLM KHIPNVQVYV DDLYLSHDDP QEHLQVLQQV
LPQGFLNSPA LFTADAVDLL KEVPNVQVYV DDIYLSHDNP HEHIQQLEKV

LPQGFLNSPA LFTADVVDLL KEISNVQAYV DDIYLSHDDP QEHLDQLEKV
LPQGFLNSPA LFTADVVDLL KEIPNVQAYV DDIYISHDDP QEHLEQLEKI
LPQGFLNSPA LFTADVVDLL KEVPNVQVYV DDIYISHDDP REHLEQLEKV
LPQGFLNSPA LFTADVVDLL KNIPGISVYV DDIYFSTETV SEHLKILEKV
LPQGFLNSPA LFTADVVDIL KEVPGVSVYV DDIYISSPTM EEHFQVLDSI
LPQGFLNSPA LFTADVVDLL KEIPNVNVYV DDIYVSTETI NQHFQVLDKI
LPQGFLNSPA LFTADVVDLC KHIPNVSAYV DDIYVSNDTA EEHLRTLEQL
LPQGFLNSPG LFTGDVVDLL QGIPNVEVYV DDVYISHDSE KEHLEYLDIL
LPQGFINSPA LFTANIVDIL KEIPDVEVYV NDIYFSNVTE EQHLITLKQV
LPQGYLNSPA FFSADVIEFL KNIPGGHSYM DDIYFTNEDL DKHLATLKQI
LPQGYLNSPA LFSADVIQLL RNIPGVHSYM YDIYFTNEDL DQHLATLKQI
LPQGYLNSPA IFSADVIQLL KDIPGVNSYM DDIYFTNENL DQHLATLKQV
LPQGYLNSPA IFSADVIQLL KDIPGVNSYM DDIYFTNENL DQHLATLKQV
LPQGYLNSPA IFSADVIQLL KDIPGVNSYM DDIYFTNENL DQHLATLKQV
LPQGYLNSPA IFSADVIQLL KDIPGVNSYM DDIYFTNENL DQHLATLKQV
LPQGYLNSPA IFSADVIQLL KDIPGVNSYM DDIYFTNENL DQHLATLKQV
LPQGYLNSPA IFSADVIQLL KDIPGVNSYM DDIYFTNENL DQHLATLKQV
LPQGYLNSPA IFSADVIQLL KDIPGVNSYM DDIYFTNENL DQHLATLKQV
LPQGYLNSPA IFSADVIQLL KDIPGVNSYM DDIYFTNENL DQHLATLKQV
LPQGYLNSPA IFSADVIQLL KDIPGVNSYM DDIYFTNENL DQHLATLKQV
LPQGYLNSPA IFSADVIHLL KGIPGVNSYM DDIYFTNENL DQHLATLKQV
VLLDYLNSPA LFSADVIELL KNIPGVHSYM DDVYFTNEDL DKHLATLKQI
FPQGYLNSPA IFSADVIQLI KDIPGVNSYM DDIYFANENL DQHLATLKQV
LPQGYLNSPA LFSADVIQLL RNIPGVHSYM DDIYFTNEDL DQHLATLKQT
LPQGYLNSPA LFSADVIQLL RNIPGVHSYM DDIYFTNEDL DQHLATLKQT
LPQGYLNSPV LFSADVIALL KNIPGVHSYM DDIDFTNKDL DKYLAILKQI
LPQGFLNSPA LFSADVISLV GQLPGVYCYV DDIYLTHDTE EEHLKILDQV
LPMGYKNSLG VFAVRIIELL QLVPDAVSYI DYIYVTDDEM QQHLARVDRI
LPQGFKNSPT LFDEALHSDL ADFPALVLYV DDLLLAAATR TECLEGTKAL
LPQGFKNSPT LFDEALHRDL ADFPDLILYV DDLLLAATSE LDCQQGTRAL
LPQGFKNSPT LFDEALHRDL APFPQLTLYV DDLLIAAASK ELCQQGTERL
LPQGFKNSPT IFDEALHRDL ANFPQVTLYV DDLLLAGATK QDCLEGTKAL
LPQGFKNSPT LFNEALNQDL DSFNSVTLYV DDLLLAAPSE AECRQATGDL
LPQGFIHSPT LFSQALYQSL HKISSEICYM DDVLIASKDR DTNLKDTAVM
LPQGFIHSPT LFSQALFSSL SKITSEICYM DDVLIASRDE ETNYKDTATM
LPQGFIHSPT LFSQALFSSL SKITSEVCYM DDVLIASTTK DINIKDTVTL
LPQGYRCSPG IFNRVLKDHL KEIDGVVLYV DDLLLAACSS DQCLAATKIL
LPQGFHASPG IFHQALYNGL ASCQGCKLYV DDILLMSEDR DHHLRSLAIL
LPQGMTCSPT ICQLIVGQVL EPLPSLCMYM DDLLLAASSH DRLEAAGEEV
LPQGMKNSPY ICQQVVAEVI RPIRDAVIYM DDILIAAAEE RQTEVIFEAV
LPQGMANSPT LCQKYVATAI HKVKQMYIYM DDILIAGKDG QQVLQCFDQL
LPQRMANSPT LCQKYVATAI HKVKQMYIYM DDILIAGKDG QQVLQCFDQL
LPQGMKNSPT LCQKFVDKAI LTVQDSYIYM DDILLAHPSR SIVDEILTSM
LPQGFINSPA LFERALQEPL RQVSQSLLYM DDILIASPTE EQRSQCYQAL

LPQGFKNSPT LFEMQLAHIL QPIPQCTIYM DDILLASPSH ADLQLLSEAT
LPQGFKNSPT LFEQQLAAVL SPIPMSTIYM DDILLASPTQ EELQQLSKMT
LPQGWKLSPA VYQFTMQKIL RGWPMIQFYM DDIYIGSDGL EEHRGIVNEL
LPQGFVLSPY IYQKTLQEIL QPFPEVQLYM DDLFMGSNSK KQHKELIIEL
LPQGWILSPL IYQSTLDNII QPFPQLDIYM DDIYIGSNSK KEHKEKVEEL
LPQGWKGSPA IFQCSMTKIL EPYPEVIIYM DDLYVGSDEI EQHRAKIEGL

VERLTAAGLK LNLKKCQVDY LGFQVSKIEQ IEPPSSLNEL QKVLGLCNYV
VKRISAAGLK LNLAKCQVNY LGFQVTKIQQ IQPPRCENDL QKILGLCNYV
VENLTKAGLK LNLKKCQVNY LGFQVTKLTN IQPPQSENDL QKILGLCNYV
LSRLTKEGFR LNYKKMEVEF LGFQITKLKE LRGITSLKQL QSMLGTLNFV
FNRLSTAGYI VSLKKSKVEF LGFSISKLMD LQPPTTLRQL QSILGLINFA
FQKLQTAGYI VSLKKSKVDF LGFQITKLLD ITPPNTLKQL QSILGLLNFA
LHILHDAGYV VSLKKSAVEF LGFNITKLLN ISPPQNLKQL QSILGLMNFA
FQILLQAGYV VSLKKSEVEF LGFNITKLLN VTPPKDLKQL QSILGLLNFA
FQILLQAGYV VSLKKSEVEF LGFNITKLLD ITPPKDLKQL QSILGLLNFA
FSILLNAGYV VSLKKSEVEF LGFNITKLLN ITPPKDLKQL QSILGLLNFA
FSLLLNAGYV VSLKKSEVEF LGFNITKLLN ITPPRDLKQL QSILGLLNFA
FKILLEAGYI VSLKKSAVTF LGFSITKIQN ITSPRTLKEL QSILGLFNFA
FRKLLETGYI VSLKKSAVNF LGFVISRLQE ITPPTTLKQL QSILGFLNFA
FQKLLQAGYV VSLKKSNVTF LGFTISKLRN ISPPNSLKQL QSILGLLNFA
FRTLMSAGYI VSLKKSKVDF LGFEITKLVN IQPPSSLKQL QSILGFLNFT
FNRLKEAGYI ISLKKSNVDF LGFQITKLEN ITAPTTLKQL QSILGLLNFA
LKILLKSGYI VSLKKSEVTF LSFNITKLLN ISAPKTLKQL QSILGLLNFA
VTILGEVGYI INLKKSKEKF LGFQLTKLLT LQPPKFFKEL QSVLGFLNIG
VTVLGEAGY- INLKKSKVKF LGFLL-KLLT LQPPKTLKEL QSVLGFLNVA
VTVLGEAGYI INLKKSKVKF LGFLLTKLLT LQPPKTLKEL QSVLGFLNVA
VTVLGEAGYI INLKKSKVKF LGFLLTKLLT LQPPKTLKEL QSVLGFLNVA
VTVLGEAGYI INLKKSKVKF LGFLLTKLLT LQPPKTLKEL QSVLGFLNVA
VTVLGEAGYI INLKKSKVKF LGFLLTKLLT LQPPKTLKEL QSVLGFLNVA
VTVLGEAGYI INLKKSKVKF LGFLLTKLLT LQPPKTLKEL QSVLGFLNVA
VTVLGEAGYI INLKKSKVKF LGFLLTKLLT LQPPKTLKEL QSVLGFLNVA
VTVLGEAGYI INLKKSKVKF LGFLLTKLLT LQPPKTLKEL QSVLGFLNVA
VTVLGEAGYI INLKKSKVKF LGFLLTKLLT LQPPKTLKEL QSVLGFLNVA
VTVLGEAGYI INLKKSKVKF LGFLLTKLLT LQPPKTLKEL QSVLGFLNVA
VTVLGEAGYI INLKKSKVKF LGFLLTKLLT LQPPKTLKEL QSVLGFLNVA
VTVLGEAGYI INLKKSKVKF LGFLLTKLLT LPRPKTLKEL QSVLGFFNVA
VTVLGEAGYI INLKKSKVKF LDFLLIKLLT LQPPKTLKEL QSVLGFLNVA
VTVL-EAGYI INLKKTKVKF LGFLLTKLLT LKPPKSLKEL QSILGFLNVA
VTVL-EAGYI INLKKTKVKF LGFLLTKLLT LKPPKSLKEL QSILGFLNVA
VAVLGEVGYI INLKKIKISW IS------IN FAAASNFKGI AVSFRFFNIG
LEILIKAGYV INIKKSKVDF LGFSLSKLAA IKPPQTLRQL QSVMGLLNFS
IIHFSASGYK INFKKSKVLF LGYKLNRCAV LQDPNTLTKL HSLLNFLNFG
LETLGNKGYR ASAKKAQVTY LGYSLKAILS IPVPKNSRQV REFLGTAGYC

LQTLGDLGYR ASAKKAQVKY LGYLLKTVMG QPTPKTPRQL REFLGTAGFC
LTELGNLGYR VSAKKAQVIY LGYTLRTVMM IPPPTTPRQV REFLGTAGFC
LLELSDLGYR ASAKKAQVTY LGYSLRTVVQ IPAPTTAKQV REFLGTAGFC
LQELGQLGYR ASAKKAQVTY LGYKLKTILR LPVPTSAREV REFLGTTGYC
LQHLASEGHK VSKKKLQVVY LGQLLTTVSQ FQQPTTIRQI RAFLGLVGYC
LHHLADEGHK VSKKKLQVVY LGQLISTVSQ FAAPTTVRQV RAFLGLAGYC
LHHLADEGHK VSQKKLQVIY LGQLLSTVSQ FSSPTTVRQV RAFLGLAGYC
LDFLATKGYK VKRSKVQVLF LGREVSSILH HPRPTTVSGL LSFLGLTGYS
LQGLKDLGVK INPKKSHVQY LGVNVGLIRT LDIPLTVQGL RSALGLFNFC
ISTLERAGFT ISPDKVQVQY LGYKLGGLVA EPRIATLWDV QKLVGSLQWL
KTTCHEKGLK INEAKTQVSY LGWRVMTVQL PETVTNLVQL QRLLGIIQWL
KQELTAAGLH IAPEKVQYTY LGFELN-VIR KDKLQTLNDF QKLLGDINWL
KQELTIAGLH IAPEKIQYTY LGFELN-VIR KDKLQTLNDF QKLLGDINWL
IQALNKHGLV VSTEKIQLKY LGTHIQ-QIR TDKLRTLNDF QKLLGNINWI
AARLRDLGFQ VASEKTRVPF LGQMVHPTLQ ISSPISLHQL QAVLGDLQWV
MASLISHGLP VSENKTQIKF LGQIISPKVP IRSRWALPEL QALLGEIQWV
LQALVTHGLP VSQEKTQIRF LGQVISPTIP IKSQWTLTEL QTVLGEIQWV
ASYIAQYGFM LPEDKRQAKW LGFELHTLPE ITEPITLNKL QKLVGDLVWR
RAILLEKGFE TPDDKLQYSW LGYQLCQLDM VK-NPTLNDV QKLMGNITWM
RKLLLWWGFE TPEDKLQYTW MGYELHQL-D IPEQPTLNEL QKLAGKINWA
RDHLLKWGFT TPDKKHQFLW MGYELHQLPD KE-SWTVNDI QKLVGKLNWA

RDYVPGYQKP LKKPDTAWTA ADQSNLEELR QAIQGARLEP RSPTAKLVAE
RDHVPNYQKP LKKEAGKWTP KNQDDLDKLK EEIQKARLEP RNLTSRLVAE
RDHVPNYQKP LRKSEKRWTA ANQQNLEELR RAIQAARLEP RSLSERLVAE
RDLIPGYAKP LRGGV--WNE KLSEIKAKLI DEALNSRLVR RDETTPLVVK
RNFLPNFAAP LPKAKGQWTM DHTTQLKTII QALNSTNLEE RRPDVDLIMK
RNFIPNYSTP LPLAKGIWET KHTAILQKII KELNASNLEQ RKPDVELIVK
RNFIPNYAKP FSTAKSNWND ELTSQLQELI TLLNQANLEE RKPTTRLIIK
RNFIPNFAQT LASSKGKWTE DNTKQLNKVI EALNTANLEE RLPDQRLVIK
RNFILNFAKP LSSAKGKWSN ENTVQLQTII KALNNANLEE RIPEKRLIIK
RNFIPNYSKP LANANGKWTE DNSNQLQHII SVLNQANLEE RNPETRLIIK
RNFIPNFSKP LATANGKWTT DNSQQLQNII SMLNSANLEE RNPEVRLIMK
RNFVPNFSKP LSTAEGNWTS EHTRYLEEIV SALNHANLEQ RDNESPLVVK
RNFVPNFSQP LSTASGNWTA EHTLRLNELI SALNHANLEQ RRGDSPLVVK
RNFIPNFSKP LSTAQGQWEP KHSQALNNLI IALNHANLEQ RNGEVPLVIK
RNFVPNYSKP LATAQGNWTL ENTEQLNQVI SALNAANLSE RKTGVPLVVK
RNFIPDFTAP LPKSTKNWQI EHSTTLETLI TKLNGAYLQG RKGDKTLIMK
HNVITDFAKP LSRAEGQWME KEGMALQEII KKLNNAYLEN RDIQKPLIIK
RILVPDYAKP LPESKGNWTL EAQQTLDNLI TLINQAVLNT RNSTVSLEIL
RILVPDYTKP LPASKGN-TL EAQQTLDNLI TLINQAVLNT RNSTVSLEIL
RILVPDYAKP LPASKGNWTM EAQQALDELI VLINQAALNT RNNTASLEII
RILVPDYAKP LPASKGNWTL EAQQALDELI VLINQAALNT RNNTASLEII
RILVPDYAKP LPASKGNWTL EAQQALDELI VLINQAALNT RNNTASLEII

RILVPDYAKP LPASKGNWTL EAQQALDELI VLINQAALNT RNNAAALEII
RILVPDYAKP LPASKGNWTM EAQQALDELI VLINQAALNT RNNAAALEII
RILVPDYAKP LPASKGNWTM EAQQALDELI VLINQAALNT RNNAAALEII
RILVPDYAKP LPASKGNWTM EAQQALDELI VLINQAALNT RNNAASLEII
RILVPDYAKP LPASKGNWTM EAQQALDELI VLINQAALNT RNNAASLEII
RILVPDYAKP LPASKGNWTL EAQQALDELI VLINQAALNT RNNAAALEII
RILVPDYAKP LPASKGNWTL EAQQALDELI VLINQAALNT RNNTASLEII
RILVPDYAKP LPPSKGNWTL EAQQTLDNLI TLINQAVLNT RNSTVSLKIL
RILVTDYAKP LPAIKGNWT- EAQQALDK-I VLIKQATLNT RNNTASLEI-
RILVPDYAKH LPASKGNWTL EAQQILDSLI TLINQAVLNT RNSTVSLEIL
RILVPDYAKH LPASKGNWTL EAQQILDSLI TLINQAVLNT RNSTVSLEIL
PLCSENQTNS L-ASKGNRTL ESQQALDNLI TSINQAALNT RNNTVSLEIL
RLFVKGFSKP LTVKP--FTP QAHEALTKLL SAIQTANLTN RDNTRALVIR
RRYIPEYAKP LEHDY--WTD AHTDILRSLQ RDLLELHLHT RDNTTNLEIR
RLWIPGFAAP LRPGT--WGT EQQLAFEDIK KALLSSALGL PDITKPFELF
RLWIPGFAAP LKTGT--WGP DQQKAYQEIK QALLTAALGL PDLTKPFELF
RLWIPGFAAP LREGI--WKE EHQRAFEAIK SSLMTAALAL PDLTKSFVLY
RLWIPGFAAP LKEKG--WAP EHQKAFDAIK KALLSAALAL PDVTKPFTLY
RLWILGYAKP LKDKV--WGS DQQKAYDELK VALLRAALAL PDPLKPFTLF
RHWIPEFSKF LKKDTAELDD QQVEAFNKLK HAITTAVLVV PDPAKPFQLY
RHWIADYSKH LKKEVIELTG EQLRAFEQLK STLLSAVLAI PDYRKDFELY
RHWIPDYSKY LKKEVEELTE KQTEAFQKLK QSLITAVLAI PDYKKDFELY
RTHIPDYTEP LRQAGHAWTP EATCAFGLLK TDLSVAALAA PDYSKPFHLD
RAWIPEFSQS LKGDCTDWTE DNLNKFKLLK DEVASAVLGL PDPTLPFRHL
RPALGIPPGP FRGS-NEWNL DMKMAWREIV QLSTTAL-ER WDPALPLEGA
KPILSLRPQV FKGKTEELTE EAQQELRNIQ QRLQASV-GR WRSDLPITLF
RPYLKLTTKP LKGDSNSLSK EALASLEKVE TAIAEQV-TH INYSLPLIFL
RPYLKLTTKP LKGDSNSLSK EALALLDKVE TAIAEQV-TH INYSLPLMFL
RPFLKLTTKP LNGDSISLTP EACKALQLMN ERLSTAV-KR LDLSQPWSLC
SRGTPTTRQL LKGIDRALSP EQLQGIAELR QALSHNR-SR YNEQEPLLAY
SKGTPTLRHS LQRHTRDLNP SQVQSLVQLR QALSQNR-SR LVQTLPLLGA
SKGTPILRQC LRGYQRALQK QQLHALHAIQ QALQHNR-SR LNPALPILGL
QSLI---GPN IEGDRQSIES IHVREWEACR QKLKEMG-NY YDEEKDIYGQ
SSGIPGLTKH IKGCLNQWTE EAQKELEENN EKIKNAGLQY YNPEEEMLCE
SQAIPDLSKA LRGNQNSWTK EARLEVQKAK KAIEEQQLGY YDPSKELYAK
SQIFPGIRKN LRGAKTDLTA EAELELAENR EILKEPHGVY YDPSKDLIAE

ISCDDSATVK VSNE-GLVTL WSLTSVEVKF PPEEKELYWS SLKDLAQGWG
IFCEEDAMVK VSND-GLITL WSLTSVEKKY PQEEKELYWG SLKDLAQGWG
ISCEDDAMIK VSNE-GLVTL WSLTSVEKKY PQEEKELYWG VLKDLAQGWG
IANTDEIELI IFNEKHPVCF ISKPANQRKL ETGGDILSLI TLKCLALDWA
VHISTAGYIR FYNHQKPIAY NNFTSTELKF TPTEKIMGLL KALDLSLGWL
VHVSTAGYIK FANKIKPIAY HNFSKTELKF TITEKVMALL KAFDLAMGWL
VNSSHAGYIR YYNEKKPILY INFSKAEEKF SMLEKLLALI KAVDLAMGWI

```
VNTSSAGYVR YYNEKKPIMY LNFSKAELKF SMLEKLLALI KAMDLAMGWI
VNTSSAGYVR YYNEKKPIMY LNFSKAELKF TLLEKLLALI KAMDLAMGWI
VNSSSAGYIR YYNEKRPIMY VNFSKAEAKF TQTEKLLGLI KAMDLAMGWI
VNTSSAGYIR FYNEKRPIMY LNYTKAEVKF TNTEKLLGLI KALDLGMGWI
LNASKTGYIR YYNKQKPIAY ASFTNTELKF TPLEKLLALI KAIDLALGWI
VNASKTGYIR YYNDLIPIAY ASFSTAELKF TPLEKLLALL KGIDLALGWV
INASTTGYIR FYNKKRPIAY ASFNHTEQKF TPVEKLLAII KGIDLAIGWL
SNCSTAGYIR FYNQRKPIQY VNFSKTELKF TPLEKQLAIL KGLDLAGGWL
VNASTTGYIR YYNEKKPISY VSFSKTELKF TELEKLLGLL KALDLSMGWL
LNSSTAGYIR MYNKKKPIQY VNFTPAEIKF KPTEKLLAII KGLDLSQGWI
VGATKGGFTS YFNESKPLLY ISFLNAEQKF LPIERNLAIL KGKDLAQGWV
VGATKREFAS YFNESKPLQY ISFSNAEQKV LPIERNLATL KGKDLAQGWV
VRASRGGMAS YYNDSKPLQY ISFSNAEQKF LPTERNLAIL KGKDLAQGWV
VRASRGGMAS YYNDSKPLQY ISFSNAEQKF LPTERNLAIL KGKDIAQGWV
VRASRGGMAS YYNDSKPLQY ISFSNAEQKF LPTERNLAIL KGKDIAQGWV
VRASRGGMAS YYNDSKPLQY ISFSNAEQKF LPTERNLAIL KGKDLAQGWV
VRASRGGMAS YYNDSKPLQY ISFSNAEQKF LPTERNLAIL KGKDIAQGWV
VRASRGGMAS YYNDSKPLQY ISFSNAEQKF LPTERNLAIL KGKDIAQGWV
VRAS-GGMAS YYNDSKPLQY ISFSNAEQKF LPTERNLAIL KGKDLAQGWV
VRAS-GGMAS YYNDSKPLQY ISFSNAEQKF LPTERNLAIL KGKDLAQGWV
VRASRGGMAS YYNDSKPLQY ISFSNAEQKF LPTERNLAIL KGKDIAQGWV
VRVSQGGMAS YYNDSKPLQY ISFSNTEQKF LPTERNLAIL KGKDLAQGWV
VGDTKLGFAS YFNESKPLQY ISFSNAEQKF LPIGRNLPIL KGKDLAQGWV
VRVSQGGFAS YYNDSKPLQY ISFSNAEQKF LPIERNLAIL KAKDLAHSWV
VGAIK-GFAS YFNESKPLQY KSF------- ---------- ----------
VGAIK-GFAS YFNESKPLQY KSF------- ---------- ----------
VSASQGGIAS YFNESKPLQY ISFSNAEQNC LPIESNVAIL KGKDITPGWV
IVTSRSAVAA YYNVKLPIQY VSFSQPEMKF SKIEKVCALI KGRVLAHEWD
VSPSTGFTYM TFNETVPMAY KSYSSVEMRF ASTDFFFAIL KERPLAQGWI
IDENGFAKGV LVQKKRPVAY LSLDTVASGW PPCLRMVLVK DAGKLTLGMT
VDEKGYAKGV LTQKRRPVAY LSLDPVAAGW PPCLRMVLTK DAGKLTMGMT
VDERGIARGV LTQAKRPVAY LSLDPVASGW PTCLKAILIK DADKLTMGMT
VDERGVARGV LTQTRRPVAY LSLDPVASGW PVCLKAILVK DADKLTLGMT
VDERGIAKGV LMQRKRPVAY LSLDPVAAGW PPCLRIIMVK DADKLTFGLT
TSHSHASIAV LTQKTRPIAF LSFDAIESGL PPCLKACSLT QADSFILGFS
TSHTHVAVAV LAQKTRPIAF LSLDAIEQGL PPCLRACNLT QADSFLLGFS
TSHTHVAIAV LSQKSRPIAY LSLDAIERGL PPCLRACNLT QADSFLLGFS
VSEKGFASAI LFQKRRVLMY HSLDHIETGQ TTCSRYVAIE KTAHLVMCKN
IGIRGHFLCS LVQKWHVLGF YSMTPVESNL GICEQYAAIS ACNLVSFGLA
VARCQGAIGV LGQGPKPCLW LFSTQPTKAF TAWLEVLKLR ASAVRTFG--
VCTTAGGIGL IAQQIQPLYW LSSTRLTRAW TSASRVLQGS QICAQYWG--
IFNTLTPTGL FWQD-NPIMW IHPASPKKVL LPYYDAILGR DHSKKYFG--
IFNTLTPTGL FWQN-NPIMW VHPASPKKVL LPYYDAILGR DHSKKYFG--
ILKTYTPTAC LWQD-GVVEW IHPHISPKVI TPYDIFCKGR HRSKELFS--
```

VHLTAGSTLV LFQKQFPLAY FQTPLTDNQA SPWGLLLYLQ TQALSSYAPG
IMLTTGTTTV VFQSQWPLVW LHAPLPHTSQ CPWGQLLLLD KYTLQSYGGE
ISLSSGTTSV LFQARWPLVW LHTPHPPTSL CPWGHLLTLD KYSLQHYGSG
LDWGKAIEYI VFQE-GKPLW VNVHSIKN-- -SQAQQIKLT QEVIIRTG--
VEITYEATYV IKQS-QGILW AGIMKANKGW -STVKNLHVA TESITRVG-V
LSLVHQISYQ VYQK-EKILW YGMSRQKKKA ENTCDIAKIR EESIIRIG--
IQKQDQWTYQ IYQE-YKNLK TGYARRKSAH TNDVKQLKIA MESIVIWG--

RWEDILLDPD LEIATQ-EEY TLFTDGSRKG AYWGFIGSAQ AGEVTAVLEG
RWEDILLDPD LEIPTD-KTY VLFTDGSRKG AYWGYIGSAQ AGEVTAVLEG
RWEDILLDPE LEIPPP-GQP TIYTDGSRKG AYWGFIGSAQ AGEVTAVLEG
KWTVILNDTQ FQFPKPI--- CYYTDGTTEQ --WGWILSAQ AAEVKAFRKA
KWQTYFEDPR IKFPQPLLHY IFYTDGSAIR AGMGIIHTAQ YAEIAAFEFA
KWQTYFEDPR LIFPQPLSEY VFYTDGSSIK AGMGIIHTAQ FAEIAAFEFA
TWMTYLEDPR ITFPSHPSQY VFYTDGSAIK AGMGVVHTAQ LAEVAAVEFA
TWMTYLEDPR IQFPDHPSQY VFCTDGSAIK AGMGIVHTAQ MAEIAAVEFA
TWMTYLEDPR IQFPDHPSQY VFYTDGSAIR AGMGIVHTAQ MAEIAAVEFA
TWMTYLEDPR IQFPNHPSEF VFYTDGSAIK AGMGIAHTAQ LAEIAAVEFA
TWMSYLEDPR IQFPTHPSEF VFYTDGSAIK AGMGIAHTAQ LAEVAAVEFA
TWLSYLEDPR ITFPEPIIEY VFYTDGSAIR SGMGIVHTAQ YAEISAVEFA
TWLSYLEDPR ITFPAPITEY VFYTDGSAIK AGMGIVHTAQ YAEISAVEFA
SWLSYIEDPR FLFPPPLHQY VFYTDGSSIK SGMGIVHTAQ YAEIAAVEFA
TWITYLEDPR FHFPEPLKEY VFYTDGSAIK AGMGTVHTAQ YAEIAACEFA
SWLSYLEDPR IRFPAP-SNF IFYTDGSAIT AGMGIVHTAQ FAEIAAFEFA
TWMTHFKNPQ LIFPQPLTSY VYYTDGSAIK AGIGIVHTAQ YAEISALEFA
QWMSHFENIE FLTPDPLLEY VIYTDGSAMK VGAVVINTAQ YAEMMAVHKA
QLMSHFENIE FPTPDPLHEY VINTDGSAM- -GAVVINTAQ YAELMAVHKA
QWMSHFENIE FPTPDPLHEY VIYTDESAM- ---------- --------KA
QWMSHFENIE FPTPDPLHEY VIYTDESAM- ---------- ----------
QWMSHFENIE FPTPDPLHEY VIYTDESAM- ---------- --------KA
QWMSHFENIE FPTPDPLHEY VIYTDESAM- ---------- -----SVHKA
QWMSHFENIE FPTPDPLHEY VIYTDESAM- ---------- ----------
QWMSHFENIE FPTPDPLHEY VIYTDESAM- ---------- ----------
QWMSHFENIE FPTPDPLHEY VIYTDESAM- ---------- ----------
QWMSHFENIE FPTPDPLHEY VIYTDESAM- ---------- --------KA
QWMSHFENIE FPTPDPLHEY VIYTDESAM- ---------- ----------
QWMSHFENIE FPTPDPLHEY VIYTDESAP- ---------- --QLCPVNKA
QWMSHFENIE FSTPDPLHEY LVYTDGSAMK ADAVVINTAQ YAELMTVHKA
QWMSHFENIE FTTPNPLHEY VIYTDGSAMK AGAVVINTAQ YAELMGVHKA
---------- -----PLHVY VIYTDGSAMK AGAVVINTAQ YAELMVVPKA
---------K CRTKLPLHVY VIYTDGSAMK AGAVVINTAQ YAELMVVPKA
QWMSHFENKE LPFPDPLHEY VIYTDGLAMK AGAVVINTAQ YAKLMAVHKA
LWCTLFSDQR LTFPPPLEQY VVYSDGSAKR AGIGIVASAQ YAEVMALLDA
QWAASLTSVD VDFPSPLDQY IIYIDGSAQP AACSTICTTQ LAELTSLVLA

HYQAMLLDER VHFPDPLPDA TWYTDGSSFI AGAAVTTSAQ RAELIALTQA
HYQALLLDDR VQFPDPLPDA TWYTDGSSFL AGAAVTTSAQ RAELIALTQA
HYQSLLLNDR VTFKDPWPGA SWYTDGSSFL AGAAVVTSAQ KAELIALTQA
HYQSLLLTER VTFKDPLTGE TWFTDGSSYV AGAAVVTSAQ KAELMALTQA
HYQGLLLDPR IIFPDALPNS VWYTDGSSFV AGAAVVTSAQ KAELIALAEA
KWEADLLRPE LTFPDPIPDP TLFSDGSYTT -GAAVVASAQ TAELLALAAA
KWEADLLRPE LTFSDPLEKP ILFTDGSYSK -GCAVVTSAQ TAELAAITLA
KWEADLLRPE LTFPDPLPHP ILFTDGSYSK -GGAVVTSAQ TAELCALTSA
KIQNKCTQSH ITFPDPLPQA WLYTDGCCYR AAYAVVASAQ LAEIIGLTRA
RWHRILTQED ITIPEPIMPT KWWIDGSRYW TGWAALVSAQ VAELVALREA
---------- ---PSPV--- TVFTDASSST ---GVVASVQ QLEARAVAMA
IWP------- ---PPPL--- TVFTDASSQT ---AVASSVQ QLEARAVQLA
---HYPPNKL IQFPQP-LNN LVFTDGSSTG ---AYTNSAQ LVELQALIAV
---HYPPNKL IQFPQP-LNN LVFTDGSSTG ---AYTNSAQ LVELQALIAV
---HLPKDPL LTFPHP-LEK VIFTDGSANG --VTYINTAQ QAEIVAVITA
PWKTLITREV F--PQPEIPA CLFSDGATGR --GAYCESAQ KGELAGLLAG
LWNTFLKTAP LAPPVPVINT CLFSDGSTSQ --AAYIKSAQ RAELLGLLHG
PWRTLLHLAL LQEPLPVLTT CLFSDGSSQK --AAYVNSAQ KGELLALLSG
DWILELQMGN IN-PS--VVP TYYTDGGKKN --LGYIGTNQ QLELRAIEEA
MWE--MQKWY YS-PEP--TS TIYTDGGKQN --AAYVVTHQ VAERMAIQMA
AWESNLINPY LKAPEP-IPG TWYIDGGRKL -KAAYWGSNQ KAEIQALLLA
TWETWWTEWQ AT-PEP-IEG TFYVDGAANR GKAGYVTTNQ KTELQAIQLA

LLERAKIITD SYYCAQALEN GYETAKIAEL RM-NEVVHQK AHVKEGGNEE
LLEAARVVTD SYYCAQALEN GFESARIAEL RT-QDVEHQK AHTKEGGNDE
LLESARLVTD SYYCAQALEN GFETAKIAEL KL-QDVEHQR AHTHEGGNDE
LEKKCFMVTD SDYIYQAVNN DFNTSKIDEL ITE-TVLHQT SHTSQNGNKE
IRRPVLIVTD SNYVAKSYSN GFVNNKIAEC KNLKHVIHEP GHQAEAGNAL
LKQPVLIVTD SDYVAKSYSN GFVNNKIADC KKHKHVIHEP GHQDLQGNNA
CKQPVLIITD SFYVAESASN GFVNNKIADC LSLKTIKHEK GHQSHTGNAL
CKKPVLVITD SFYVAESASN GFVNNKIAEC LSIKTIQHEK GHQINTGNAL
CKKPVLIITD SFYVAESTSN GFVNNKIAEC LSLKTIQHER GHQIYTGNAL
CKKPVLIVTD SFYVAESASN GFLNNKIAEC LQLKIIMHEK GHQPMTGNNL
CKKPVLIVTD SFYVAESVSN GFFNNKIADC IQLKIIIHEK GHQTASGNNL
CKKPVLIVTD SDYVARSVSN GFVNNKISDS LLLKIIVHEP GHKSYTGNNL
CKKPVLIVTD SDYVARSASN GFLNNKISDS LLLKTIVHEP GHQSKTGNSL
CKKPVLIVTD SDYVARSVSN GFVNNKISES LLLHTIVHEP GHQSSTGNAL
IKKPVLIVSD SVYLVKSFSN GFLNNKIAAC YQNKFLLHVP GHQLLTGNAL
LKKNILVVTD SNYVAKAYSN GFVNNRVADL KRLRVVTHEP GHQLDSGNNL
VKKPILIVTN SMYLAKSFSN GFVNNKIANC KQNKHMVHEP GHQQGTGNLL
IEIPVLIGND SFYIARGISN GFLDNKIATL LDDKITVHVP GHSKYGGNTL
IEIPVLICTD SFYIARGISN GFLDNKLATL LDDKITVHVP GHSKYGGNTL
IEIPVLICTD SFYIARGISN GFLDNKLATL LDEKITVHVP GHSKYGGNTL
----VLICTD SFYIARGISN GFLDNKLATI LDEKITVHMP GHSKYGGNTL

```
IEIPVLICTD SFYIARGISN GFLDNKLATI LDEKITVHMP GHSKYGGNTL
IEIPVLICTD SFYIARGISN GFLDNKLATL LDEKITVHVP GHSKYGGNTL
----VLICTD SFYIARGISN GFLDNKLATI LDEKITVHVS GHSKYGENTL
----VLICTD SFYIARGISN GFLDNKLATI LDEKITVHVS GHSKYGENTL
----VLICTD SFYIARGISN GFLDNKLATI LDEKITVHVP GHSKYGGNTL
IEIPVLICTD SFYIARGISN GFLDNKLATI LDEKITVHVP GHSKYGGNTL
----VLICTD SFYIARGISN GFLDNKLATL LDEKITVHVP GHSKYGRNTL
IEIPVLICTD SFYIARGISN GFLDNKLATL LDEKITVHVP GHSKYGGNTL
IEIPVLICTE SFYIARGISN GFLDNKLATL LDDKIIVHMP GHSKYGRNTL
IEIPVLICTD FFYIARGISN GFLENKLATL LDEKITVHVP GHSKYGGNTL
IEIPVLICTD SFYIARGISN DFLDNK-ATL LDDKITVHVP GHSKYGGNTR
IEIPVLICTD SFYIARGISN DFLDNK-ATL LDDKITVHVP GHSKYGGNTR
IEISVLVCTD SFSIARGVSN GFLDNKLATL LNEKINVHVP GHSKYGGNTL
VKQPVLICTD SVYAQRGYIR NFHDSRLDQL KRDKRVIHVP GHT--GGNGL
LTHLTLIISD SHYCVRAYLN NYRDSKVATF KGQLHVVHTL GHQRVGGNTL
LKMKLTVYTD SRYAFATTRR GLLTSELLEA LFLPSIIHCP GHQKGDGNRL
LKMKLNVYTD SRYAFATARR GLLTSELLKA LFLPSIIHCP GHQKGNGNRM
LREIINIYTD SRYAFATAQR GLLTSALLEA IHAPAIIHCP GHQKGEGNRM
LRLSINIYTD SRYAFATAQR GLLTSALLEA LHLPAIIHCP GHQKAKGNQM
LERRVTVYTD SRYAFGTVER GFVTAELLMA VQMPAVVHIP GHQSAQGNRR
CHLTVNIYTD SRYAYGVVHR GFVTSALLKQ IMKPSVIKIE AHTKGVGNAA
CQYSVNIYTD SRYAFGVLHR GFVTSALLEA VQLPSLMKCP AHTKGVGNAA
CLHSVNIYTD SRYAFGVLHR GFVTSALLEA VMMPSVMKCP AHTKGMGNAA
LTLTVNIYTD SAYAHGAVRR NFTTTGLISA VALPAIMKCK GHQVLNGNDA
LRLPLTLYTD STYVLGICRR GMVNADIWQL IEHDGIVKVK AHTKCSLNND
LLLPTNVVTD SAFVAKMLPS TAA---ILED ALSQAVLHVR SHSEVPGNDV
SDAALNVVTD SLYVARLVPD LAS---WLYN SLSRFICHVQ AHQSLEGNDA
LSAPLNIYTD SAYLAHSIIK HIS---QCQQ LIYNYIGHVR AHSGLPGNQR
LSAPLNIYTD SAYLAHSIIK HIS---QCQQ LIYNYIGHVR AHSGLPGNQK
FEEPFNLYTD SKYVTGLFSP RTK---HLQR LIHKYIGHIR GHTGLPGNAY
LAAPLNIWVD SKYLYSLLLQ ------LYKS LLRHFVGHVR SHSSASLNNY
LSSCLNIFLD SKYLYHYLQG ------LLPR LLSRYLHHVR SHTNLPLNAL
IRASLNIFLD SKYLIKYLLG ------HLPT LLHNYLHHIR SHTNLPLNEY
CKQKMNIVTD SRYAYEFM-- ------IMEL VHNKGVHWVP GHKGIPQNEE
LEDQVNIVTD SYYCWKNI-- ------IIQN IREKYFAWVP GHKGIYGNQL
LKAEMNIITD SQYVINII-- ------VLEE LEKKFIDWVP GHKGIPGNEE
LQDAVNIVTD SQYALGII-- ------IIEQ LIQKYLSWVP AHKGIGGNEQ

VDVTEAVHEG HAGREPVREI RRVLKDCVEC GRYNAGPWGS ICMDVPLGEK
VDVVVTMHEG HAGRGPMKQI QQVLKRCTVC GKYNAGPWGS ICMDVPMGEK
VDVVRSVHEG HAGKEPVKQV QRVLRDCEVC GQYNAGPWGS VCMDVPMGEK
VDLLEKLHIK HPTLN--SEY YKIKVNCPNC RKAIDSENDE ISMDFPIRNC
ADLCQLAHDA HLGLKRMIDA SRIVLNCTVC AQTNSTPFQV WYMDYPLPYQ
ADILELAHKG HLGIKNLKDI SKYIRTCTNC IITNTDPFQK YYMDYPLPYY
```

ADLASQAHKS HSGLKNMKDV VKVIGQCQQC LVTNPSPFDK FFIDYPLPYL
ADIVLQAHNA HTGLKNMKDV VKQLGRCKQC LITNASPFDK FFIDYPLPYL
ADIVQKAHNA HTGLKNMKDV VRQLGRCQQC LVTNAFPFDK FFIDYPLPYL
ADIISTAHNA HTGLKNLKDV VKSIRQCKQC LVTNATPFDK FYIDYPLPYL
ADIILQAHNA HTGLKNLKDV VKVIRQCKQC LVTNAAPFDK FFIDYPLPYL
ADLVKIAHES HAGLRNMKDV ISHIRTCKPC LTTDGSPFDK FFIDYPLPFV
ADLVKTAHEA HTGLKNMKDI ITVLRQCKPC LQTDSTPFDK FFIDYPLPFS
ADLVKQAHEA HTGLRKMKDV SHCLRTCMPC LQTNSTPFDK YYIDYPLPYS
ADLITKAHNS HMGAKNMKDV KHVLTICSQC QQVNSFPFDK IYMDYPLPYL
ADLIKEAHNS HAGLKKMKDI SSFLSTCNVC KMVNPLPFDK FYMDYPLPYV
ADLAQQAHNI HGGIKNMKTV RSVVANCEKC QVTNASPFEK FYMDYPLPHK
VDLLQEAHV- YVSLQSYRNV QRHLTQCEPF RTNKPLPFDK VYMDYPLPHN
VDLL--CHK- -----TFKQV QRRLAQCEPC RTNKPLPFDK VYMDYPLPHN
VDLLKEAHSG HTSLKSYRQV QRHLAQCEPC RTNKPLPFDK MYMDYPLPHN
VDLLKEAHSG HTSLKSYRQV QRHLAQCEPC RTNKPLPFDK MYMDYPLPHN
VDLLKEAHSG HTSLKSYRQV QRHLAQCEPC RTNKPLPFDK MYMDYPLPHN
VDLLKEAHSD HTSLKSYKQV QRHLAQCEPC RTNKPLPFDK MYMDYPLPHN
VDLL--THSG HTSLKSYRQV QRHLAQCEPC RTNKSLPFDK MYMDYPLPHN
VDLL--THSG HTSLKSYRQV QRHLAQCEPC RTNKSLPFDK MYMDYPLPHN
VDLLKEAHIG HTSLKSYKRV QRHLAQCEPC RTNKPLPFDK MYMDYPLPHN
VDLLKEAHIG HTSLKSYKRV QRHLAQCEPC RTNKPLPFDK MYMDYPLPHN
VDLLKEAHSG HTSLKSYRQV QRHLAQCEHC RTNRPLPFGK MYLDYPLPHN
VDLLKEAHSG HTSLKSYKQV QRHLAQCEPC RTNKPLPFDK MYMDYPLPHN
VDLLQEAHKS YVSLQSYKQV QWCLA-CEPC RTNKPLPFDK VYMDYPLPHN
VDLLR----- ---------- ---------- ---KPLPFDK VYIDYPLPHN
VDLLQEAHNH HVSLQSYRQV QRRLAQCEPC RTNKSLPFDK VYMDYPLPHS
VDLLQEAHNH HVSLQSYRQV QRRLAQCEPC RTNKSLPFDK VYMDYPLPHS
VDLLQEARNS HV-LLSYKQV QWYFAQCEPC RTKKPLPFDK VYMDYPLPHN
ADLIKEAHCI HLGKTGLTEC KRYVANCPEC LAVNVTLFDK IFLDFPLPYT
AALIKAAHEA HVGTLGLKEA RKYVLECDIC QQIKTSPFGC AYLDRPLSYK
ADLISHLHKT HLSTLNRLHL RQVTESCRAC AQINAGPGTH WEVDFEIKYK
ADLLDFLHQT HLSALNRRTL KDITETCQAC AQVNASPGTH WEIDFEVKYK
ADFVKGLHQT HLGRLNLSVA QKIINSCKAC AFTNATPGVY WEVDFEVKNK
ADYVQQIHRT HLGQLRLGVA DSVVKHCVPC QLVNANPGAH WEVDFEVKNK
ADFFSPTCIP PTGTIRQAQI QEIVDQCIGC QAMRPSPGRN WEVDFEVKYK
ADILLAIHGS HTHSYKAQTI DLILGHCQIC LKHNPKPFAH LQIDFQMCPM
ADIMRAIHGS HTHAYLAQAI DEVIAGCVVC LKHNPKPFSH LQVDFHMSPM
ADIMRAIHGS HTHAYLAKSI DEIMAKCAVC LKHNPKPFSH LQIDFHMSPM
ADLIREAHGT HEGQRYMEMT DLFCDNCTIC GNYNPKCFQD ISIDYDMGKR
VDLAQQYHHG HPSKVDMQHC KEITNTCLTC AKYKVLPCQK LQVDHPLGYR
ADEAKDLHTL HIGKA--QQA REVVQTCPHC NSAPALPLQI WQTDFLEPRS
ADEAVQKHAL HLGQE--TEV ERLVKSCSHC QKTPLWPNQI WQTDFMYPNH
ADSAQNAHTH HLNLM--EQA RQIVKQCPIC VTYLPVPNMI WQMDVHYSLK
ADSAQNAHTH HLNLM--EQA RQIVRQCPIC ATYLPVPNMI WQMDVHYSLK

ADSAQESHAH HQNFQ--EQA REIVKLCPNC PDWGHAPRVL WQMDVHVSLK
VDTPEQWHKT HCNRW--WDP RSPATLCETC QRLNPTPNHI WQADIHYKTY
TDSPADLHST HCGLQ--TEA SNILRSCHAC RKNNPQPNHI WQGDIHFKLY
TDTPQDLHKT HCNSS--QQA KSLLQTCYTC NIINSQPNHI WQGDVHYKRY
IDLAEEEHNW HQDLE--TAA EDIVQQCDVC QENKMP-IDH WQVDYHYE-K
ADEAQDEHEW HTSRN--TVA KQITQECPHC TKQGSG-PNH WQADCHLD-K
VDEAEINHEF HSDTE--MVA EEIRRKCPVC RIRGEQ-PGI WQMDCHFD-K
VDKAQEDHEY HTNSD--VVA KEIVASCDKC QLKGEA-PGM WQLDCHLE-K

YLLVLVDSMS GYVAVKPVRQ ANGSSVVSMC CYMGIPKELR
YLIVLVDSMS GFVVVRPVRK ANGNSVVSMC ACLGIPRELR
YLLVLVDSMS GFVTTKAARK ANGNSVVGMC SALGIPKELR
YFLLITDNAS EYTQIFPCKQ ATEEVVIESF KFRGIPNRVR
HALVIVDAGT GFTWIYPTKA QTANATVKAL TGTAVPKVLH
HVLVIVDEGT GYTWLYPTKA QTANATVKAL TGTAIPKVLH
HVLVVVDAMT GFVWLYPTKA PSANATVKAL TSIAVPKVIH
YVLVIVDGMT GFTWLYPTKA PSTSATVKSL TSIAIPKVIH
HVLVVVDSMT GFTWLYPTKA PTTNATVKAL TSIAVPKVIH
HVLVVVDSMT GFVWLYPTKA PSTSATVKAL TSIAIPKVLH
HVLVVVDSMT GFVWLYPTKA PSTSATVKAL TSIAVPKVIH
YVLVVVDAAT GFTWLYPTKA PSTNATITSL LGTAVPRVLH
YVLVIVDART GFTWLYPTKA PSTNATINSL LGTAIPKVLH
YVLVVVDSAT GFCWLYPTKA PSTRATVKSL LGIAVPKILH
YVLVLVDSCT GFTWLYPTKA PSANATVKAL TGTAVPKVLH
HVLVVVDAAT GFTWLYPTKA QTSKATIKVL TGLAIPKVLH
HILVVDDARM GYCWLFPTKA QNANATVKAL SGTAIPKVLH
HCLVIVDACT GFIWIYPT-Y QSASTTLASF ISLGLLRILH
HCLVIVDART SFVWIYPT-D QSASTTLTSF IWLGLPRILH
HCLVIVDACT GLVWVYPTRD QSASTTLTSF VSLELL-ILY
HCLVIVDACT GFVWIYPTRD QSASTTLTSF VSLELL-ILH
HCLVIVDACT GFVWIYPTRD QSASTTLTSF VSLELL-ILH
HCLVIVDACT GLVWVYPT-D QSASTTLTSF VSLELL-ILY
NCLVIVDACT GFVWVYPTRD QSASTTLTSF VSLELL-ILH
NCLVIVDACT GFVWVYPTRD QSASTTLTSF VSLELL-ILH
HCLVIVDACT GFVWIYPTRD QSASTTLTSF V-LELLCVL-
HCLVIVDACT GFVWIYPTRD QSASTTLTSF V-LELLCVL-
HCSVIVDACT GFVWIYLTQD QSASTTLTSF VLLGLPCVLH
HCLVIVDACT GFVWIYPTRD QSASTTLTSF IPLGLPRILH
HCLVIIDACT GFVWIYPTRD QSASTTLTSF ISLGLPCILH
HCLVIVDACT VFVCIYPTQD QSASTTLTSF IPLGLPLILH
HFFVIVDACT SFVWIYPT-D QSASATLTSF ISFGLPRILH
HFFVIVDACT SFVWIYPT-D QSASATLTSF ISFGLPRILH
HCLIIVDACT GFVWIYLP-D QSASTTASTY PTLWQGRCLH
AILILVESLS SFTWLLPCRD QSASTTVTAF PASIEPKCFH

YILVAVDSCS RFLWVWPQRS ADARTVIKDV LGTYACRTFH
YLLVFIDTFS GWAEAYPAKH ETAKVVAKKF PRYGIPQVLG
YLLVFIDTFS GWVEAFPTKK ETAKVVTKKF PRFGMPQVLG
YLLVFVDTFS GWVEAFPTKT ETAQIVAKKL PRYGVPKVIG
YLLVFVDTFS GWVEAYPTKK ETSTVVAKKF PRFGIPKVIG
YLLVMVDTFS GWVEAF-TKR ETAQVVAKAI PRYGVPEVLG
YALVIIDVFS KWPEIIPCNK EDAKTVCDII PRWGLPDQID
YALVIVDVLS KWPEVFSCNK EDATTVCDIL PRWGLPDQID
YALVIVDVFS KWPEVFSCNN EEAKTVCNII PRWGLPEQID
YLLVMVDRFS RWVEAIPTAK EDAKSVIKWI PRYGVPRQIR
YLTTMVDVYT GWFWAKPCRG PTTGATIAAI SIWGVPYSIQ
WLAVTVDTAS SAIVVTQHGR VTSVAAQHHI AVLGRPKAIK
WLAVTVDTYS GVIVATAHRG TKSRHAIHHI AYLGRPSVIK
YIHVSIDTFS GFLLATLQTG ETTKHVITHF SIIGLPKQIK
YIHVSIDTFS GFLLATLQTG ETTKHVITHF SIIGLPKQIK
YVHVTVDTYS HFTFATARTG EATKDVLQHF AYMGIPQKIK
ALHVFVDTYS GATHASAKRG LTTQMTIEGI VHLGRPKKLN
RLHVWVDTFS GAISATQKRK ETSSEAISSI AYLGKPSYIN
CLHVWVDTFS NAVSITCKTK ETSSETVSAI TILGKPLSIN
IILVWVETNS GLIYAERVKG ETGQEFRVQW YAMFAPKSLQ
IILTFVESNS GYIHATLLSK ENALCTSLAW ARLFSPKSLH
IILVGIHVES GYIWAQIISQ ETADCTVKAL LSAHNVTELQ
IILVAAHVAS GYIEAEVIPA ETGQETAYYL AGRWPVKVIH

**Data set S2. The alignments used to build the phylogenetic trees of endogenous/exogenous foamy virus in Fig. 3.**

Gag-Pol-Env
15 2093
BFV          ALQHNDIIIC RATSGPWGIG IRIHLQDPAG QPLPVVSAPM ATLENILNNF
EFV          AQADNDIINI RFTSGQWGIG VRLRLVDNTG QPLAVVAVSH NAARNIFNNV
FFV          ARGHGDIIAV RFTGGPWGPG VTIRLQDNTG QPLQLIAGPY NLIRTAFLDL
SFVmac       AARHREVIAL RMTGGWWGPA VSLLLQDDQG QPLPTLEAPW QDLRLAFDNI
SFVagm       ALNHREVIGL RMLGGWWGPG VSIFLQDDSG QPLQTIEAPW GELRQAFEDL
EFVcpz       ASLHGEIIGL RLTEGWWGQL VRLILQDEDN EPLQVLSGPL AELQLAFQDL
SFVgor       ASMHQEIIGL RMTEGWWGQM TRLILQDDDG EPLQVISGTL AELRLAFLEL
SFVora       AARHLETIGL RMLGGWWGEQ ARIILQDDDG EPLQVISAPW DQLRRAFHDL
SFVmar       AQRNGQTYAI AMTEGWWGDH IRVVFQDTSG NPLSVVGCTF NQASRVFDQI
SFVsqu       ARPHLTHYVL RMTEGWWGPF VRIILQDESG NPLSILETTL DEMDRVFDGI
PSFVgal      SQQHLDTLLV RITGGNWGPG IEVLLRD-TL GPLQILHLNY QDAIIIFDMI
SloEFV       AQGHQDILTV RVDAGPWGVG IQIDLQDGNR QPLPLLNVSY AQLMAKLGDV
SpuEFV       HYRDGDDYCL EIRNGEWGIG ISMRLRDVVF NPLTVLPINR QDLNLMAGIV
CoeEFV       ALADEQFLIC MMDYNDFGI- VGTLVLDNAG EIINILTFDY QELVNEGLQP
SnRV         ASPEAELEKV LKGLEEWGYK VQGGLAD-SK KALEKQKNRI AELEKEVADL

HIPHGVSRYG PLEGGDYQPG EQYSQGFCPV TQAEIEEEIT ILRERLMHLP
QPAGGPNRHG PLHDGQFQVG DDPSEHFVPI EENLIRREVR LLRERLLHLP
EPARGPERHG PFGDGRLQPG DGLSEGFQPI TDEEIRNEIR LLRERLQVIP
DVGEGTLRFG PLANGNYIPG DEFSLEFLPP AMQEIDAQIR GLEGQLRVIP
DVAEGTLRFG PLANGNWIPG DEYSMEFQPP LAQEIDAQIR GLERRIQAIP
DLPEGPLRFG PLANGHYVEG DPYSRSYRPV TMAETEVETR ALRRQLAVIP
DLPEGALRYG PLANGHYIQG DPYSLSYRPV TMAETEVEVR ALRRQLAVIP
DVGNGALRFG PLANGNYIPE DPYSTSYRPV NPQEMTIEIR GLRQELQVIP
DIGEGPSRFG PLADGMFLIT DNAWMDFVPL SAMEIQRENR DLNTNLRVLP
NLSPGTERYG PLCDGNFLYT DDAWNDFNPM SALEVKQENE DAHERLRVLP
IPSEGVHRHG PMFDGLWIHG DDYSMNFQPI TAHELTEEVE LLTERMAVIP
PFLTGVYRHG PLYTGHWLPG DDLSHHFLPI TMAELQEEVA LLRERFQALP
EIPRNIRRHG PISNADYISG SGY--NYTPA TTEELEQEYT LLRRNLRVLN
QFGAGLRRHG NLRGVLYHQG GPGW-QFAPL EPAEYRNQYQ ALRRRLQQLN
SAGRGADQVI AMDKELKKTA EKYQAKLEEL EEQLAESQVE GLKEEAEELP

ITHIRAVIGE TPANIREVPL WLARAVPALQ GVYPVQDAVM RSRTVNALTV
ITHIRAVIGE TPAQIRDVPL WLAQSIPALT GVYPAMDAGT LTRLVNAITA
INHLRSVIGN TPPNPRDVAL WLGRSTAAIE GVFPIVDQVT RMRVVNALVA
IQHIRAVTGN APSNPREIPM WIGRNASAIE GVFPIPTSDI RSRVINALLG
IQHIRAVTGN TPTNPRDIPM WLGRHSAAIE GVFPMTTPDL RCRVVNALIG
IQHIRSVTGE PPRNPREIPI WLGRNAPAID GVFPTTTPDL RCRIINALLG
IQHIRAVTGE PPRNPREIPI WLGRNAPAIE GVFPVNSPEV RCRVINAIIG

IQHIRAVTGE VPNNPRDIPM WIGRNAPAIE GVYPVTTPDL RARIINALIG
ISQIKAVIGE TPTDHKAVPL WVAKHAAAIE GVFPTGSPEV RCRVLNSLLT
ISQIRTVIGN TPVDPKKVPL WIAKSASAIE GVMPTNTPDI RCRLVNALLP
INVIRSVCGD TPSNPQDIPL WMGRIIPAIE GVFPIDNPNL RMRVVNALLA
MNHLRAVVGA TPNDPQAIAL WLGRNVQAIE GVMPINNGPM RKQVVNALLA
MANIRAVTGP TPKDFKEIPM WFESHLSALE AVTSTASPLQ KDEVMQ--FS
IGAIKNILGD PPNRGEDFLD WFQARHTQIE TITEGYTAEQ RRQLYQYILP
-AAVTAVGGA DPTRETGSAR WVCQVVTAMS PSWADVPPEI RNRVA-----

RHPGLALEPL ECGSWQECLA ALWQRTFGAT ALHALGDTLG QIANSDGIVM
RHPGLALGMN EAGSWHEAVH LIWQRTFGAT ALHALSDVLK GIAQRNGVVM
SHPGLTLTEN EAGSWNAAIS ALWRKAHGAA AQHELAGVLS DINKKEGIQT
RQLGLNLDPQ HCITWASAIA TLYVRTHGSY PLHQLAEVLR RVSNSEGAAA
GSLGLSLEPI HCVNWAAVVA ALYVRTHGSY PIHELANVLR AVVTQEGVAT
GNLGLSLTPG DCITWDSAVA TLFIRTYGQY PLHQLGNVLK GIADQEGVAT
GNLGLALTPT ECATWDSAVA TLFIRTHGTY PMHQLGNVIK GIVDQEGVAT
GKSGIHLTAP EAVTWASAVA AIFTRTHGSF PMHNLSAILT GIANGEGVES
GHGGMMLHPV DCVSWTNAAS VLFQRVHGVV PLHQLPKTLE EVAKTEGLLV
QHGGLILQPH ECNSWTQIAS ALYTRVNGMI PLHALPQTLS QVTKEEGILV
LHPGLAITEL NAQTWGQVLA VLHMRALGHT ALHQLPALLE TIVKTDGILP
SHATLHVTDQ EAQDWNSTIA AIYQRAHGTL ALHHLPTVLK DIANSDGVIV
SPSICFFDRA GVNNWESVLA NLYVKTHGQV GIADLNEILR KITQEQGIVR
TTWA-SVPIA HCNNVQATLD HLYIQITGQP PIIRLTKQIP LITASQGILV
-------TEK EIKAW----- LMKQGPGGGQ GLLEFTKLRQ GPTEN-----

AIELGLLFSD DNWDLVWGIC RRFLPGQAVC VAVQARLDPL PDNATRIVMI
ALEMGLMFTN DDWDLTWSVI RRCLPGQASV VTIQARLDAL PNNQARIIQA
AFNLGMQFTD GNWSLVWGII RTLLPGQALV TNAQSQFDLM GDDIQRAENF
AWQLGMMLTN QDYNLVWGMV RPLLPGQAVV TAMQHRLDQE VSDAARIVSF
GFQLGIMLSN QDYNLVWGIL RPLLPGQAVV TAMQQRLDQE VNDAARITSF
AYTLGMMLSG QNYQLVSGII RGYLPGQAVV TAMQQRLDQE IDDQTRAETF
GYTLGMMLSG QNFPLVYGII RGFLPGQAVV TAIQQRLDQE VDDQTRSDTF
AYNLGMMLSN GDFNLVYGIV RGLLPGQAAV AYMQQRLDAE PSDALRAQNF
AYNIGMTFTS NNFDLIWGII RPLVPGQAAV AMLQGYLDQY PRPQDKIEHF
AYQIGMTFTG QNFPLTWGIL RPLLPGQAVV AMMQGYLDQY PTDDLKAVNF
AYNMGMEVTQ QDFSYVWGIL RTLLPGQAFV LSMQNELDRL PA-AQRPGMF
AFTMGMMFSN DDYALVSGII RPLLPGQAAV VAVQAQLDIL PDDNAKASAF
AYGVGMKFLS -NHDLIWGIL KTLCKGDVLK AAIQSKLDLL TTEQEKIRSF
AYDHAYRGLR GDHVMAFELV REGIP-MAVR LQITAALQAL PTNQERHAQF
---------- ---------- ----PSNYLE KALELYLDSQ PGDRDGNKDD

SHIIRLDPLG RPMLPRRNNQ PPRGRQNAQT PRQEGNRLQN SQLPGNQPRY
GFIIRLDPLG RPLFPGGLTQ RDTGNQPQTQ PQQQNTFSNQ TNQRGSQRRY
PRVINLNIHG QSIRPRVQTQ PLQRQQQHS DVPEQRDQRG PSQPPSGGGY

VNHLNLNARG QNLRVSTGGQ TTARSSQGQQ PRQSESGDQN NQRQLNRGGY
NGHLNLNARG QSIRAQSAST SGNRTNQGNQ GQR-----DN NQRQSGQGGY
IQHLNLNARG QSIRASVTPQ PRPNSGRGRQ CPAPGQNDRG SNIQNSQGGY
IQHPNLNARG QSIRTGANST PRPGAGRGRQ RSDGNLQDNN PSSQNNQRGY
IQHLHLNHRG QSIRTSLPTS TRP-SGRGLS TPNRGSNNAN NNTQSSFGGY
PRFLRLNFLG QSIRSTQAPS RPNPGRGTAT PQAAGTSRSS SGSFGEQPRY
ASILRLNYMG QNIRNQSSPS LTVRGQPATT PTVTTSGSTS STTESSNPNY
PGLLQLNSRG QNIKTNTQQQ APKPRQQASA SPSSQGDNRS PQPQGSGPRY
PEIVTLNILG QPMHPPGSQQ CSNGGNRGKQ DQEGSQGQNH QQPQNQLRHY
PKIVQRDYLG NNPRKNLEQE DSKGKNQGSN RKQFQAQKDQ DSENPSAATY
ADIVRLNYNL AMILTPTQKQ GYSRGQVKWT PKQEKEESRP QSSQGKKHRY
PAFLQKNTSW QEMRDQFADK TGVGGQEGKG PRQNTTWKPK SGAIAAKQLA

PLRPNPQQPQ RNRNFSRGPA PVNEQSRGRG RSSQVEIKGN HLKGYWDSGA
FFRPRPSQPQ RRQLPQQ--Q QGSRRGPGRN TNSGVQIKGN SLKGFYDTGA
NFRRNPQQPQ RGPRPGGNPR GGGRGQGPRN GGGSVERKGV KIKGYWDSQA
NLRPRTYQPQ RSRTSGAGRE QGGRGNQNRN QRSAAEIKGT KLKAHWDSGA
DLRPRTYQPQ RSRTSGAGRG QQGRGNQNRN QRRAAEIKGT KLKAHWDSGA
NLRSRTYQPQ RSQTYKPAAG RGGRGNQNRN QRSSAEVKGT KLLAHWDSGA
NLRPRTYQPA RSRAPRSGAS RGGRGNQNRN QRPSAEVKGN KLIAHWDSGA
NLRPNTFRPQ RSRGPGGGGR AGGRRNQNRN SGQGAEVKGT KLKAHWDSGT
NLRPQVNRPS RSSTPRGQGE RSGRQRNNNA RQNQVEVKKT ELNGFWDTGA
NFRPRVNQPP RENTTQNNTR RPGSQGQNRN RNQNVEIHDQ KLIGYWDTGA
NFRPRVQPPD RNQQPGQGGQ RTQQTNQNRN QGNAVQLRAH HLKGFVDTGS
NLRPNVNHPR WEQQQQDQSK DRGVQSNSQP QRSGVQHRGK IF--------
DLRKGSQFPH RDFNSRGGIP TGNKMAENRQ AQGSVEIANK TFAALIDTGA
DLRPRHDGPW QNQGGQKWWK KGQDKQQPKY KKEGVCVGDT VVLALIDTGA
DIEKRLKDLT T--------- ---------- ----VEIEGH KIECLVDTGA

EITCVPAIYI IEEKKLITTI HNEKEHDVYY VEMKIEKRKV QCEVIATALD
EITCVPAIFL IEEERTIQTI HGITKEKVYY LTFKIQGRKL AAEVIGTQLD
DITCVPKDLL QGEQQNVTTI HGTQEGDVYY VNLKIDGRRI NTEVIGTTLD
TITCVPEAFL EDETMLIKTI HGEKQQDVYY LTFKVQGRKV EAEVLASPYD
TITCVPQAFL EEENIWIKTI HGEKEQPVYY LTFKIQGRKV EAEVISSPYD
TITCIPESFL EDEQTLIKTI HGEKQQNVYY LTFKVKGRKV EAEVIASPYE
SITCIPESFL EEEKTIIKTI HGQKEQKVYY LTFKVNGRKV EAEVIASPYD
TITCIPTVFL TDEDVLIKTI HGERRQPAYY LTFKINGRKV QAEVIASPYD
QITCIPEAFL KEEEAQIKTL HGTKLQSVYY LKFKVLGRKV EAEVTTSPFD
QITCIPQVYL EQEKHVIETV NGKTQRDAYY IKLKINGKKI ETEVIPSPFS
SITCFPKYTL VEEQYDISTI HGTVSQPVYY IKFKVNGKKV EAEVTESPLD
DIMTQWQRFL VSQYEEIQTI HGKQKMPMYY LTFKVDGHKF VGQVIPTELD
NQSCIQSKCL P----HVYST KGQCILESYF LEIQIAGKCL EIEAICDFTK
EHSVIDKCLV PKGHQKLQGL NGSVSFPVFN LTINISNTDI LLKVKANLQP
EVSLTSLQLQ AQRADHVDTT VGQKGKGCWR ISENILGNDL LRSLIVDQCN

YVLVAPVDIP WYKPGSTLSP QGQMRLKKLL DQYQALWQCW ENQVGHRRIE
YVIIAPSDIP WYKKYTNLSS EGKKYLKDLF IKYDNLWQKW ENQVGHRRIT
YAIITPGDVP WILKKSILSK KGKEELKQLF EKYSALWQSW ENQVGHRRIR
YILLNPSDVP WLMKKTALPK EQKELLQKLF LKYDALWQHW ENQVGHRRIK
YILVSPSDIP WLMKKTMLTG SYKEKLQSLF LKYDALWQHW ENQVGHRRIK
YILLSPTDVP WLTQQTALPE EQKQQLKALF TKYDNLWQHW ENQVGHRKIR
YILLSPMDVP WLVQKTALSE EFKKQLQTLF LKYDNLWQHW ENQVCHRKIR
YILLCPADVP WLQQQTALEG QFKQQLQNIL STFDTLWQHW ENQVGHRKIP
YVIISPSDIP WYKPQANINN EEKKQLAKLL DKYDVLWQQW ENQVGHRKIP
YALITPNDIP WFKPGADITK EEKGMLYKLL DKYDPLWQQW ENQVGNRQIT
YVILCPSDVP WLSTKTRVN- --QKQLQKLF IQYDDLWQKW ENQVGHRKIT
YALITAKVVP WIKLKSNISE QGKDKLRLLL HKYDSLWQKW EKQVSHRKIP
HDVVIAHEVN FIQNQADCAK NEKTKLRDIL YSLKPYFQQF DNQIGHRKIQ
YKLIISHQLP FFPQPTMLSQ ASKLKLKALL EKHYEVLQRH KNQTGFRDIE
GVLWQASEDN WMAAEYSIKS PGHYNLPELL ATKDELWNNV EAFATHRNLQ

PHKIATGALK PRPQKQYHIN PRAKADIQIV IDDLLRQGVL RQQNSEMNTP
PHKIATGTLN PKPQKQYRIN PKAKADIQIV IDDLLKQGVL KQQTSPMNTP
PHKIATGTVK PTPQKQYHIN PKAKPDIQIV INDLLKQGVL IQKESTMNTP
PHNIATGTLA PRPQKQYPIN PKAKPSIQIV IDDLLKQGVL IQQNSTMNTP
PHHIATGTVN PRPQKQYPIN PKAKASIQTV INDLLKQGVL IQQNSIMNTP
PHNIATGDYP PRPQKQYPIN PKAKPSIQIV IDDLLKQGVL TPQNSTMNTP
PHNIATGDYP PRPQKQYPIN PKARSSIQVV IDDLLKQGVL VQQNSTMNTP
PHNIATGTHP PRPQKQYPIN PKAKESIQIV INDLLKQGVL IQQNSIMNTP
PHNIATGTVA PRPQRQYHIN TKAKPSIQQV IDDLLKQGVL IKQTSVMNTP
PHIIATGTIN PKPQKQYHIN PKAKPSIQIV INDLLKQGVL KQQNSIMNTP
PHHIATETVA PKPQRQYHVN PAARPDIQIV IDDLLKQGVL KKQNSPMNTP
PHHIATGTVA PKPQKQYHIN YKAKLAIQTV INDLIKQGVL LHQNSSMNTP
PHDLSVKT-Q PKPQKQYPIN KAAINDIQKV INDLIAQGAL IKQYSTMNTP
PHQLK-GKIH P-AQKQYPIN TKAIPSIQVV INELLEQGVL VKQTSPTNMA
GMTASFTADH PKMIKQYPVP DASHASIKET VEALLEQGVL RKCNSTVNSA

VYPVPKADGR WRMVLDYREV NKVTPLVATQ NCHSASILNT LYRGPYKSTL
VYPVPKPDGR WRMVLDYRAV NKVTPAIATQ NCHSASLLNT LYRGQYKTTL
VYPVPKPNGR WRMVLDYRAV NKVTPLIAVQ NQHSYGILGS LFKGRYKTTI
VYPVPKPDGK WRMVLDYREV NKTIPLIAAQ NQHSAGILSS IYRGKYKTTL
VYPVPKPDGK WRMVLDYREV NKTIPLIAAQ NQHSAGILSS IFRGKYKTTL
VYPVPKPDGR WRMVLDYREV NKTIPLTAAQ NQHSAGILAT IVRQKYKTTL
VYPIPKPDGR WGMVLDYREV NKTIPLIAAQ NQHSAGILAT IVRKKYKTTL
VYPVPKPDGR WRMVLDYREV NKTIPLIAAQ NQHSAGILAS IYRGTYKTTL
IYPVPKPDGK WRMVLDYRAV NKTVPLIGAQ NQHSLGILTN LVRQKYKSTI
IYPVPKTEGK WRMVLDYRAV NKTIPLIAAQ NQHSAGILTN LVRQKYKSTI
VYPVPKPDGK WRMVLDYRAV NKTVPAIGAQ NCHAPGILSS LYRAKFKTTL

IYPVPKTNGS WRMVLNFRAV NKVIPLIAVQ NQYSIEILTQ MQREQYKTTL
VYPVPKPNGK WRMVLDYRAL NRVSPSFTVQ NLHMSGMLGN LERHKYKTTL
VLPVPKPDGT WRLVLDYRAL NKHSEPVRAQ NQHSSGILAN IERKAYKSSV
IWPVGKPDGS WRLTIDYRPL NSAVPTVAS- ---TPELFAK LEK-KYQSSL

DLANGFWAHP IKPEDYWITA FTWGGKTYCW TVLPQGFLNS PALFTADVVD
DLANGFWAHP IQESDQWITS FTWNGKSYVW TTLPQGFLNS PALFTADVVD
DLSNGFWAHP IVPEDYWITA FTWQGKQYCW TVLPQGFLNS PGLFTGDVVD
DLTNGFWAHP ITPESYWLTA FTWQGKQYCW TRLPQGFLNS PALFTADVVD
DLSNGFWAHS ITPESYWLTA FTWLGQQYCW TRLPQGFLNS PALFTADVVD
DLANGFWAHP ITPDSYWLTA FTWQGKQYCW TRLPQGFLNS PALFTADAVD
VLANGFWAHP ITPESYWLTA FIWQGKQYCW TRLPQGFLNS PALFTADVVD
DLANGFWAHP ITPNSYWLTA FTWQGKQHCW TRLPQGFLNS PALFTADVVD
DLSNGFWAHP ITKDSQWITA FTWEGKQHVW TRLPQGFLNS PALFTADVVD
DLSNGFWAHP IDQDSQWITA FTWEGKQYVW TRLPQGFLNS PALFTADVVD
DLSNGFWSHP ITPESYWLTA FTWQGSQYVW TRLPQGFLNS PALFTADVVD
DLSNGFWAHP IRKESYWLMA FTWEGKQLVW TRLPQGFINS PALFTANIVD
DLSNGFWAHP IREEDRYLTA FTWQGTQYCW TRLPQGYLNS PAMFSADVIQ
DLANGFWSHP IREEDWPKMA FTWCGFQYLW TRLPQGFLNS PALFSADVIS
DISNGFWSIR LEEECQYLFA FTFDTQQYTW TRLPQGFHAS PGIFHQALYN

ILKDIPNVQV YVDDVYVSSA TEQEHLDILE TIFNRLSTAG YIVSLKKSKL
LLKDIPNVEV YVDDVYFSND TEEEHLKTMD LLFQKLQTAG YIVSLKKSKL
LLQGIPNVEV YVDDVYISHD SEKEHLEYLD ILFNRLKEAG YIISLKKSNI
LLKEIPNVQA YVDDIYISHD DPQEHLEQLE KIFSILLNAG YVVSLKKSEI
LLKEVPNVQV YVDDIYISHD DPREHLEQLE KVFSLLLNAG YVVSLKKSEI
LLKEVPNVQV YVDDIYLSHD NPHEHIQQLE KVFQILLQAG YVVSLKKSEI
LLKEISNVQA YVDDIYLSHD DPQEHLDQLE KVFQILLQAG YVVSLKKSEV
LMKHIPNVQV YVDDLYLSHD DPQEHLQVLQ QVLHILHDAG YVVSLKKSAI
LLKNIPGISV YVDDIYFSTE TVSEHLKILE KVFKILLEAG YIVSLKKSAL
LLKEIPNVNV YVDDIYVSTE TINQHFQVLD KIFQKLLQAG YVVSLKKSNL
LCKHIPNVSA YVDDIYVSND TAEEHLRTLE QLFRTLMSAG YIVSLKKSKI
ILKEIPDVEV YVNDIYFSNV TEEQHLITLK QVLKILLKSG YIVSLKKSEI
LLKNIPGVNS YMDDIYFTNE NLDQHLATLK QVVTVLGEAG YIINLKKSQI
LVGQLPGVYC YVDDIYLTHD TEEEHLKILD QVLEILIKAG YVINIKKSKL
GLASCKKLLQ YVDDILLMSE DRDHHLRSLA ILLQGLKDLG VKINPKKSHF

AKETVEFLGF SISQNGRGLT DSYKQKLMDL QPPTTLRQLQ SILGLINFAR
GQHTVDFLGF QITQTGRGLT DSYKSKLLDI TPPNTLKQLQ SILGLLNFAR
ANSIVDFLGF QITNEGRGLT DTFKEKLENI TAPTTLKQLQ SILGLLNFAR
AQREVEFLGF NITKEGRGLT DTFKQKLLNI TPPKDLKQLQ SILGLLNFAR
AQHEVEFLGF NITKEGRGLT ETFKQKLLNI TPPRDLKQLQ SILGLLNFAR
GQRTVEFLGF NITKEGRGLT DTFKTKLLNV TPPKDLKQLQ SILGLLNFAR
AQKTVEFLGF NITKEGRGLT EAFKAKLLDI TPPKDLKQLQ SILGLLNFAR

AQKVVEFLGF NITKTGRGLT DAFKEKLLNI SPPQNLKQLQ SILGLMNFAR
LRYEVTFLGF SITQTGRGLT SEFKDKIQNI TSPRTLKELQ SILGLFNFAR
CRYEVTFLGF TISKYGRGLT EEFQEKLRNI SPPNSLKQLQ SILGLLNFAR
GVSAVDFLGF EITDDGRGLT SAFKEKLVNI QPPSSLKQLQ SILGFLNFTR
AKEEVTFLSF NITKEGHGLT AKFREKLLNI SAPKTLKQLQ SILGLLNFAH
CRSKVKFLGF LLTDSGRGLT QEFKEKLLTL QPPKTLKELQ SVLGFLNVAR
CRKVVDFLGF SLSDEGRGLS DQYREKLAAI KPPQTLRQLQ SVMGLLNFSR
CKDQVQYLGV NVGADTRSLI DARSQLIRTL DIPLTVQGLR SALGLFNFCR

NFLPNFAELV APLYQLIPKA CIPWTMDHTT QLKTIIQALN STENLEERRP
NFIPNYSELI TPLYQLIPLA YIPWETKHTA ILQKIIKELN ASENLEQRKP
NFIPDFTELI APLYALIPKS YVPWQIEHST TLETLITKLN GAEYLQGRKG
NFIPNYSELV KPLYTIVANA FISWTEDNSN QLQHIISVLN QADNLEERNP
NFIPNFSELV KPLYNIIATA YITWTTDNSQ QLQNIISMLN SAENLEERNP
NFIPNFAELV QTLYNLIASS YIEWTEDNTK QLNKVIEALN TASNLEERLP
NFILNFAELV KPLYSLISSA YIEWSNENTV QLQTIIKALN NADNLEERIP
NFIPNYAERV KPFYSLISTA NILWNDELTS QLQELITLLN QADNLEERKP
NFVPNFSEII KPLYSLISTA NIKWTSEHTR YLEEIVSALN HAGNLEQRDN
NFIPNFSELI KPLYELISTA SISWEPKHSQ ALNNLIIALN HADNLEQRNG
NFVPNYSELV KPLYNLVATA RISWTLENTE QLNQVISALN AADNLSERKT
NVITDFAELT KPLYLVISRA HIQWMEKEGM ALQEIIKKLN NASYLENRDI
ILVPDYAQRT KPLYNLIPLA F--WTLEAQQ ALDKLIVLIN QAAELNTRNN
LFVKGFSELA KPLYDLITVK PYHFTPQAHE ALTKLLSAIQ TAPNLTNRDN
AWIPEFSRKT QSLYDMLKGD KLKWTEDNLN KFKLLKDEVA SACVLGLPDP

DVDLIMKVHI SNTAGYIRFY NHGGQKPIAY NNALFTSTEL KFTPTEKIMA
DVELIVKVHV SPTAGYIKFA NKGSIKPIAY HNVVFSKTEL KFTITEKVMT
DKTLIMKVNA SYTTGYIRYY NEGEKKPISY VSIVFSKTEL KFTELEKLLT
ETRLIIKVNS SPSAGYIRYY NEGSKRPIMY VNYIFSKAEA KFTQTEKLLT
EVRLIMKVNT SPSAGYIRFY NEFAKRPIMY LNYVYTKAEV KFTNTEKLLT
DQRLVIKVNT SPSAGYVRYY NESGKKPIMY LNYVFSKAEL KFSMLEKLLT
EKRLIIKVNT SPSAGYVRYY NETGKKPIMY LNYVFSKAEL KFTLLEKLLT
TTRLIIKVNS SSHAGYIRYY NEGSKKPILY INYVFSKAEE KFSMLEKLLT
ESPLVVKLNA SPKTGYIRYY NKGGQKPIAY ASHVFTNTEL KFTPLEKLLV
EVPLVIKINA SNTTGYIRFY NKNGKRPIAY ASHVFNHTEQ KFTPVEKLLT
GVPLVVKSNC SPTAGYIRFY NQGDRKPIQY VNYIFSKTEL KFTPLEKQLT
QKPLIIKLNS SPTAGYIRMY NKGGKKPIQY VNFIFTPAEI KFKPTEKLLT
TASLEILVRA SHKGGFASYY NYGDSKPLQY ISYVFSNAEQ KFLPIERVLC
TRALVIRIVT SPRSAVAAYY NVGDKLPIQY VSYNFSQPEM KFSKIEKVCV
TLPFLVQKDD NGMWHVLGFY SR-------- ---------- KMTPVESNLG

TIHKGLLKAL DLSLGKEIHV YSAIASMTKL QKTPLSERKA LSIRWLKWQT
TIHKALLKAF DLAMGQPIWV YSPIHSMTRI QKTPLTERKA LSIRWLKWQT
TVHKGLLKAL DLSMGQNIHV YSPIVSMQNI QKTPQTAKKA LASRWLSWLS

TMHKGLIKAM DLAMGQEILV YSPIVSMTKI QRTPLPERKA LPVRWITWMT
TIHKGLIKAL DLGMGQEILV YSPIVSMTKI QKTPLPERKA LPIRWITWMS
TMHKALIKAM DLAMGQEILV YSPIVSMTKI QKTPLPERKA LPIRWITWMT
TMHKALIKAM DLAMGQEILV YSPVVSMTKI QKTPIPERKA LPIRWITWMT
TLHKALIKAV DLAMGTEIMV YSPIVSMTKI QKTPLPERKA LPVRWITWMT
TMHKALIKAI DLALGQPIEV YSPIISMQKL QKTPLPERKA LSTRWITWLS
TMHKAIIKGI DLAIGQPIEI YSPIVSMQKL QKITLPERKA LSTRWLSWLS
VLHKAILKGL DLAGGEDIHF YTPIASISKL QRTPIPERKA LHVRWLTWIT
TMHKAIIKGL DLSQGAQVHI YSPLASPTHI QKTPLPERKG LHSQWITWMT
MCNLAILKGK DLAQGQKMIV KTPIASLRQV KKGSIPNAKA LHSRWVQWMS
MVNMALIKGR VLAHEQDIEI HTAIPALQTY TRASEPNAKA LATRWDLWCT
ICEQYAISAC NLGFGRKIIV HSPVKFI--L QTTPNVSNQR LA-RW-H---

YFEDPRIKFH HDATLPDLQN LPQQDTGKEM TIPLLHYEAI FYTDGSAIRS
YFEDPRLIFH YDDTLPDLQN LPQTTLGNEV DIPLSEYEVV FYTDGSSIKS
YLEDPRIRFF YDPQMPALKD LPAVDTGKDN KKHPSNFQHI FYTDGSAITS
YLEDPRIQFH YDKSLPELQQ IPNVTEDVIA KTHPSEFAMV FYTDGSAIKH
YLEDPRIQFH YDKTLPELQQ VPTVTDDIIA KIHPSEFSMV FYTDGSAIKH
YLEDPRIQFH YDKTLPELKH IPDVYTSSIP PLHPSQYEGV FCTDGSAIKS
YLEDPRIQFH YDKTLPELKN IPDVLTENSS KIHPSQYNSV FYTDGSAIRS
YLEDPRITFH YDKTLPELKD VPSVYQNDIP IVHPSQYSMV FYTDGSAIKN
YLEDPRITFY YDKTLPDLKN VPETVTDKKP KMPIIEYAAV FYTDGSAIRS
YIEDPRFLFI YDKTLPDLKE MPPTQTDDYN PMPLHQYLAV FYTDGSSIKS
YLEDPRFHFY YDETLPPLAE LPEIAQGN-- QLPLKEYTSV FYTDGSAIKN
HFKNPQLIFH HDPTLPDIQN LPQPLSEDNM QIDLTSYTAV YYTDGSAIKN
HFENPQIEFE YVEPSNDLEN LPAFSIEPVS SKPLHEYQKV IYTDGSAMSC
LFSDQRLTFK TVTNMPDLDC LPPFEDEVAV ASPLEQYVEV YYSDGSAKRN
-----RILTQ EDITIETDAS IPEPYEGEQH QCPTGE---K WWIDGSALRE

PKPNKTHSAG MGIIQAKFEP DFRIVHLWSF PLGDHTAQYA EIAAFEFAIR
PKKDKQHSAG MGIIAVRYQP QMNIIQEWSI PLGDHTAQFA EIAAFEFALK
PTKEGHLNAG MGIVYFINKD NLQKQQEWSI SLGNHTAQFA EIAAFEFALK
PDVNKSHSAG MGIAQVQFIP EYKIVHQWSI PLGDHTAQLA EIAAVEFACK
PNVNKSHNAG MGIAQVQFKP EFTVINTWSI PLGDHTAQLA EVAAVEFACK
PDPTKSNNAG MGIVHAIYNP EYKILNQWSI PLGHHTAQMA EIAAVEFACK
PDPTKSHNAG MGIVQVKFSP ELQVINQWSI PLGNHTAQMA EIAAVEFACK
PNPTKTHSAG MGVVQGKFNP EFQVVNQWSI PLGNHTAQLA EVAAVEFACK
PDKNKSHSSG MGIVHAVFKP ELTIEHQWSI PLGDHTAQYA EISAVEFACK
PDPTKTHSSG MGIVQAIYEP NFQIKHQWSI PLGDHTAQYA EIAAVEFACK
PNPKKAHSAG MGTVEVTYNP EYKVLHEWSF PLGDHTAQYA EIAACEFAIK
PNPQKTHSAG IGIVKGKFDP NFSIIKQWRF PLGDHTAQYA EISALEFAVK
KQGKHMWKAG YAVVIGTFND EYHMSDSIQM PLGNNTAQYA ELMAV----H
QQKNY---AG IGIVKGKFTT HFEPEETKAV PLGPASAQYA EVMALLDAVK
DKKNQLGGAL EGHV------ ---------- -----SAQVA ELVAL----R

RATGIRGPVL IVTDSNYVAK SYNEELPYWE SNGFVNNKKK TLKHISKWKA
QAIRKMGPVL IVTDSDYVAK SYNQELDFWV SNGFVNNKKK PLKHVSKWKS
KCLPLGGNIL VVTDSNYVAK AYNEELDVWA SNGFVNNRKK PLKHISKWKS
KALKISGPVL IVTDSFYVAE SANKELPYWK SNGFLNNKKK PLRHVSKWKS
KALKIDGPVL IVTDSFYVAE SVNKELPYWQ SNGFFNNKKK PLKHVSKWKS
KALKVPGPVL VITDSFYVAE SANKELPYWK SNGFVNNKKE PLKHISKWKS
KALKITGPVL IITDSFYVAE STNKELPYWK SNGFVNNKKK PLKHVSKWKS
QALKITGPVL IITDSFYVAE SANKELPYWK SNGFVNNKKK PLKHVSKWKS
KANNISGPVL IVTDSDYVAR SVNEELPFWR SNGFVNNKKK PLKHISKWKN
KALQVTGPVL IVTDSDYVAR SVNNELNFWR SNGFVNNKKK PLKHISKWKS
KASLLRGPVL IVSDSVYLVK SFNEELPFWI SNGFLNNKKK PLQHISKWKT
KAMMDKGPIL IVTNSMYLAK SFNEELDIWI SNGFVNNKKK PLQHISKWKV
KAIEISPPVL ICTDSFYIAR GINEELPIWR SNGFLDNKRK PLKHAHRWQK
QATD-TGPVL ICTDSVYAQR GYTEDLHYWA IRNFHDSRNA KLKYADKWKQ
EALRLQRPLT LYTDSTYVLG ICTKYLAVWK RRGMVNSNQN ILQEI--WQL

IAECKNLKAD IHVIHEPGHQ PAEASPHAQG NALADKQAVS GSYKVFSNEK
IADCKKHKAD IHVIHEPGHQ NDLQSPYAMG NNAADKLAVK ASYTVFSVQL
VADLKRLRPD VVVTHEPGHQ KLDSSPHAYG NNLADQLATQ ASFKVHMTKN
IAECLQLKPD IIMHEKGHQ QPMTTLHTEG NNLADKLATQ GSYVVHCNTT
IADCIQLKPD IIIIHEKGHQ PTASTFHTEG NNLADKLATQ GSYVVNINTT
IAECLSIKPD ITIQHEKGHQ PINTSIHTEG NALADKLATQ GSYVVNCNTK
IAECLSLKPD ITIQHERGHQ PIYTSIHTEG NALADKLATQ GSYVVNNNDK
IADCLSLKTG ITIKHEKGHQ PSHTSVHTEG NALADKLATQ GSYVVNNIIK
ISDSLLLKRD IIIVHEPGHK PSYTSIHTQG NNLADKLATQ GSYTVNNIVN
ISESLLLHKN ITIVHEPGHQ PSSTSVHTQG NALADKLAVQ GSYTINNITK
IAACYQNKKD IFLLHVPGHQ KLLTDEHAQG NALADKLAVQ SSHKVLFIKK
IANCKQNKPS IHMVHEPGHQ KQGTSIHTKG NLLADQLAVQ SSHMVGMVTL
LATLLDDKPL ITVMHVPGHS K--YGSHVNG NTLVDLLAKE AMKVLTRSQK
LDQLKRDKPL VRVIHVPGHT P--GTVHSCG NGLADSLAQG AVKILTRAQR
IEH--DSTQT LGIVKVKAHT QRKCSTHEQL NNDVDQPAKQ YAKVIAPLQL

PSLDAELEQV LSTPNPQGYP NKEYKLVNGL CYVDREEGLK IIADRVKLCQ
PSLDAELHQL LDKPNPKGYP SKEYTLRDGQ VYVKRTDGEK IIDDRVKILE
PKLDIEQIKA IQARLPVGYP KQTYELQNNK CMVLRKDGWR EIRERYKLIK
PSLDAELDQL LQGHYPPGYP KQKYTLEENK LIVERPNGIR IVADREKIIS
PSLDAELDQL LQGQYPKGFP KHQYQLENGQ VMVTRPNGKR IISDRPQIIL
PNLDAELDQL LQGNNVKGYP KQTYYLEDGK VKVSRPEGVK IISDRQKIVL
PNLDAELDHL IQGKYPKGYP KQTYYMEDGK VKVNRPEGTK IILERAGIVQ
PSLDAELDQV LQGNLPKGYP KHVYTLEEGK VIVKRPEGTK IIADRKLLAS
PSLDAELEQL INGHSVKGYP SRKYILKEGQ VFVLRPEGEK IISDRLALVK
PSLDTELRAV LEGKLPKGYP KNKYEYNSPN LIVIRKEGQR IISDRPKLVK
PSLDAELIQV MEGKYPKGYP HKVYAQDNGK IIVTLPNGQR EIGDRLALIT

PSLDKELEQV LDSPNPKGYP VKIYLLENGN VIIEQDEGKR IIMERVKLAQ
KCLDGELTQC ISPINPKGYP SADYALKDGK CVVTFTNGKH VIDTRPNLIQ
THMDSILQQC LDPPNPPGYP TAQYTNEHGQ CVVTMPGGTY VIHARPGLIK
PDFQKQLAEI MPQPSPEAYH DKTYIGKEGA AILGQAEGVH VSTEGYALAQ

LAHDSAHLGR SALLLKLQQK YWWPRMHIDA SRIVLNCTVC AQTNSTNQKP
LAHKSGHLGK NTMYIKILNK YWWPNLIKDI SKYIRTCTNC IITNTDNVPN
EAHNISHAGR EAVLLKIQEN YWWPKMKKDI SSFLSTCNVC KMVNPLNLKP
TAHNIAHTGR DATFLKVSSK YWWPNLRKDV VKSIRQCKQC LVTNATNLTS
QAHNIAHTGR DSTFLKVSSK YWWPNLRKDV VKVIRQCKQC LVTNAATLAA
QAHNLAHTGR EATLLKIANL YWWPNMRKDV VKQLGRCKQC LITNASNKTS
KAHNLAHTGR EATLLKIANL YWWPNMRKDV VRQLGRCQQC LVTNAFNQTS
QAHKLSHSGR EATLLKLSNT YWWPNMRKDV VKVIGQCQQC LVTNPSNLTS
IAHEFSHAGR EATVLRLQDK YWWPNMRKDV ISHIRTCKPC LTTDGSNLTP
QAHELAHTGR EATLLRLQNQ YWWPKMRKDV SHCLRTCMPC LQTNSTNLTT
KAHNISHMGR EAVLAKIQNV YWWPNMKKDV KHVLTICSQC QQVNSFNLKP
QAHNTIHGGW EATLIKLKNK YWWPNMIKTV RSVVANCEKC QVTNASSQIP
EAHNHVHHSE S---LK--RS YWWPGLRKQV QRRLAQCEPC LRTNPGPVTR
EAHCHIHLGG NNTGKTLRRL YWWPGLYTEC KRYVANCPEC LAVNVTPTTR
QYHHYGHPSE ESLRKVLTKR FVWEDMGQHC KEITNTCLTC AKYKV--LRA

RPPLVIPHDT KPFQVWYMDY IGPLPPSNGY QHALVIVDAG TGFTWIYPTK
KSYIVQEKTG LPFQKYYMDY IGPLPPSDGY YHVLVIVDEG TGYTWLYPTK
ISPQAIVHPT KPFDKFYMDY IGPLPPSEGY VHVLVVVDAA TGFTWLYPTK
PPILRPVKPL KPFDKFYIDY IGPLPPSNGY LHVLVVVDSM TGFVWLYPTK
PPILRPERPV KPFDKFFIDY IGPLPPSNGY LHVLVVVDSM TGFVWLYPTK
GPILRPDRPQ KPFDKFFIDY IGPLPPSQGY LYVLVIVDGM TGFTWLYPTK
GPILRPTRPL KPFDKFFIDY IGPLPPSNGY LHVLVVVDSM TGFTWLYPTK
GPILRPERPT KPFDKFFIDY IGPLPPSNGY LHVLVVVDAM TGFVWLYPTK
IPPKPQLRPE KPFDKFFIDY IGPLPPSHGF VYVLVVVDAA TGFTWLYPTK
TRPFQQIRPS KPFDKYYIDY IGPLPPSEGY SYVLVVVDSA TGFCWLYPTK
QPPQTIARHV HPFDKIYMDY IGPLPPSDGY LYVLVLVDSC TGFTWLYPTK
TPPKTIIHPD KPFEKFYMDY IGPLPSSHGH KHILVVDDAR MGYCWLFPTK
PPYLKNPKPL SPFDKVYMDY IGPLPPSHGH NHCLVIVDAC TSFVWIYPTR
PPNLRPPRGA L-FDKIFLDF VGPLPRSNGY TAILILVESL SSFTWLLPCR
GPPMGVGRSA EPCQKLQVDH VGPLPGTHGY RYLTTMVDVY TGWFWAKPCR

AQTANATVKA LTHLTGTAVP KVLHSDQGPA FTSSILADWA KDRGIQLEHS
AQTANATVKA LNHLTGTAIP KVLHSDQGSA FTSATLVAWA KDKGIQMEYS
AQTSKATIKV LNHLTGLAIP KVLHSDQGSA FTSEEFAQWA KERNIQLEFS
APSTSATVKA LNMLTSIAIP KVLHSDQGAA FTSSTFADWA KEKGIQLEFS
APSTSATVKA LNMLTSIAVP KVIHSDQGAA FTSATFADWA KNKGIQLEFS
APSTSATVKS LNVLTSIAIP KVIHSDQGAA FTSSTFAEWA KERGIHLEFS
APTTNATVKA LNVLTSIAVP KVIHSDQGAA FTSSTFADWA KERGIQLEFS

APSANATVKA LNMLTSIAVP KVIHSDQGAA FTSSTFADWA KEKGIHLEYS
APSTNATITS LNILLGTAVP RVLHSDQGSA FTSSTFADWA KEKGIQLEFS
APSTRATVKS LNFLLGIAVP KILHSDQGSA FTSSDFANWA KEKEITLEFS
APSANATVKA LTHLTGTAVP KVLHSDQGSA FTSSTLVDWA KERGIRLEYS
AQNANATVKA LNFLSGTAIP KVLHSDQGSA FTSATLQQWT KDRGIQLEFS
DQSASTTVRT LTSFISLGLP RILHSDKGGA FTSHQMQSFA KSFGIVLEYS
DQSASTTVTA LSFFPASIEP KCFHSDQGGG FTSQLFKKMC SERNIRVEYS
GPTTGATIAA LEHISIWGVP YSIQSDNGTA FTSKAMQEWA NTYGIEWKVG

APYHPQSSGK VERKNSEIKR LLTKLLAGRP TKWYPLIPIV QLALNNTPNT
SPYHPQSSGK VERKNSEIKR LLTKLLVGRP TKWYPLIPTV QLALNNTPNA
TPYHPQSSGK VERKNSEIKK LLTKLLVGRP LKWYNLISSV QLALNNTHVV
TPYHPQSSGK VERKNSDIKR LLTKLLIGRP AKWYDLLPVV QLALNNSYSP
TPYHPQSSGK VERKNSDIKR LLTKLLVGRP AKWYDLLPVV QLALNNSYSP
TPYHPQSSGK VERKNSDIKR LLTKLLVGRP TKWYDLLPVV QLALNNTYSP
TPYHPQSSGK VERKNSDIKR LLTKLLVGRP TKWYDLLPVV QLALNNSYSP
TPYHPQSSGK VERKNSDIKR LLTKLLVGRP TKWYDLLSTV QLALNNAYSP
TPYHPQSSGM VERKNREIKR LITKLLVGRP TKWYPLLPTI QLALNNTYSV
TPYHPQSSGK VERKNQEIKK LLTKLLVGRP AKWYPLIPSV QLALNNTYSP
TPYHPQSSGK VERKNSEIKR LLTKLLVGRP LRWYPLIPTV QLALNNTPNV
TPYHPQSSGK VERKNGKIKR VLTKLLYGWP QKWYPLIPFV QLSINNIPSS
TPYHPQSAGV VERKNGEIKR ALTKLLVGRS RQWYSLLPLV QLGLNNLPRS
TPHHPQSAGV VERKNRGLKA ALTKLVRNRP RKWFQVLDIV QTGLNNTPIA
AIYHPQSQGK VERKHRLLKD RL-KRATHEG KNWVQALPSI LLFINSMH-P

RQKYTPHQLM YGADCNLPFE NLDTLDLTRE EQLAQTTARS WTPSPGLLVQ
KIGKTPHQLM YGVDCNLPFQ DLSTLDLTRE EQLA-----K WTPCPGLLVQ
STKYTPHQLM FGIDCNLPFA NKDTLDWTRE EELAPTCS-G WSPYVGQLVQ
SSKYTPHQLL FGVDSNTPFA NSDTLDLSRE EELSPASSRS WSPSVGQLVQ
SSKYTPHQLL FGIDSNTPFA NSDTLDLSRE EELSPASIRA WSPSVGQLVQ
VLKYTPHQLL FGIDSNTPFA NQDTLDLTRE EELSPASSRS WSPVVGQLVQ
SLKHTPHQLL FGIDSNTPFA NQDTLDLTRE EELSPASTRS WSPIGGQIVQ
ILKHTPHQLL FGVDANIPFA NQDTLDLTRE EELSPASSRS WTPAVGLLVQ
HYKKTPHQLL FGVDGNVPFA NQDTLDLTRE EELSPASKRA WLPSVGLLVQ
KIKLTPHQLL FGVDGNIPFA NSDTLDLKRE EELAPSTAKT WTPSVGLLVQ
SSGKTPHQLL FGVDCNLPFA NKDTLDLTRE KQLSPPPSRS WRPSVRLLVQ
QTHQTPHKLM FGVDSNLPFA NVDDANLSRE EQLSSSCTSG WKPFIGQFIQ
DCHLTPYKLL FAKDMTTPL- ---------- EQLASSSSYL STGAVG--TD
RNEHTPFFLM YGQHMNTPLT NAEPSHPDRM EALAPESNRL WIPKVGEWVQ
RDQFSAYELM TGRVPHLGGY HPHPLETAKE EEVRAEAHSH WTQEPGCIVL

ERVARPAQLR PKWRKPTPIK KVLNERTVII -DHLGQDKVV SIDNLKPAAM
ERVNRPAQLR PKWKKPTPIL KVLNPKTVVI AGPGGQERIV SIDNLKKTPM
ERVYRPSQLR PKWRKPTKVL EILNPRTVII VDHLGQRKSV SIDNLKPTAQ

```
ERVARPASLR PRWHKPTAIL EVVNPRTVII LDHLGNRRTV SVDNLKLTAM
ERVARPASLR PRWHKPTPVL EVINPRAVVI LDHLGNRRTV SVDNLKLTAM
ERVARPASLR PRWHKPSTVL EVLNPRTVVI LDHLGNNRTV SIDNLKPTSM
ERVPRPASLR PRWHKPSRIV DILNERTVVI VDHLGNNRTV SIDNLKLTPM
ERVARPASLR PRWHKPVKIL EVLNPRTVVI LDHLGNNRTV SVDNLKLTAM
ERVARPSQLR PKWKKPTPIL EVVNDRTVVI LDNQGQRRTV SIDNLKLTPM
ERVYRPSQLR PKWKKPTPIL EVLNERTVVI -DNNGQRRTV SVDNLKYTPM
ERVYRPSALR PKWRKPTPIL EVHSDRLVTI KDHLGNIKKV STDNLKLTPM
ERVQKYTPLC PRWKKPTKIL TVFDDHTVEI LDPLGQRRKV SIDNLKPTAY
DRIKSRTGFS PAQHRYQPCS SILQEKAVSR -DPSSRPRST NPQSFDITKL
EKAVRANSLR DKYHKPTQII EVLTPKAVVI AATGANRKTV SVDNLKRTP-
VRKFTGDAFS PKWEGPYVIT ETTKAKVQAM SDKVTQKGTV STADMSPTTQ

APPMTLQQWL QWRYNLETTN LLQMNPKMES SLRI-RCWIV FILTVLSILL
TPPMTLPEWM QWRYRQNVNK IREAIPDVQI PLCL-RIWLL FVCVLISVMI
EHVMTLKEWM EWNAHKQLQK LQSTHPELHV PLKM-RMWIL FFLLCFSIVT
APPMTLEQWL LWQAHQALEN VTTLTEEQKQ PTRMDKLWIV LVCVLLLVVF
APPMNLQQWL LWETHLALEN ISSLTEEQKQ PTRMDRVWLF LICVLLIIVF
APPMTLQQWI IWKAHEALQN STTVTDQQKE PTRKDKIWML LVCVLLIVVL
APPMSLQQWI IWQAHQALQN STLVTEEQKE PTRMDRVWML LACILFIIII
APPMTLQQWL VWKANEALKS TTAVSEEEKE PTKVDRVWLF LFCVLLIVVL
APPMSLTEWI LWNMTIMTSN LTGITPDQKK PTIQQRLWIL FCVIVLTVAL
ARPMTLHEWL KWAVRQLTEN LQSLPPEQKE PSWTQKCWII FTLIIASVIL
EEPMTLQQWL RWQLQRKVQQ TPTLTKSETL PLPYTRMWCI GLFCLLLILL
TPVLNLQDWM VW-------- ---------- PTAQVRFWFF FGLIIIGLIL
IPIQHLITLL LWSHRQTSDN IFGLGTEDLH PQTNLRQWSI FVFFFITIIT
---------- --KTKMATNK VTRSAGPKKK PSKVGGCYKF LIALLIMAGL
LPPPQGTGVL IWYQCMWCQN TTSKNAPRQK PPQIPGTW-- ----------

ISVLIAVFRL QWKGAIESPG PILVWNNSRL RRAIHLAQKP VQVNFTSIPQ
IAVIVTVFRM QWKAAIDVPG PVLFWNHTRL RRAVHLADRA VNINITHIPQ
LSTIISILRY QWKEAITHPG PVLSWQVTRR RRDIQYHKLP VEVNISGIPQ
ISCFVTMSRI QWNKDIAVFG PVIDWNVSRS IRVEHATETY VEVNMTSIPQ
VSCFVTVARI QWNRDINVFG PVIDWNVTRS LKVEHPHISY ISINMSSIPQ
VSCFLTISRI QWNRDIQVLG PVIDWNVTRS LRMQHPVPKY IEVNMTSIPQ
VSCFITLSRI QWNKDIQVLG PVIDWNVTRA LRAQHPVPKY VEVNMTSIPQ
VTCFITIARI QWNQDIQVYG PVIDWNISRS IRTHYPIPKN VEVNMTSIPQ
IICFTTAARI QWRHAIITPG PVIDWNSTRD LKVTE--ERY VEVNATGIPQ
VTCFVVMARI QWRNAITVPG IILDWNSTRD LRILE--ENI VEINTTSLPQ
FSLVIVILRL QWRNAIVTPG PIIAWNESRS RRSVEPAPVD IEINITALPQ
GFILSAVFRL QWKNAIHHPG PIISWNLTRE RRAIHPAPRN VHLEICGLQQ
FTIIFSVLRL TWAHTVSLPA TPIHWNLSRP KRALHIEVVP VLVETAGIPF
IALGTGILRI LWAYAETKIQ VPMEDNNSMS RRTLSKGEDN ETLEWHSLPL
---------- -WCHDNSI-- ----WRCTQG RR-------- -------IQL
```

```
GLFLEPHPKP IISKERVLGL SQVVMVDSST LTQKLNLEGE AKSLLIKTIN
GVFLEPFPKP IIDKERVLGI SQIVMIDSGS IAQSMNLDLY MKHLLVDMIN
GLFFAPQPKP IFHKERTLGL SQVILIDSDT ITQGH--IKQ QKAYLVSTIN
GVLYVPHPEP IILKERILGL SQVMMINSEN IANTANLTQE TKVLLADMIN
GVMYTPHPEP IILKERVLGI SQVLMINSEN IANVANLSQE TKVLLTDMIN
GVYYEPHPEP IVVTERVLGL SQVLMINSEN IANNANLTQE VKKLLAEVVN
GVFYQPHPEP IIHTERVLGL SQVLMINSEN VANSANLSQE TKVLLTEMIN
GVYYEPHPEP IIVKERVLGL SQVIMINSET VANSANLTQE AKVLLADMVN
GVILLPHPKP IIQKNRVLGL SQILLINSES LASIFNIKQE HKSILTEIIQ
GILFEPHPKP IIGKERVLGL SQVILINSES IATSLEIKQE HKHILVEMIK
GMLLVPHTKP VVKKERALGF SQIIMSSDS MANSMGLKKE DIHLLVDLLN
GMFWEQFPKP IIHKKRTLGI SQILLIDTPL V---WYIPLK DKKILTQLID
GIIHNPFPKP IVSQRSELLV PFTLNIDTRA LAYCSGLSKD ANTHLAKTIE
GMTTRVVPIP LSLTRGLVQI PVNLVLNSKH LAQ----SAK GRKMLSNNLV
GVIVSNHGDH HTVQHYMLGS GYTVPVSTAT RVQMQKIGPG EWKIATSMVG

EELISLQDVV LNFDLPLGDP HTQEEYIAKR CYQHFGHCYV VKEWPTREII
EEMVALSNVV LPFELPVGDP STQDQYIHKR CYQQFAHCYI VRVWPTSEII
EEMEQLQKTV LPFDLPIKDP LTQKEYIEKR CFQKYGHCYV IKVWPSQDLI
EEMNDLANQM IDFEIPLGDP RDQKQYQHQK CFQEFAHCYL VKGWPSSTVI
EELQDLSNQM IDFELPLGDP RDQDQYIHHK CYQEFAHCYL VSPWISEGII
EEMQSLSDVM IDFEIPLGDP RDQEQYIHRK CYQEFAHCYL VKSWPTEGLI
EEMQSLSDVM IDFEIPLGDP RDQEQYIHRK CYQEFAHCYL VQPWPNEGLI
EELQGLADVM IDFEIPLGDP RDQDQYIHRK CFQEFAHCYL VKGWPSEKLI
EEMRSLQDIT LNFDLPIGNP KTQHEYIQSR CFQEFKDCYL VKPWPTDDVL
EELLSLQNVM LNFDLPLGDP KTQQEYISQR CFQEFKHCYL VKPWPTDDVV
EEMEQLQNII LEFDLPIGDP HDQSYYIEQR CKAALQHCYV VKGWPTDGAI
NEFAQLQEIV LPFTLPLDQP YTQEQYQQKG CFQEFGHCYL VKYWLTSKII
EDLQDLDSRN AHFLVPGTDP WHQTSYADKM CFASYGHCYF VRKWPRPHVY
LLMEQAKLQN PSFEVASGSP EQPSSYLADQ CFAELGHCYL IEYLPDPYIM
LCLDEWEIEC TGFPPPCSLS ITQQQDTVGG SYDSWNGCFV KAPWMRPQPR

QDQCPLNNRL VAFCSPTLYS SWWNYTQSSR EREELFRRKL ETFLNTGCLN
QDQCPLPDRV TIFCSDQLYG NWWYLNRTSE QKEETYRKKM LNLTESSILK
QDQCPLPPKV GIFCSDALYS NWYPRDLPSS VQQSFAQAYI TKVLMQPTLR
ADQCPLPGYI VLFCSDQIYG KWYNIDLTAQ ERENLLVRKL INLAKSSQLK
VDQCPLPRYI VLFCSDQLYG KWYNIENNIQ ENEQLLKTKL YNLTTYSKLK
ADQCPLPGYV VLFCSDQLYS KWYNIENSIE QNEKFLLNKL DNLTTSSLLK
VDQCPLPGYI ILFCSDVLYS KWYNLQNSIL QNENELTKRL SNLTIGNKLK
VDQCPIPGYV VLFCSDRLYN KWYTTENTLL QNEELLIIKL TNLTKDAQLK
ADMCPLPGWV VIFCSDQLYG QWYNASNSKK TNEELLLSKL DKLLNTNKLR
QDMCPLPGWV VLFCSDQLYE KWYIPYGLTE ENDVKLMNKL KTLLNTNKLK
LDQCPLPDYT VIFCSDQLYG SWYYKNDVTQ QRDQTLMLKL RNLTMHAQLK
```

QDHCLIPTSK PSFCSTNMYN S--------- -------GRL LELEL--KLK
ADHCDRPQRV AIFCTSFLYN SWWDESNLSV DGSLELKSIL TSCLATGKLK
ADECLRPSRI AIMCSPLLYG KWKEEQHRCD GREEL----- -DILL---LK
QPKLILPFYI ---------- EFIQKVTTPR Q------DKL TTVLSGNSWK

PEALPGTWHT LGKGEWFRDL TTYDFCKKPE AVFGLNKTYY SWSCLFQPKW
DRALPPTWTP KGQARLFREL NPLDFCTKPE AVMLLNQSYY TWSCLFYPKY
DIAFPKELSP VGSGMLFRPI NPYDICNMPR AVLLLNKTYY TFSCYYRPLY
DRAMPAEWDK QGKADLFRQI NTLDVCNRPE MVFLLNSSYY EFSCLYYPQW
ARALPKEWNN QGNARLFRSF NPLDVCNRPE AVLLLNTTYF TYSCLYYSEY
KRALPKEWSS QGKNALFKEI NVLDVCSKPE LVILLNTSYY SFSCLYYPYQ
NRALPYEWAK GGLNRLFRNI SVLDVCSRPE MVLLLNKTYY TFSCLYYPEY
ERALPPSWST EGKSLLFREA NTLDICNIPE AILLLNTTYY NFSCLYFPKW
SRALPAEWNT QGQNRLFRNL SRIDYCKLPE AVVLLNSTKY DYSCLYYSYY
ARALSAYWHP QGQNKLFRNI TRLDYCKYPE AVILLNTTKS DYSCLYYSEY
DRALPPDWTT QGQNRLFRSI TTFDVCQRPE MVFLLNTTYY TYSCSYFPSY
DCALPSEWK- --QNNLFKSP TITQFCNHPE LIYFLNTTYT TYSCLYYAKQ
PKCLASQWHD NGANEMFIGV TGTSFCDIPR YPIFLNRSES IVSANLMGNY
SSCLSPEWHV NG---YLMEQ NHDTFCSMPR FPAFLNTTLY HVTCEKYPT-
DKIRKQKWQK CYFSGKLRDY EEIDTCPKPL IGPLVTKTLK TGV-------

DSAEIRSDLG YLAYLGAFPS PICIEARNLT DQDYKVTSIY AECVKQGKQY
DQFELANDFG FLAYQKMFPS PICIQNYSLS TEPYKVQSLY QECIQKGTSY
DGTENTEDWG WLAYTDSFPS PICIEEKRIW KKNYTLSSVL AECVNQAMEY
DTPEALYDFG FLAYLNSFPS PICIKNQTIR EPEYKISSLY LECMNASDRH
SSPEAQFDFG FLSYLNAFPG LKYIENQTVR EPEYEVYSLY MECMNSAEKY
DSLESTYDFG FLAYQKNFPA PICIEQQEIR DKDYEVYSLY QECKLASKVH
SNPEALFDFG FLSYMKNFPG PQCIESTVIR QQDYEVYSLY QECKLASKIH
DTAEELYDFG FLAYLGHFPS PICIKEHKIK EVKYSVYSLY QECINKASTY
SSPSFLWDFG WLAYNNHFPS PVCVKETKIR EAKYEVYSLY GECIQATKTY
SSPSYLQDFG WLAYQGHFPS PICEKETKIR MPGYTVYSLF GECLNAAQQH
YGREALNDFG YLAFTDMFPA PTCIETKEVR KPQYKVYSAF QECMIKSQQY
TVSEIKRDWR QLAYSKKFPA PIC----KIV VPKYKVKSLY QKCIAKAKKH
TAKE-RASLG NKRWFNLIQG PLFVNATPFF ADNYAIYSLY QKCKTLSEKY
-------TLG NKRFYGIHKG PLSI----ID RPNYSVYSLY QICKDRVKIY
---------- ---------- ---------- ------IKVI QRLLREYQKT

DIIDVTRQLT SKLVFLGDLP ADRAFSLNNW RRLQITGQSM NQAITTLSKL
DLEDVINQLL RVLIDLGKVP ASRAFTPDNF RKLQTSGLSM NQAISTLAKI
GIDEVLSKLD LIFGNLTHQS ADEAFIPENL RRIQEAGLGL ANAITTVAKI
GIDSALLALK TFLQSVNEMP LARAFVGNNY ERLRSMGYAL TGAVQTLSQI
GIDSVLFALK TFLTPVNEMS TARAFVGTNI EKLRSMGYSL TGAVQTLSQI
GIDTVLFSLK NFLRPVNEMP NARAFVGNNY AKLKSMGYAL TGAVQTLSQI
GIDSVLFSLK NFLKPVNEMP NARAFVGNNY SKLRSMGYAL TGAVQTLAQI

GIGNVVEGIK ELLTPVNEMP NARAFVGNNY VKLRSMGYSL TGAVQTLSKI
TIDQVLVGLH GFLTPVQDLP KERAFLGTNW QKLQKAGYAI TNAVTQIAKI
GIERALIGLH AFMTPLQEVN KERAFIGHNW GKLQAVGFTI TNTVSKIARI
DINDVIAKLE ALFTPLQGRP QNRAFM-TNF HKIQSIGFNL ANAISTVSKI
HVESVQLLNE LFIINIKELP TEDKRWGPNT QRLEKVSLMM ANSTATVSKL
SLFSVLQALE EFIASLLQMN PKRAIWGNNA QIFRKTNFLL AKSMEKISRL
PLTDVLSGLK KL-VYVLDMV PTKAIWGTND QYLTQSATLT NKAINLLSQT
GVTFNLPQVQ SLPNPGHHIP KSR----NRW KDLQIAGLGV QQKLMGLTRE

SDLNDENLAA GIHLLQDHIV TLMEATLHDV SLLGHMTSIQ HLHTHLATFK
SDLNDENLAA GIHLLQEHIV TLMEATVHDI SMLEAAHGLQ ILHTHLSTLR
SDLNDQKLAK GVHLLRDHVV TLMEANLDDI VSLGEGIQIE HIHNHLTSLK
SDINDERLQH GVYLLRDHVV TLMEAALHDV SIMEGMLAIQ HVHTHLNHLK
SDINDERLQQ GVSLLRDHVV TLMEAALHDI TIMEGMLAIQ HVHTHLNHLK
SDINDENLQQ GIYLLRDHVI TLMEATLHDI SVMEGMFAVQ HLHTHLNHLK
SDINDQNLQQ GIYLLRDHIV TLMEATLHDI SIMEGMFAVQ HVHTHLNHLR
SDINDENLQQ GLYLLRDHLV TLMEATLHDI SLMEGMLAVQ HLHTHLNHFK
TDLNNEAIVS GIYLLKDHIV TLMEATLHDV SALGNVVTIQ HFHTHLAQFK
IDLNNEHLVS GLYLLKDHLV TLMESTLHDI SILGNAVAIQ HFHTHLTQLK
SDLNDNQLAK GMHVLRNHLV TLMEATLHDI SKFESGLALQ HLHTHLAQLR
SDLNEYLFAD GLHILKDHVV TLLEANMKDT QHIDELTTAM LILSYIQNFR
QDANNINLRN GVYLVKDALT QVALIVKHDL AVLSDELIME IIVTQLQKII
MDDNFAQLEQ GFKVLQSSLM HTLRVLTKDM ATLGDAIKVK QVEAQIQRGL
ATFEAWNALK GISTLRQLLL QIKGTNTTLC SAMGPLMA-- ---TNIQQIM

NLLIGNRVDW SVLYYELIHL THNSGYLTHV TIHHPYEIVN QDCEELTFLH
LLLTENRVDW NLIYYELIHL IKNAGHLARV ELQHPYEIVN QDCEQLTYLE
LLTLENRIDW RFIYYEIIHL VRNAGYLSKV WIQQPFEVLN QECGTNIYLH
TMLLMRKIDW TFIYYEIVHL LESAGHLTHV KVKHPYEIIN KECSDTQYLH
TILLMRKIDW TFIYYEITHL VESAGHLTLI RVKHPYEVIN KECTYEQYLH
TMLLERRIDW TYMYYELTHL VQSAGQLTHV TIAHPYEIIN KECTETKYLH
TMLMERRIDW TYMYYELIHL IKSAGQLTHV TVSHPYEIIN RECSNTLYLH
TMLLERRIDW TFIYYEIIHL IHSAGQLTHV TIDHPYEILN RECEETKYLH
LLLVENRIDW NYIYYEIIHL VDSAGSLTLV TIQHPYTIVN QECGETKYLH
LLLMENRMDW TFIYYEIIHL VYSADNLVQI NVEQPYEILN VECGKSTYLH
STLQENRVDW SILYFEIIHL VFNSGQLTQV FVEQPYNLVS MECNIPTYLH
IPSTEGRIDW RILYYQIIHL VKTGTQLTLT KIHQPYTHIS -ECSELYYLE
FSLSNGHVPW TIGYFQILHV LRNMSPQMFL ETQHTYILTN VKILDICFLH
ALLENNKVPW ALLQMQLPFL VKTNNLLALA KVRPPYQYIA -KCGNINYLK
FALQHGNLP- ---MLSLPRA VA----VYDM GVRH------ ---NRTLYVD

LVDCHEQDYL ICEEVMEVEP CGSDCPVLAE NIQAPYVYLH PLKNGSYLLM
LKGCQELDYL VCEEILQHEP CGSDCPVTAQ KIKDPYVWIY PLKNGSYLIM
MEECVDQDYI ICEEVMELPP CGSDCPVLTK PLTDEYLEIE PLKNGSYLVL

LEECIREDYV ICDIVQIVQP CGSDCPVTAL KVKTPYIQVS PLKNGSYLVL
LEDCISQDYV ICDTVQIVSP CGSDCPVTAE KVKEPYVQVS ALKNGSYLVL
LKDCRRQDYV ICDVVEIVQP CGSDCPVWAE AVKEPFVQVN PLKNGSYLVL
LEECRRLDYV ICDVVKIVQP CGSDCPVWAE PVKEPHVQIS PLKSGSYLVL
LEQCIKQDYV ICDIVERVQP CGTDCAVYAK AIKSPYTEIL PLKNGSYLVL
MEECTEQDYK ICEQVTEVLP CGSDCPVLAK TVKPGYVHIE SLRNGSYIYM
IDKCEEQDYV ICEVIQEKQP CGSDCPVKAR TIEKGYTYIQ PLKNGSYVVM
IEECVNQDYL ICDIVEEVLP CGSDCPVMAK AVKAPFVSIT PLKNGSYVIL
PKGCEQRDYL ICEEINLHQT CGSKCPVTGK AV--PYLEFI PLKNGSYVVM
IS-------- ---------P CGTSCPIRIQ SSNISFVFID SLTNGSYLIL
VKECYLRGFR YCDNWEKQLP CGQSCPITVK PI--TFAQVT SLGNGSYIIM
LSKCVETGTI YCDRKFTKGP SETWC----- ---------- -LNSGTWSSL

ASHTDCSLPP YEPVVVTVND SLECYGKPLK RPLVGIIAKL KSLKIKVTST
SSHTDCAIPP YEPVLVTVND TVRCFGTTLK KPLVGLIAKI KGLKIEITST
SSTTDCGIPA YVPVVITVND TISCFDKEFK RPLTSLIAKI KGIQIEITSS
SSTKDCSIPA YVPSVVTVNE TVKCFGVEFH KPLTGIIASL QSLEIEVTST
TSRTDCSIPA YVPSIVTVNE TVKCFGVEFH KPLVGIIANL QNLEIEVTST
ASSTDCQIPP YVPSIVTVNE TTSCYGLNFK KPLVGIIAKI KGLKIEVTSS
ASSTDCQIPP YVPSVVTVNE TTQCFGVTFK KPLVGIIAKI KGIKIEVTSS
SDSTSCNILP YIPSIVTVNE TVECFGVLFK KPLLGIIAKL KNIKIEVTST
AHYQDCGIKP YVPQIVTVNA TVKCLGYEIQ PPLVGILAKL KNIQIQVTST
SHFQDCHIKP YIPQIVTVNA TVKCLGEVFQ PPLVGIITKL KGFQVQITST
ADTSACTIPA YSPVLVTTND TLQCYGHILK RPLVGVIAQL KDLKFQVTSS
SYTIDCNIPP YQSSIFTIND TVTCFEKILK KHLVGILAKL KKIEVKATDT
AGKSECNIPA LQPSIVTVNT TITCYGRQLF PPPTGIIAKL QRSEITLFNT
VSYSECDIPS MRTVIVTPST RMTCYNMTLF -PLVAVIARL QDYEITLSYT
KNETNCSFSL TRPVCFHLNS TAQWRGHVL- -PFVGNSQEA PNTEIWEGLI

WESIKDQIHR SEQELLRLDL HEGDYSDWIL QLGNALEDVW PVAASAVSTI
WENIKDQIKR SEAELLRLDL HEGDYAEWTK QLGKALEDIW PAAAQTVSKI
WETIKEQVAR AKAELLRLDL HEGDYPEWLQ LLGEATKDVW PTISNFVSGI
QENIKDQIER AKAQLLRLDI HEGDFPDWLK QVASATRDVW PAAASFIQGV
QESIKDQIER AKSQLLRLDI HEGDFPAWIQ QLASATRDVW PAAARALQGI
GESIKDQIER AKAELLRLDI HEGDTPAWIQ QLAAATKDVW PAAASALQGI
GESIKDQLER AKAELLRLDI HEGDTPAWIR QLAAATEDVW PAAASALKGI
QENIKDQIER AKAELLRLDI HEGDSPAWIK QLAAATEDVW PTLATGLKSI
WESIKDQVEK SQTELLRLDI HEGDTPAWIK QLAESTKDIW PTTANIFGKV
WESIKGQVEQ AQAELLRLDL HEGDSGQWIK QLASASKDIW PAAATVLGKI
WESIKDQIAR SKELLLQLDL HEGSAPEWIN RLAAAAADIW PATGQALKGL
WASIEEQIED TKSDLLRLEL HKGDTPEWIK QLGEALEDVW PAAASATKTI
HDATEDILQE VKQLLQRIDI HEGDFPLWLN RLASAVSAAW PSLANMANSI
HDNITQLLAR SRQLLSMVRF EAAEIPQWLQ LIGEGLIELI PQVKSAIEWG
EEAIREH-NK VQDILTKL-- -EQQHQNWKQ NTDNALQNMK DAIDSMDNNM

GTLLEKAAGT LFGNVFSILA YAKPVIIGII LIILLLLVIR ILR
GDFLGKIADG IFGTTFSLLT YAKPVIIGIV VIVLLILIIR ILS
GNFIKDTAGG IFGTAFSFLG YVKPVLLGFV IIFCIILIIK IIG
GNFLSNTAQG IFGSAVSLLF YAKPILIGIG VILLIALLFK IIS
GNVLSNTAQG IFGTTVSILS YAKPILIGIG VILLIAFLFK IVS
GNFLSGAAHG IFGTAFSLLG YLKPILIGVG VILLIILIFK IVS
GNFLTGAAQG LFGTAFSILG YLKPILIGIG IIILVILIFK ILR
GNFLSDAAQG IFGTAFGILG YVKPILIGVG IILLIVVIFK IIS
GEFLSGTFGG LFGT----LG YIKPIILGIV ILLLIVIVVK IIS
GDFLGGTAGS IFGI----FG YLKPIFIGLT ILILIVLVFK ILS
GDFLQSTVGS LLGTGLSFLS YLKPILIGIG LIFLVVILFK IIS
ASFVSSATKG IFGGIIDILT YTKPIVILII ITILIVLIFR ILK
AHADTSIGTS ILGTGLQILT YLKPIFIAIV LIVLLIIVIK IFR
IDLTAKTITQ TVTSVLQSLN FLVPVLIIAG VVFVLILVLK FFS
LTFYEYTQYG LF--IVCLLA FLFAVIFGCG VTVRLREVFT ILS
